# Promiscuous enzymes cooperate at the substrate level en route to lactazole A

**DOI:** 10.1101/2020.05.12.092031

**Authors:** Alexander A. Vinogradov, Morito Shimomura, Naokazu Kano, Yuki Goto, Hiroyasu Onaka, Hiroaki Suga

## Abstract

Enzymes involved in ribosomally synthesized and post-translationally modified peptide (RiPP) biosynthesis often have relaxed specificity profiles and are able to modify diverse substrates. When several such enzymes act together during precursor peptide maturation, a multitude of products can form, and yet usually, the biosynthesis converges on a single natural product. For the most part, the mechanisms controlling the integrity of RiPP assembly remain elusive. Here, we investigate biosynthesis of lactazole A, a model thiopeptide produced by five promiscuous enzymes from a ribosomal precursor peptide. Using our *in vitro* thiopeptide production (FIT-Laz) system, we determine the order of biosynthetic events at the individual modification level, and supplement this study with substrate scope analysis for participating enzymes. Combined, our results reveal a dynamic thiopeptide assembly process with multiple points of kinetic control, intertwined enzymatic action, and the overall substrate-level cooperation between the enzymes. This work advances our understanding of RiPP biosynthesis processes and facilitates thiopeptide bioengineering.

## Main text

Ribosomally synthesized and post-translationally modified peptides (**RiPPs**) are structurally and functionally diverse natural products united by a common biosynthetic logic.^1^ Usually during RiPP maturation, biosynthetic enzymes utilize the N-terminal sequence of a ribosomally produced precursor peptide as a recognition motif (leader peptide; **LP**) and install post-translational modifications (**PTMs**) in the C-terminal section of the same substrate (core peptide; **CP**). This mode of action leads to relaxed substrate requirements around the modification sites, which is often exemplified by one RiPP enzyme introducing multiple PTMs in a single substrate. In one extreme case, a single enzyme epimerizes 18 out of 49 amino acids in polytheonamide A precursor peptide during its biosynthesis.^2,3^ Unique enzymology of RiPP biosynthetic enzymes has come under intense scrutiny in the recent years, which explained observed substrate specificities in many cases.^4,5,14,15,6–13^ During biosynthesis of complex RiPPs, when multiple enzymes capable of differentially modifying their substrate act together, a multitude of products can often form, and yet usually the biosynthetic pathway manages to produce a single natural product. Molecular mechanisms controlling the integrity of RiPP biosynthesis are only beginning to be elucidated,^2,16–23^ and many details remain unclear, especially in the cases where enzymes can apparently compete over the substrate. For example, during biosynthesis of some thiopeptides, Ser and Thr residues in the precursor peptide CP are selectively modified to either oxazoline/oxazole or dehydroamino acids by cyclodehydratase/dehydrogenase and dehydratase enzymes, and the basis for such a cooperative action despite the potential for competition has not yet been firmly established.

In this study, we aim at investigating the roots of cooperative biosynthesis of lactazole A, a cryptic thiopeptide from *Streptomyces lactacystinaeus* (Fig. 1).^24^ Lactazole biosynthesis involves 5 dedicated enzymes colocalized with the precursor peptide gene (*lazA*) into *laz* biosynthetic gene cluster (**BGC**; Fig. 1a–c). During thiopeptide maturation, LazD and LazE operate as a single cyclodehydratase enzyme,^25–27^ responsible for the installation of azoline PTMs (Fig. 1d), which are further dehydrogenated to azoles by the C-terminal domain of LazF in an FMN-dependent manner.^9,28^ Dehydroalanines (**Dha**) are accessed from Ser by the combined action of LazB and the N-terminal domain of LazF, which utilize glutamyl-tRNA^Glu^ for dehydration similarly to class I lanthipeptide synthetases (Fig. 1e).^29–32^. The remaining enzyme, LazC, performs macrocyclization and eliminates LP as a C-terminal amide (**LP-NH_2_**) to yield the thiopeptide (Fig. 1f).^33,34^

**Figure 1.**
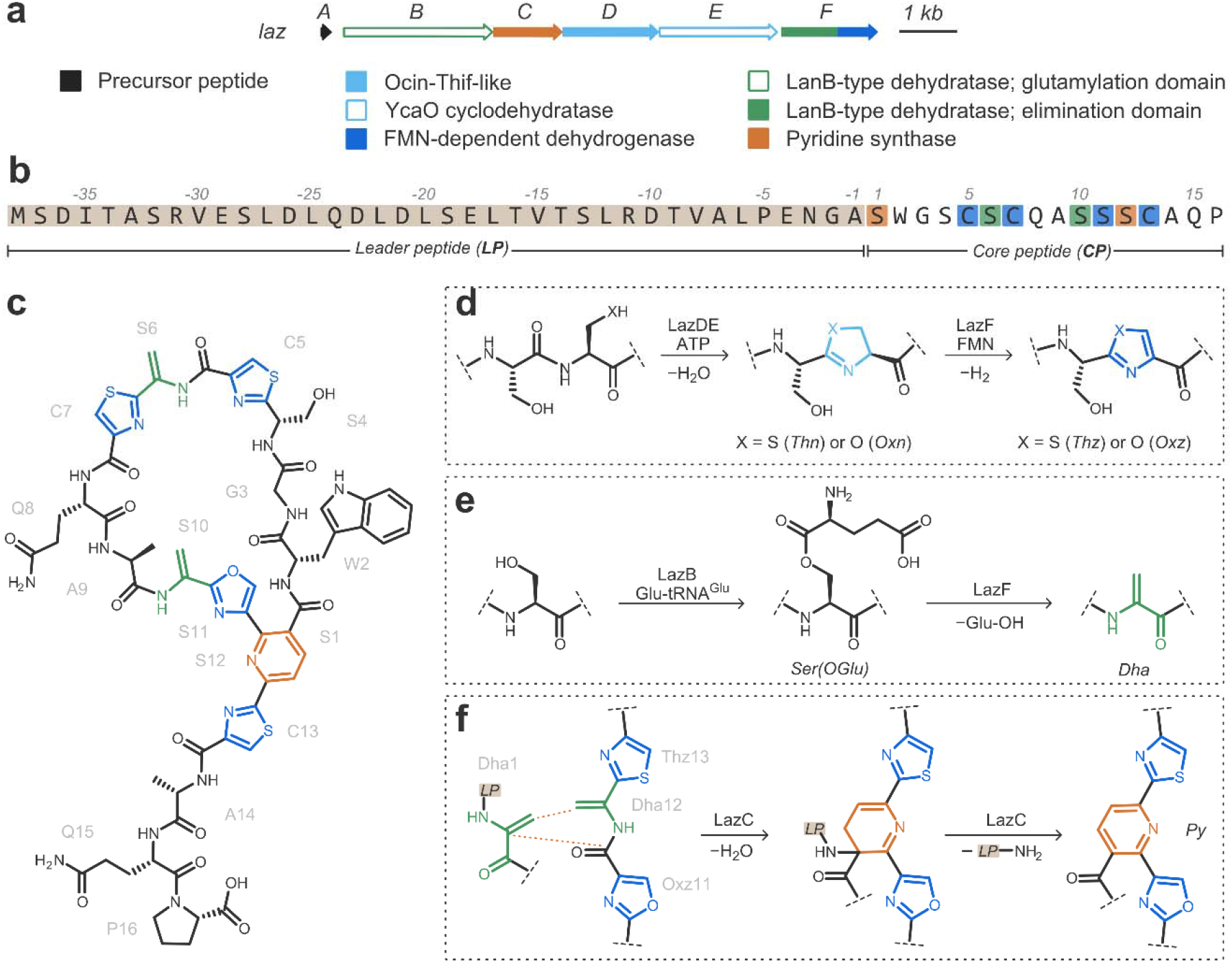
Biosynthesis of lactazole A. a) The biosynthetic gene cluster from *Streptomyces lactacystinaeus* responsible for lactazole A production (*laz* BGC; GenBank accession: AB820694.1; MIBIG accession: BGC0000606). Genes encoding the enzymes responsible for synthesis of azolines are color coded in light blue, azoline dehydrogenase in blue, dehydroalanine — green, and pyridine — orange. b) Primary amino acid sequence of LazA precursor peptide. Residues eventually converted to Dha, azoles and pyridine are highlighted in green, blue, and orange, respectively. c) Chemical structure of lactazole A. d) LazDE performs a cyclodehydration reaction furnishing azoline heterocycles, which are further dehydrogenated by FMN-dependent LazF. e) Ser dehydration is catalyzed by LazBF via a two-step process involving Ser glutamylation by LazB. f) Macrocyclization is achieved by LazC, which utilizes two Dha residues, Dha1 and Dha12, to form the central dihydropyridine and concomitantly macrocyclize the peptide. The same enzyme consequently eliminates LP-NH_2_ to aromatize the structure.

Lactazole A is an example of a RiPP assembled by multiple enzymes that can compete over the substrate. LazA CP contains 6 Ser residues, 4 of which are converted into Dha, 1 into oxazole (Oxz), and 1 which remains unmodified (Fig. 1b, c). Previously, we reconstituted biosynthesis of lactazole A *in vitro* by combining flexible in vitro translation (**FIT**)^35^ — utilized to access LazA precursor peptide — with recombinantly produced Laz enzymes.^36^ Using this platform, termed the FIT-Laz system, we showed that selective biosynthesis of lactazole A occurred only when all enzymes were present in the reaction mixture from the beginning, whereas stepwise treatments led to either under- or overdehydrated thiopeptides containing 3 or 5 Dha, respectively. These results suggest that cooperation between Laz enzymes extends beyond the “azoles form first, Dha second” model observed for thiopeptides studied to date,^20,21,37^ and indicate that lactazole may be a good model system to study concerted action of multiple enzymes in RiPP biosynthesis. Furthermore, our previous findings^36^ indicated that Laz enzymes can convert extensively mutated LazA analogs to corresponding thiopeptides, exemplified by the synthesis of over 90 lactazole-like thiopeptide containing up to 25 nonnative amino acids. How the enzymes maintain the integrity of biosynthesis for a diverse set of substrates constitutes the second major question of this study.

According to our aims, we intercepted biosynthetic intermediates and reconstructed the order of events leading up to the final macrocyclization reaction during the lactazole A assembly process at a single PTM resolution. When supplemented with substrate preference studies for individual enzymes, these results help in rationalizing the roots of cooperation between Laz enzymes, and establish the basis for selective lactazole A production. Bioengineering of RiPPs to harness their potential for human health holds a lot of promise,^38,39^ but it requires thorough understanding of the underlying biosynthesis mechanisms. Our study elucidates how several promiscuous enzymes coordinate the assembly of a complex RiPP, facilitating thiopeptide bioengineering, and ultimately, functional reprogramming.

## Results

### Biosynthetic timing

Because our previous results indicated that modification of amino acid residues 4–7 in the CP of LazA (Fig. 1b) is not essential for macrocyclization,^36^ we hypothesized that maturation of LazA is modular, i.e. residues 10–13 undergo PTM independent of residues 4–7. Accordingly, we sought to establish the order of PTM installation for two LazA mutants, LazA S4-C7A (**LazA^min^**; Fig. 2a) and LazA S10-C13A (**LazA^aux^**; Fig. 2b), before proceeding to the wild type peptide (**LazA^wt^**).

**Figure 2.**
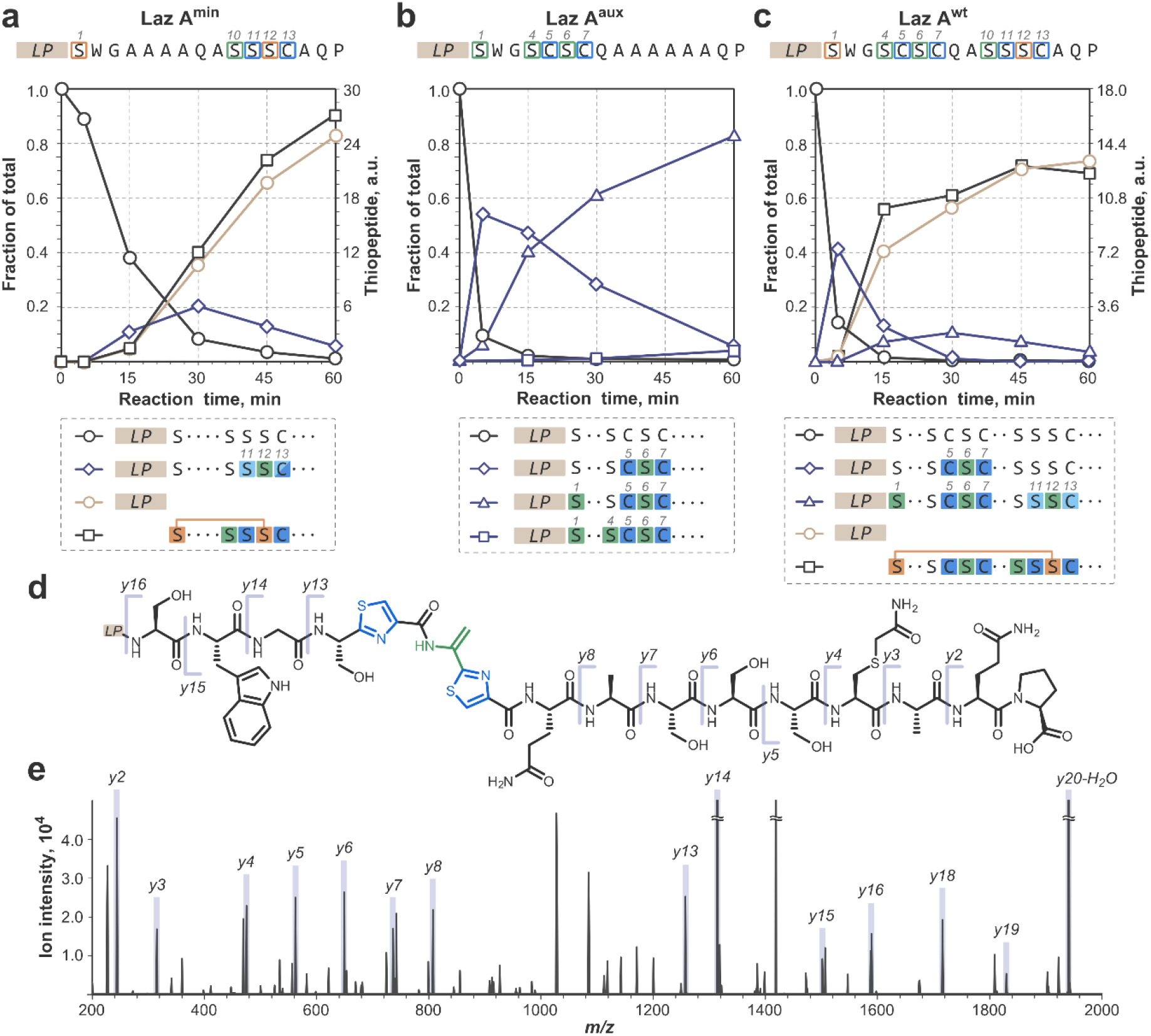
Time course analysis reveals the order of enzymatic action during lactazole A assembly. a) LazA^min^ maturation time course. In vitro translated precursor peptide was incubated with LazBCDEF/GluRS/tRNA^Glu^ for 5, 15, 30, 45 and 60 min, after which reaction outcomes were analyzed by LC-MS and DDA MS/MS. Displayed are the changes in the amounts of starting material (S.M.), LP-NH_2_, thiopeptide (lactazole S4-C7A) and the key intermediate, Ser10-Oxn11-Dha12-Thn/Thz13, as a function of time. For full product distribution and characterization of intermediates refer to S.I. 3.3. b) LazA^aux^ maturation time course; data as panel in a), except LazC was omitted from the enzyme mixture, and the 45-min time point was skipped. Displayed are the changes in the amounts of S.M. and three key products as a function of time. See also S.I. 3.4. c) Lactazole A biosynthesis time course; data as panel in a). Maturation of LazA^wt^ proceeds in a modular fashion: after a 5-min treatment with Laz enzymes, the key Ser4-Thz5-Dha6-Thz7 intermediate accumulates similarly to the LazA^aux^ case (compare to panel b). Fast accumulation of this peptide points to a modular biosynthetic logic. The second key intermediate, Thz5-Dha6-Thz7/Oxn11-Dha12-Thn13, accumulates analogously to the LazA^min^ case (compare to panel a). See also S.I. 3.5. d) The chemical structure of the key Ser4-Thz5-Dha6-Thz7 LazA^wt^ intermediate observed after a 5-min treatment with the enzymes. e) MS/MS spectrum supporting structural assignment of the intermediate from panel d). See also Fig. S37.

First, we investigated *in vitro* modification of LazA^min^ utilizing the FIT-Laz system.^36^ Synthetic DNA bearing *lazA^min^* gene was *in vitro* transcribed and translated, generating the precursor peptide, which was incubated with a mixture of recombinantly produced enzymes (LazB, LazC, LazD, LazE, LazF, *Streptomyces lividans* GluRS and synthetic *S. lactacystinaeus* tRNA^Glu^ (**LazBCDEF/GluRS/tRNA^Glu^**); S.I. 2.3) at 25 °C. Reactions were stopped by the addition of cold methanol containing iodoacetamide (**IAA**), and the outcomes were analyzed by LC-MS. Selective IAA alkylation on unmodified Cys residues and quantitative acidic hydrolysis of Oxn under HPLC conditions enabled unambiguous identification of PTMs based on mass shifts (S.I. 3.1), but not their location within the CP. Accordingly, the captured intermediates were further analyzed by CID tandem mass spectrometry (**MS/MS**) in a data-dependent acquisition (**DDA**) mode (S.I. 2.4-2.5). Our initial experiments showed that LazA^min^ maturation is complete in under 3 h (Fig. S6). To intercept biosynthetic intermediates at a finer temporal resolution, we performed a time course study and quenched the reactions after 5, 15, 30, 45 and 60 min (Fig. 2a and S7). These experiments revealed formation and consumption of multiple linear intermediates during thiopeptide assembly, and enabled assignment of the PTM installation order (Fig. 4a, S.I. 3.3).

Maturation of LazA^min^ started with formation of oxazoline at Ser11 (Oxn11), followed by heterocyclization at Cys13 (Thn13; Fig. S10). After that, Ser12 was dehydrated to Dha12 (Fig. S11), and Thn13 was dehydrogenated to give thiazole13 (Thz13; Fig. S15), arriving at a prominent intermediate, Ser10-Oxn11-Dha12-Thz13 (Fig. 2a). The following steps, Oxn11 dehydrogenation and Dha10 formation, happened fast relative to the temporal resolution of our experiments, and we were unable to capture the corresponding intermediates even when using diluted enzyme mixtures or performing reactions at 4°C (Fig. S16). The next observed peptide bore the Dha10-Oxz11-Dha12-Thz13 motif required for macrocyclization (Fig. S12). Dehydration of Ser1 to Dha1 was independent of other modifications, because most intermediates formed as a mixture of Ser1 and Dha1 throughout the time course. Treatment of LazA^min^ with LazBF/GluRS/tRNA^Glu^ for 2 h confirmed that LazDE-independent dehydration of Ser1 is kinetically competent (Fig. S17). Additionally, the time course study indicated that LazBF dehydrates Ser adjacent to azolines. We confirmed the azoline-dependent activity of LazB by treating LazA^min^ with the enzyme mix lacking dehydrogenase (LazBDE/GluRS/tRNA^Glu^; Fig. S18).

The order of Ser10 dehydration and Oxn11 dehydrogenation could not be determined from the time course. LazF can convert Oxn11 to Oxz11, but this process is kinetically incompetent. Although a 17 h treatment of LazA^min^ with LazDEF yielded the Oxz11/Thz13 product, an analogous 2 h reaction mainly resulted in the formation of Oxn11/Thz13 (Fig. 3a). In contrast, a 2 h incubation of LazA^min^ with LazBDEF/GluRS/tRNA^Glu^ afforded fully modified Dha10-Oxz11-Dha12-Thz13, suggesting that Dha formation is required for facile oxidation of Oxn11. To test which Dha is important, we prepared two LazA^min^ mutants, S10A and S12A, and treated them with LazBDEF/GluRS/tRNA^Glu^ for 2 h. The S10A mutant yielded the Ala10-Oxn11-Dha12-Thz13 peptide (Fig. 3b), and modification of LazA^min^ S12A led to a complex mixture of products, none of which bore Dha10 and/or Oxz11 (Fig. 3c). These results suggest that installation of Dha12 is required for dehydration of Ser10, and in turn, Dha10 greatly facilitates dehydrogenation of Oxn11. Thus, during LazA^min^ maturation, the key intermediate Ser10-Oxn11-Dha12-Thz13 is converted to Dha10-Oxn11-Dha12-Thz13, which sets the stage for Oxn11 oxidation and the follow-up macrocyclization (Fig. 4a). The elusive Dha10-Oxn11-Dha12-Thz13 intermediate could be captured for LazA^min^ S11T treated with LazBCDEF/GluRS/tRNA^Glu^ for 2 h (Fig. S19).

**Figure 3.**
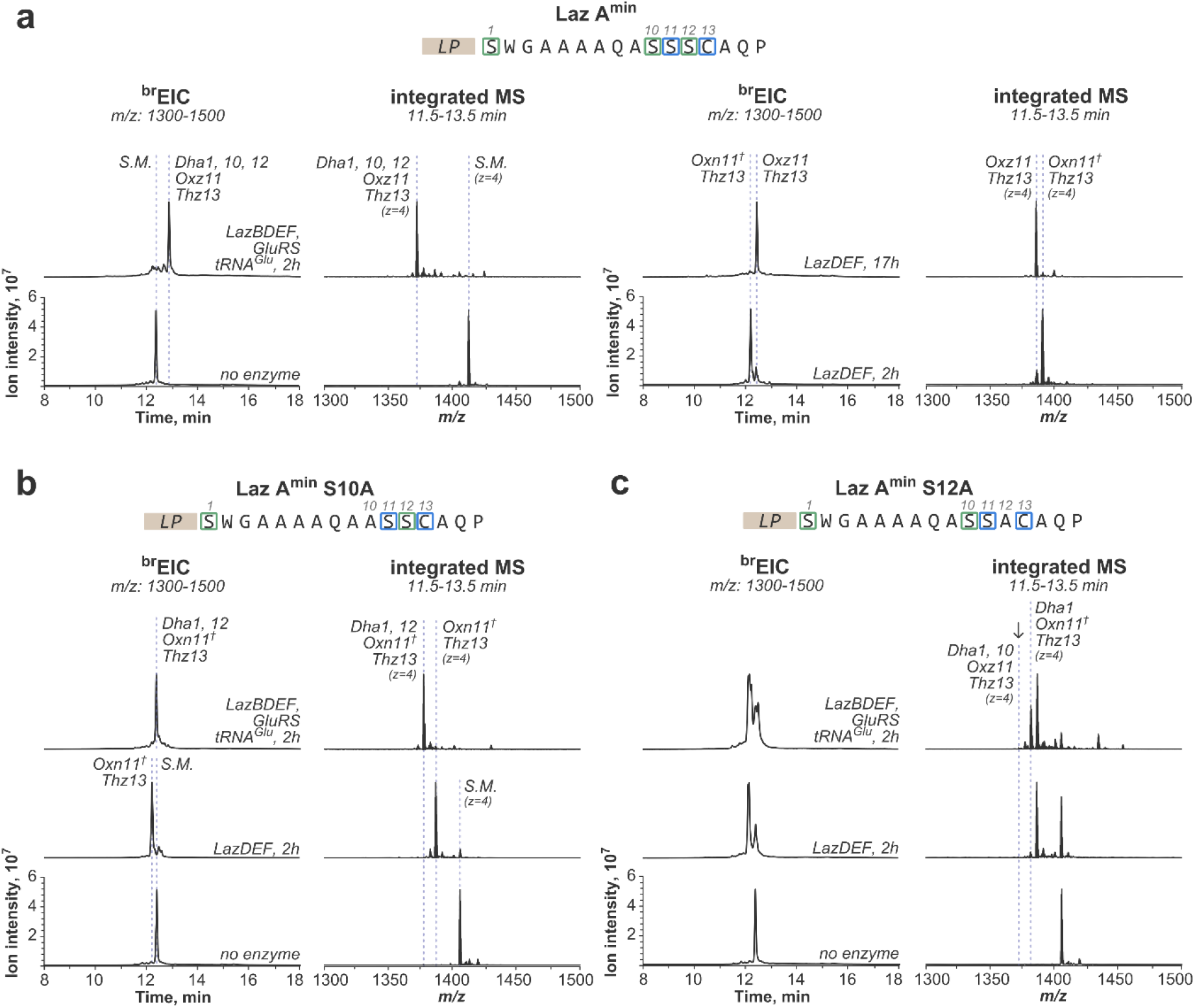
Both Dha10 and Dha12 are required for facile Oxn11 dehydrogenation. a) Dha-dependent Oxn11 dehydrogenation in LazA^min^. In vitro translated LazA^min^ was incubated with LazBDEF/GluRS/tRNA^Glu^ (2 h), LazDEF (2 h or 17 h) or with buffer only, and the outcomes were evaluated by LC-MS. Displayed are ^br^EIC chromatograms and integrated mass spectra showing the product distribution. These data suggest that in the absence of LazB, LazF-mediated dehydrogenation of Oxn11 is kinetically incompetent. b) Kinetically incompetent Oxn11 dehydrogenation in LazA^min^ S10A. Analogous to panel a), data for LazA^min^ S10A. In the absence of Dha10, Oxn11 oxidation does not happen on a relevant time scale. c) Kinetically incompetent Oxn11 dehydrogenation in LazA^min^ S12A. Analogous to panel a), data for LazA^min^ S12A. The S12A mutation disrupts azoline installation by LazDE and leads to formation of multiple products. Nevertheless, when the peptide was treated with LazBDEF/GluRS/tRNA^Glu^ for 2 h, neither Dha10 formation nor Oxn11 dehydrogenation took place, suggesting that Dha12 is required for dehydration of Ser10.†: Oxn underwent quantitative hydrolysis during HPLC (see S.I. 3.1 and 3.2).

**Figure 4.**
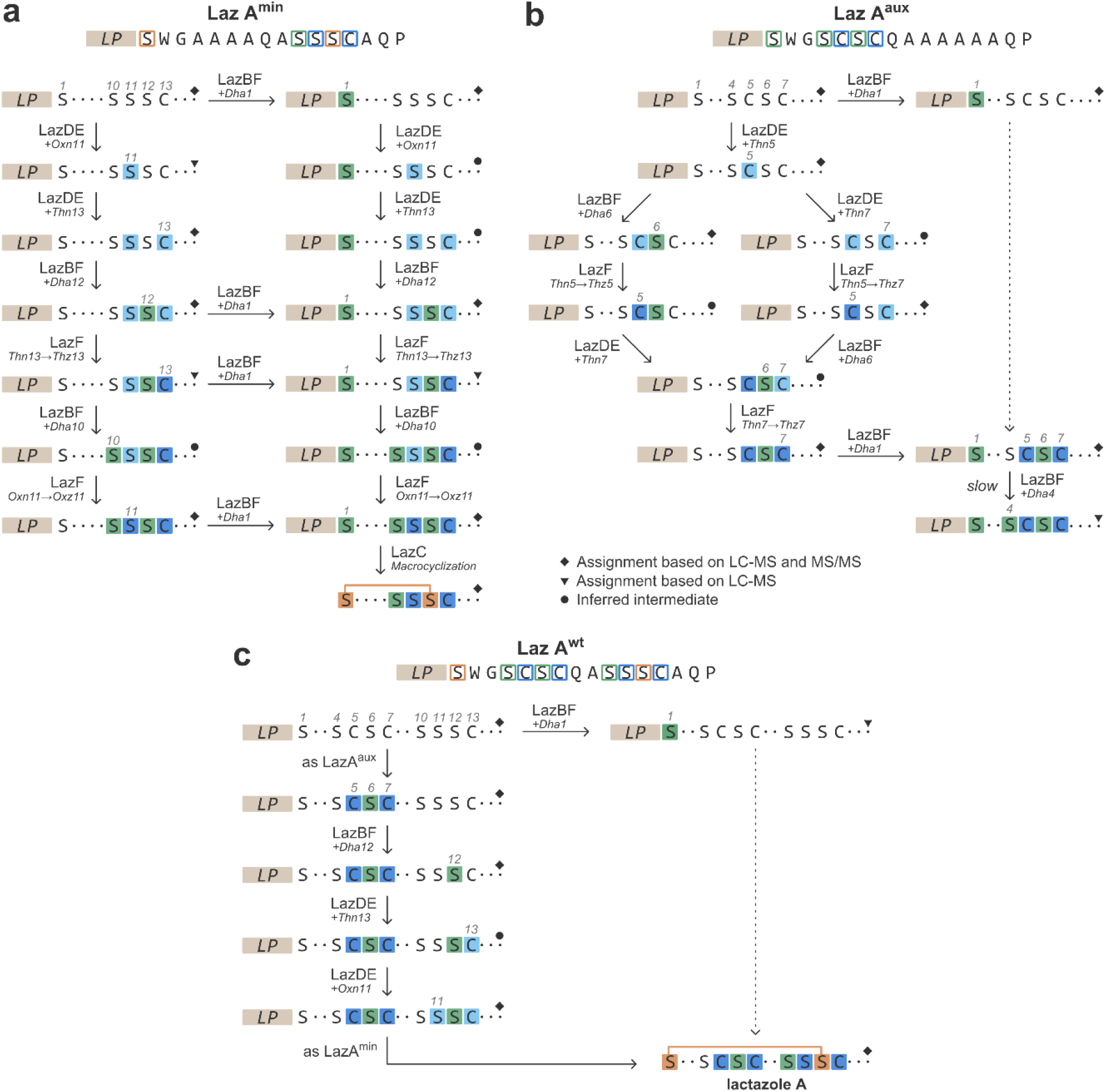
Proposed lactazole A biosynthesis pathways. a) The order of PTM installation for LazA^min^. b) Modification of residues 4–7 as studied in LazA^aux^. c) Modular assembly of lactazole A from LazA^wt^. See also S.I. 3.3 – 3.5.

To test whether this curious sequence of events has biosynthetic significance, we forced oxidation of Oxn11 prior to adding LazB to the reaction mixture. A 17 h treatment of LazA^min^ with LazDEF furnished the Oxz11/Thz13 product, which upon further incubation with LazBCF/GluRS/tRNA^Glu^ for 2 h accumulated the Ser10-Oxz11-Dha12-Thz13 intermediate, and a mixture of thiopeptides bearing either Ser10 or Dha10 (Fig. S20). Similarly, LazA^min^ S11C, readily modified by LazDEF to the Thz11/Thz13 intermediate, underwent slow dehydration at Ser10 and formed a mixture of thiopeptides (Fig. S21). These results indicate that premature oxidation of Oxn11 hampers Dha10 formation, and highlight the importance of Dha-dependent Oxn11 dehydrogenation during the biosynthesis. Lastly, we found that the order of modifications described here persisted for three LazA^min^ variants, although maturation speed and efficiency varied between the mutants (Fig. S22). The Ser10-Ser11-Ser12-Cys13 motif is conserved in over 60 thiopeptides from predicted lactazole-like BGCs (Fig. S23),^40^ which lends further support to our findings.

Next, we studied modification of LazA^aux^. Analogous to the experiments above, LazA^aux^ precursor peptide was incubated with LazBDEF/GluRS/tRNA^Glu^ for 5, 15, 30 and 60 min, and reaction outcomes were analyzed by LC-MS and DDA MS/MS (Fig. 2b and S.I. 3.4). Heterocyclization at Cys5 initiated the chain of PTMs, which then briefly bifurcated and converged on the key product, Ser4-Thz5-Dha6-Thz7, bearing the wild type modification pattern (Fig. 4b). This sequence of events was fast — the key intermediate comprised over 50% of total substrate after only 5 min. As before, dehydration of Ser1 to Dha1 was independent of other PTMs, and was generally slow compared to the modifications at residues 5–7, taking over 1 h to completion. Even slower was dehydration of Ser4 to Dha4 inside the Ser4-Thz5-Dha6-Thz7 motif, as this product comprised less than 4% of total substrate after 1 h. These results confirm that Ser inside the Ser-azole-Dha-azole pattern is a poor substrate for LazBF.

Finally, we performed time course analysis for LazA^wt^ (Fig. 2c; S.I. 3.5). As anticipated, biosynthesis proceeded largely in a modular fashion (Fig. 4c). The key intermediate of LazA^aux^ maturation, Ser4-Thz5-Dha6-Thz7, also quickly formed for LazA^wt^ and comprised over 40% of total substrate after a 5-min incubation with the full enzyme set (Fig. 2c–e). The biosynthetic order leading up to this peptide was identical to LazA^aux^, including the bifurcation event. Modification of residues 10–13 proceeded once the Ser4-Thz5-Dha6-Thz7 motif was installed. With the exception of Ser12 dehydration, which took place prior to the heterocyclization of either residue around it (Fig. S38), modification of residues 10– 13 generally followed the sequence established for LazA^min^, including the Dha-dependent Oxn11 dehydrogenation.

Altogether, these experiments unravel lactazole A biosynthesis one PTM at a time, revealing a carefully orchestrated sequence of events, in which the actions of different enzymes are intertwined to ensure the integrity of biosynthesis.

### Substrate specificity

Primary macrocycle assembly for thiopeptides studied to date separates azole and Dha formation into two stages, where Dha installation follows azole formation.^20,21,37^ Such a separation can be due to the specificity of a Dha-installing enzyme toward the native azole pattern,^20^ or due to an additional enzyme acting as a gatekeeper and preventing premature Dha synthesis.^21,37^ Intertwined enzymatic action observed in the experiments above is distinct from these models, indicating that lactazole biosynthesis integrity is maintained differently. The time course study revealed some of these mechanisms (for instance, Dha-dependent dehydrogenation at Oxn11 serving to prevent formation of underdehydrated thiopeptides), but some issues, especially the nature of enzyme competition over Ser residues, remained elusive. To gain deeper insight into the nature of cooperation and competition during lactazole biosynthesis, we performed analysis of individual enzymes and their innate substrate preferences. To this end, we sought to dissect and narrow down the substrate recognition requirements for each enzyme.

#### LazDE (azoline formation)

Because the action of LazDE is mostly independent of other enzymes, characterization of its substrate scope is relatively straightforward. First, we investigated whether any specific sequence elements in LazA CP are critical for LazDE activity, and prepared 4 LazA variants bearing randomized CPs with an Ala-Cys-Ala tripeptide grafted in the middle (Fig. 5a, LazA^CP1-4^). The peptides, produced with the FIT system, were incubated with LazDEF for 2 h, and reactions were analyzed by LC-MS. Efficient modification of 3 out of 4 tested substrates (Fig. 5a, LazA^CP1-3^) indicated that LazDE is a promiscuous enzyme able to act in “unfamiliar” sequence environments. Next, we studied relative heterocyclization rates for Cys vs. Ser and Thr. Consistent with a number of previously characterized YcaO enzymes,^41–43^ modification efficiency decreased from Cys to Thr to Ser (Fig. 5a, LazA^CP1^, LazA^CP1^ C7T, LazA^CP1^ C7S). Heterocyclization of Ser/Thr in sequences bearing a Thz in position +2 proceeded similarly (Fig. S42a).

**Figure 5.**
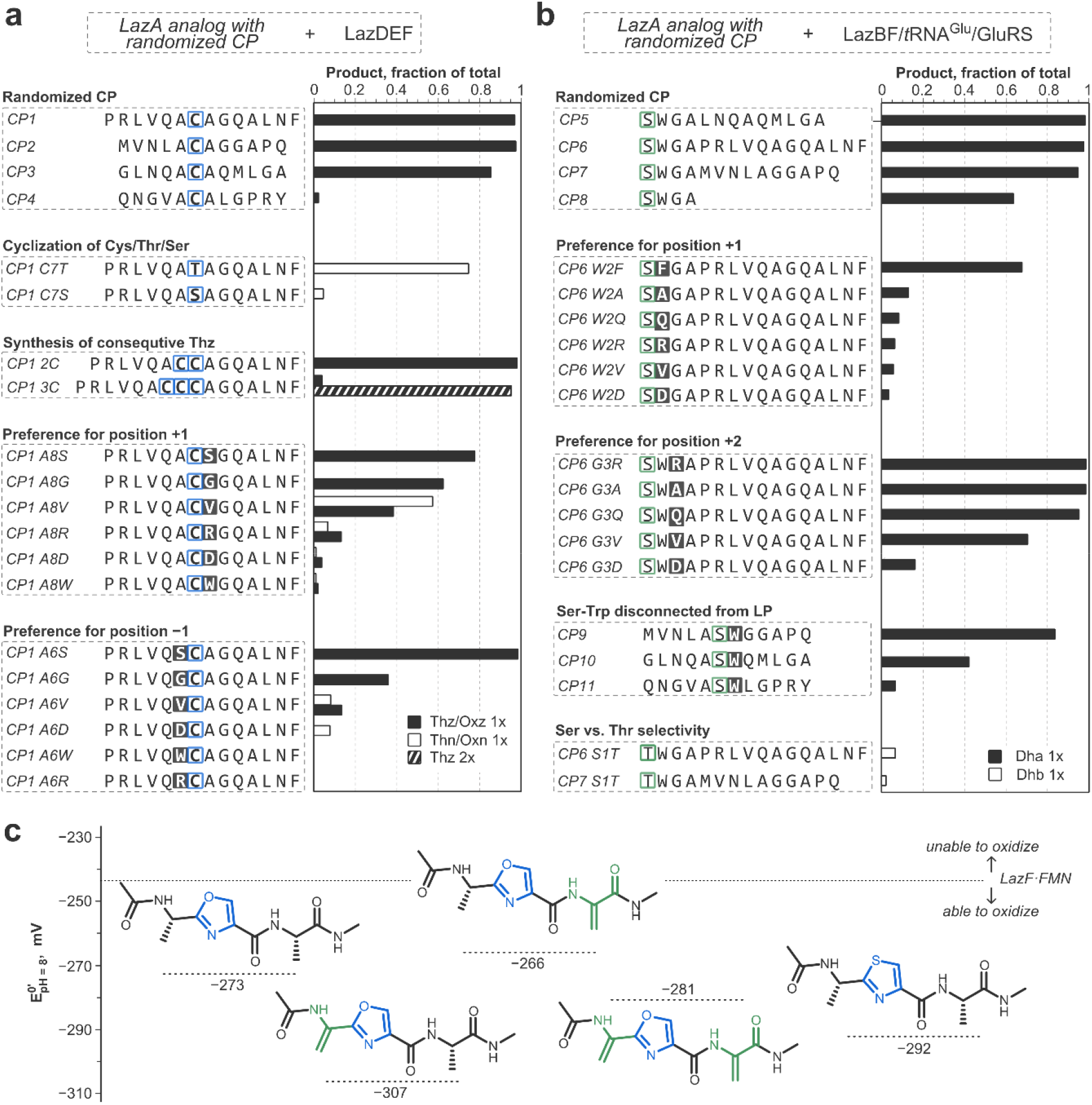
Substrate scope of LazDE (panel a), LazF dehydrogenase (panels a and c), and LazBF (panel b). a, b) In vitro translated LazA analogs with randomized CP sequences were incubated with either LazDEF (panel a) or LazBF/GluRS/tRNA^Glu^ (panel b) for 2 h. Reaction outcomes were analyzed by LC-MS and quantified as described in S.I. 2.5. The data reveal substrate preferences for individual Laz enzymes. The data for additionally tested substrates are summarized in Fig. S42. c) Comparison of the reduction potentials between LazF and several Oxz/Thz-containing tripeptides. The values for tripeptides were calculated as described in S.I. 2.6; for LazF — experimentally determined as described in S.I. 2.7 and Fig. S45. The oxidizing ability of LazF-bound FMN is sufficient to dehydrogenate all studied substrates, but the reaction potential increases when Dha flanks the substrate Oxn in position –1, i.e. Dha in position –1 promotes Oxn dehydrogenation.

The structure of lactazole A does not contain adjacent azoles, and even a prolonged incubation of LazA^wt^ with LazDEF does not result in consecutive heterocyclizations. To test whether LazDE is unable to form consecutive azoles, we prepared two substrates containing 2 or 3 Cys in a row (Fig. 5a; LazA^CP1^ 2C and LazA^CP1^ 3C). Treating these peptides with LazDEF led to one or two cyclodehydrations respectively, supporting our hypothesis. In contrast, substrates containing up to 4 Cys residues all separated by an Ala (Fig. S42b) resulted in formation of Thz at every Cys residue.

To ascertain whether the distance from the LP to the cyclizable amino acid affects modification rate, we prepared 4 LazA variants containing a single Cys in position 2, 4, 6 or 8 (Fig. S42e). LazA^LP ruler Cys2^ was unmodified after a 30-min treatment with LazDEF, suggesting that similar to PatD, a YcaO enzyme from a cyanobactin BGC,^42,44^ LazDE requires a spacer sequence between LP and residues undergoing modification, which explains how Ser1 escapes cyclodehydration. LazA^LP ruler Cys4^ and LazA^LP ruler Cys6^ gave over 80% Thz product compared to the 28% heterocyclization yield for LazA^LP ruler Cys8^, indicating that reaction rate decreases with distance from the LP, which rationalizes fast modification of Cys5 and Cys7 relative to Ser11 and Cys13 in LazA^wt^.

Finally, we studied the effect of amino acids adjacent to the cyclodehydration site using 12 single-point mutants of LazA^CP1^ (Fig. 5a). In position +1 (Cys-Xaa), LazDE preferred small/hydrophobic residues such as Ala, Ser, Gly and Val, while modification next to charged Asp and Arg as well as bulky Trp was impaired. In position –1, Ser or Ala were strongly preferred, while other mutants suffered from inefficient processing. In addition, we also grafted “native” tripeptides from LazA^wt^ into LazA^CP1^, and compared their relative modification rates after a 30-min treatment with LazDEF (Fig. S42d). These data uncovered a preference for Ser over Ala in position –1. The selectivity for Ser in position – 1 was especially apparent for Ser-Ser motifs. Whereas the Ala-Ser-Ala substrate remained essentially unmodified after a 2 h treatment with LazDEF, Ser-Ser-Ala and Ser-Ser-Ser peptides were quantitatively cyclodehydrated under the same conditions (Fig. S42c). Altogether, this study narrows down the primary recognition sequence of LazDE to the tripeptide Xaa_1_-(Cys/Thr/Ser)-Xaa_2_, where small Ser, Ala and Gly residues are strongly preferred in Xaa_1_ and Xaa_2_ positions, and Cys is modified faster than Thr and Ser (Fig. 6a).

**Figure 6.**
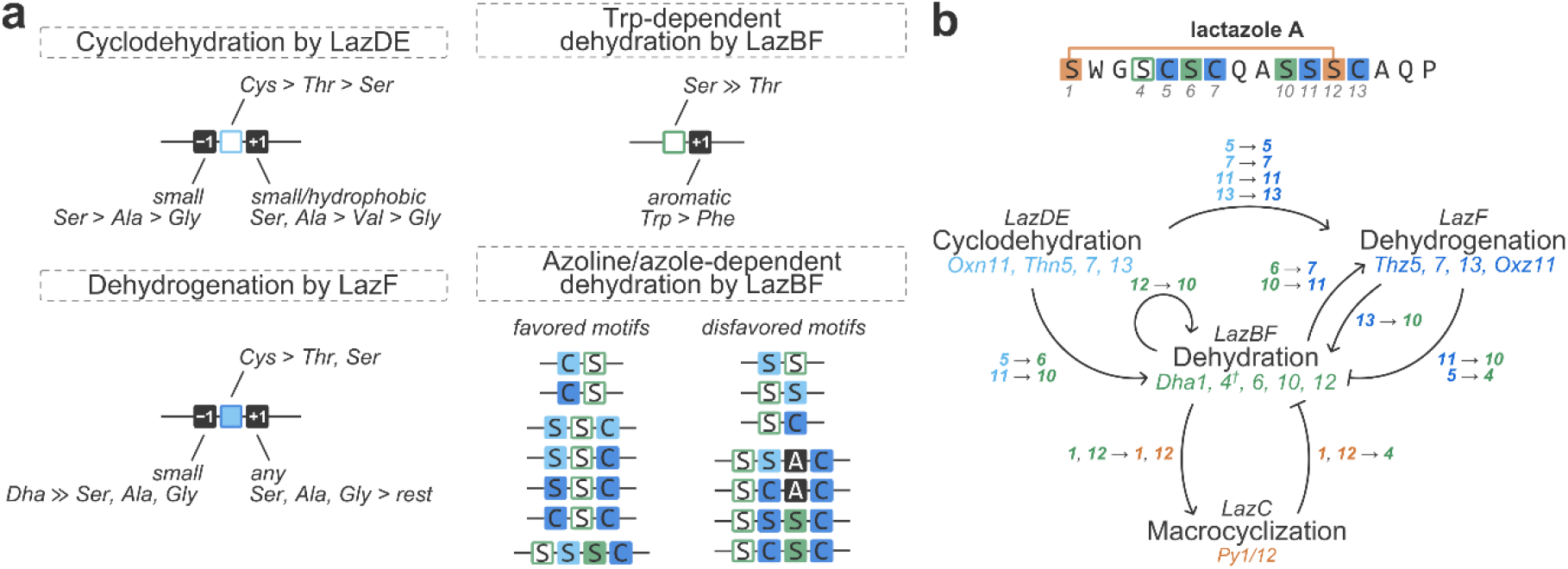
Dissection of lactazole biosynthesis. a) Summary of innate substrate preferences of LazDE, LazF dehydrogenase, and both LazBF Trp-dependent and azoline/azole-dependent modes of actions. Laz biosynthetic enzymes are in general characterized by “local” action, and require short 1–3 amino acid-long motifs for activity. b) Observed interactions between enzymatic activities in lactazole A biosynthesis. Pointed-end arrows indicate activation/promotion of the recipient activity at specified residues of LazA^wt^ CP (for instance, cyclodehydration at residue 5 enables dehydrogenation at residue 5, or dehydration at Ser12 promotes dehydration at Ser10). Blunt-end arrows indicate inhibition (in the broad meaning of the word) of the recipient activity at specified positions (for example, macrocyclization utilizes residue 1 and 12 and prevents dehydration of Ser4). ^†^Although Ser4 is not dehydrated in wild type lactazole A, formation of Dha4-lactazole is possible as discussed elsewhere in the text. This analysis visualizes the central role of LazBF during lactazole biosynthesis.

**LazBF (Dha formation)** has a two-fold activity profile. During lactazole biosynthesis, it converts Ser1 to Dha1 in an azoline/azole-independent fashion, while formation of remaining 3 Dha is azoline/azole dependent. We first focused on a more tractable azoline/azole-independent activity. As before, we began by preparing 3 LazA analogs with randomized CPs and grafted a Ser1-Trp2-Gly3-Ala4 tetrapeptide adjacent to the LP for each sequence (Fig. 5b; LazA^CP5–7^); one more substrate had the CP truncated after Ala4 (Fig. 5b; LazA^CP8^). The peptides were produced with the FIT system and treated with LazBF/GluRS/tRNA^Glu^ for 2 h. Efficient dehydration of all substrates, including the truncated peptide, indicated that LazBF is also a promiscuous enzyme that acts “locally”, i.e. it recognizes at most a few amino acids around the modification site. To further narrow down the recognition requirement, we prepared a series of single point mutants of LazA^CP6^ in positions 2 and 3 (Fig. 5b). In position +1, an aromatic residue was essential for good modification efficiency, although residual dehydration occurred in all cases. Position +2 tolerated more variation, and only a negatively charged Asp compromised processing. Additionally, we found that LazA mutants bearing a randomized CP with Ala-Ser-Trp grafted in the middle also underwent LazBF-catalyzed dehydration (Fig. 5b; LazA^CP9–11^), indicating that proximity to the LP is not a critical recognition element. Finally, we checked whether LazBF can accept Thr as a substrate to generate dehydrobutyrine (**Dhb**) residues. After a 2 h treatment, two Thr-containing substrates (Fig. 5b; LazA^CP6^ S1T and LazA^CP7^ S1T) were 7% and 3% modified, suggesting that LazBF strongly prefers Ser as a substrate, although synthesis of Dhb is possible. Combined, these results narrow down the primary recognition sequence of the azoline/azole-independent LazBF activity to the Ser-Trp dipeptide (Fig. 6a).

The substrate requirements for the azoline/azole-dependent Dha formation proved more difficult to generalize. First, we studied modification of single point Ala mutants of LazA^min^ and LazA^aux^ by LazBDEF/GluRS/tRNA^Glu^ using LazBF/GluRS/tRNA^Glu^ and LazDEF treatments as controls to gauge LazDE-dependent Dha formation (Fig. S43a, b). We found that Thz-Ser, Oxn-Ser-Thn/Thz, and Ser-Oxn-Dha-Thz motifs were generally dehydrated, but Ser-Thz, Oxn-Ser, Ser-Oxn, or Ser-Thz-Dha-Thz motifs were poor substrates (Fig. 6a). Processing of randomized CPs containing similar local environments recapitulated these findings (Fig. S43c, S44). However, based on these results, we were unable to generalize the rules governing Ser dehydration around azole/azolines: it is an intricately controlled activity, which would require a dedicated study. Nevertheless, these results help in rationalizing slow dehydration of Ser4 in Ser4-Thz5-Dha6-Thz7 motif during lactazole biosynthesis.

Notably, throughout the study, the product of LazB action, Ser(OGlu), was not detected so long as LazF was present in the enzyme mixture. This observation suggests that the burden of substrate discrimination lies primarily on LazB, because LazF appears to accept any Ser(OGlu)-containing peptide as a substrate.

#### LazF dehydrogenase (azole formation)

The aforementioned LazDE study provided a number of clues to the substrate scope of LazF dehydrogenase. First, like other Laz enzymes, LazF acts in unfamiliar sequence environments. The enzyme prefers small side chains such as Ala, Ser and Gly, on either side of the substrate azoline (Fig. 5a; Fig. 6a), and dehydrogenation of Thn happens much faster than Oxn or 5-MeOxn. Only one Oxn substrate not flanked by a Dha (Fig. S42c, LazA^CP1^ 2S) underwent up to 20% dehydrogenation after 2 h.

LazF-mediated dehydrogenation of Oxn11 emerged as the key step of lactazole biosynthesis, and thus, we sought to explore it in greater detail. How does Dha facilitate Oxn dehydrogenation given a relaxed specificity profile of LazF and its strong preference for Thn? We hypothesized that π-conjugation between the double bond of Dha in position–1 and the π-system of substrate Oxn may tune its reduction potential and facilitate dehydrogenation. To see whether this hypothesis is plausible, we calculated reduction potentials for several Oxz and Thz-containing tripeptides flanked by Dha or Ala on either side of the heterocycle (Fig 5c). We utilized density functional theory at the B3LYP level of theory and used the 6-311+G(d) basis set following previously established methods (S.I. 2.6).^45–47^

According to our calculations, Ala-Oxz-Ala had E^0’^ = –273 mV (pH 8), some 19 mV above a Thz analog, consistent with the notion that Thn undergoes dehydrogenation easier than Oxn.^48^ We also found that an Ala to Dha substitution in position –1 for an Oxz-containing peptide lowers its reduction potential by 34 mV, suggesting that the Dha-Oxn motif undergoes dehydrogenation easier than Ala-Thn. Dha in position +1 had a minor effect, and a tripeptide Dha-Oxz-Dha had E^0’^ = –281 mV (pH 8), halfway between Ala-Thz-Ala and Ala-Oxz-Ala. These data support our original hypothesis. To see whether these calculations are in line with the dehydrogenation ability of LazF, we experimentally determined reduction potential for LazF-bound FMN following a modified method of Massey,^49,50^ and established E^0’^ = –244±1 mV (pH 8; Fig. S45, S.I. 2.7). This value indicates that LazF-bound FMN provides sufficient oxidizing power to dehydrogenate all studied substrates, but the reaction potential increases from Ala-Oxn-Ala to Dha-Oxn-Dha to Ala-Thn-Ala, which helps in explaining Dha-dependent acceleration of Oxn oxidation. A similar mechanism might be at play during oxidation of Thn7. A Thn inside the Ala-Thn-Gln motif (Fig. S43a) was not dehydrogenated on a 30-min time scale, but during maturation of LazA^aux^ and LazA^wt^, an intermediate Thz5-Dha6-Thn7-Gln8 was too fast-lived to be captured.

The reduction potential determined here matches well with FMN-dependent azoline dehydrogenases from other RiPP classes,^51^ indicating that electrochemically LazF is not unique. Although Oxz-containing thiopeptides are not common, a number of such structures have been characterized.^25,52,53^ Most Oxz in thiopeptides, especially those in berninamycin-like structures,^54^ are flanked by a Dha residue in position –1 (Fig. S46), suggesting that Dha-assisted Oxn dehydrogenation may be a general phenomenon in biosynthesis of Dha/Oxz-containing natural products.

#### LazC (macrocyclization)

Even though we did not study LazC in isolation, a number of results clarify its function during lactazole assembly. First, LazC-catalyzed macrocyclization is fast compared to other Laz enzymes: during the time course studies, the macrocyclization substrate, LazA Dha1/Dha10-Oxz11-Dha12-Thz13 never accumulated to over 5% of total (Table S1, S3). Conversely, maturation of a LazA^min^ variant containing 5 mutations in the CP (Fig. S22c) stalled due to inefficient LazC action, with the macrocyclization precursor peptide comprising over 70% of total after a 3 h incubation. These data suggest that LazC might be more sensitive to the overall substrate structure than other Laz enzymes.

More importantly, LazC exerts kinetic control over lactazole assembly. The minimal recognition requirement around the 4π component of LazC (Oxz11-Dha12) ensures that as soon as Dha10-Oxz11-Dha12-Thz13 modifications are installed, macrocyclization will terminate the biosynthesis. This checkpoint controls the fate of Ser4, which in the absence of LazC is slowly dehydrated by LazBF. When LazC is present in the enzyme mixture, it macrocyclizes the substrate before this dehydration happens.

## Discussion

To the best of our knowledge, this study maps a multienzyme biosynthesis process of a RiPP at a single PTM resolution for the first time. Our results provide important clues on how Laz enzymes maintain integrity of lactazole assembly.

LazBF dehydrates 4 out of 6 Ser in the CP of LazA^wt^, and, as we previously found,^36^ depending on the order of enzyme addition, over- and underdehydrated thiopeptides bearing 5 or 3 Dha respectively can also be produced, hinting at a deep-seated cooperation between Laz enzymes. The results of this work reveal the extent of this cooperation. Coordination of the dehydratase activity emerges as the central theme of lactazole biosynthesis, and every enzyme is involved in the regulation of LazBF-mediated Dha synthesis (Fig. 6b). Innate substrate preferences of LazDE — specifically, its inability to cyclize amino acids adjacent to azoline/azoles, selectivity for Ser in position –1, and preference of Cys over Ser as substrate — discriminate a single Ser residue (Ser11) for cyclodehydration, leaving the remaining five Ser as potential LazBF substrates. Concomitantly, cyclodehydration of Cys5, 7 and 13 prepares Ser4, 6, 10 and 12 for azoline/azole-dependent dehydration. LazF dehydrogenase exerts a finer level control, promoting dehydration at Ser10 and impeding facile Dha synthesis at Ser4 through dehydrogenation of Thn13 and Thn5, respectively. Coupling of dehydrogenation to Dha formation, observed during the oxidation of Dha6-Thn7 and Dha10-Oxn11-Dha12-Thz13 motifs, serves as another important control mechanism, as it accelerates the biosynthesis and ensures that Dha10 is installed prior to LazC-catalyzed macrocyclization, preventing formation of the underdehydrated thiopeptide. Finally, through macrocyclization and LP cleavage, LazC emerges as a kinetic regulator of biosynthesis, restricting the action of LazBF at Ser4, which would otherwise be slowly dehydrated to Dha4. These control mechanisms are tightly coupled to the substrate preferences of LazBF, which — despite its apparent promiscuity — evolved to accurately sense PTMs introduced by other enzymes.

“Local” action also characterizes other Laz enzymes (Fig. 6a). Their primary substrate requirements may be narrowed down to 1–3 amino acids around the modification site, which allow them to act multiple times during lactazole biosynthesis and explain previously observed successful modification of substrates with extensively mutated CPs. The local action of the enzymes also explains the unusual punctuation of LazDE-mediated cyclizations by LazBF-catalyzed dehydrations and azoline dehydrogenation throughout the process: every enzyme acts as soon as it finds a substrate in a suitable local environment, regardless of the overall modification pattern. Nevertheless, enzymatic activities are tuned to the substrate, and with the exception of Ser1 dehydration, *in vitro* lactazole biosynthesis proceeds through a unified pathway, allowing a brief bifurcation during modification of Ser6 and Cys7.

Our results can explain lactazole biosynthesis without invoking a supramolecular enzyme complex, sometime hypothesized for thiopeptide assembly^55^ and often observed during biosynthesis of other RiPPs,^18,55,56^ although a possibility of such a complex is not ruled out. Instead, it appears that the enzymes communicate with each other via the substrate, and interactions between installed PTMs influence the assembly process in a major way. Such substrate-assisted assembly is emerging as a general phenomenon in RiPPs biosynthesis.^5,57^

Despite apparent structural similarities between thiopeptides, their biosynthetic logic appears to be fairly divergent. For example, during micrococcin maturation, Thz installation is separated from Ser dehydration by an obligatory C-terminal decarboxylation step,^21^ and in thiomuracin biosynthesis, glutamyl transferase TbtB is highly selective for the overall azole pattern.^20,58^ The *laz* BGC is also unique in its minimalistic size, unusual gene architecture (for instance, the fusion between glutamate elimination and dehydrogenase domains) and low sequence similarity to orthologous enzymes from other thiopeptide families.^40^ Perhaps, lactazole-like thiopeptides evolved from a goadsporin-like linear azole/Dha-containing peptide^59,60^ independently of other thiopeptide BGCs.

In summary, by identifying several control mechanisms responsible for integrity of lactazole assembly, this study begins to address how multiple promiscuous RiPP enzymes capable of competing over their substrate cooperatively modify it to converge on a single natural product. Our results rationalize the ability of *laz* BGC to synthesize diverse thiopeptides and inform on future bioengineering applications of Laz enzymes, including functional reprogramming of lactazole achieved by screening and *de novo* discovery of lactazole-inspired compounds from mRNA display libraries.

## Acknowledgements

We thank Dr. Kazuya Teramoto and Dr. Takayuki Kuge for their help with protein expression. This work was supported by CREST for Molecular Technologies, JST to H.S.; KAKENHI (JP16H06444 to H.S. and Y.G. and H.O.; JP17H04762, JP18H04382, JP19K22243, and JP20H02866 to Y.G.) from the Japan Society for the Promotion of Science (JSPS); a grant-in-aid from the Institute for Fermentation, Osaka (IFO) to H.O.; Amano Enzyme, Inc. to H.O.; and the A3 Foresight Program, JSPS to H.O. DFT computations were performed using Research Center for Computational Science, Okazaki, Japan.

