## Supplementary information for "Promiscuous enzymes cooperate at the substrate level en route to lactazole A"

### 1. General

Reagents were purchased from Nacalai Tesque, Wako Pure Chemical Industries, Sigma-Aldrich Japan, Kanto Chemical, or Watanabe Chemical Industries unless noted otherwise and were used as received. Oligonucleotides were purchased from Eurofins Genomics (OPC purification grade) and were used without further purification. All PCR amplifications were carried out in a BioER TC-96GHBC thermal cycler. Protein purification and synthesis *of S. lactacystinaeus* $tRNA^{Glu}$ were done as previously described.[1]

NMR spectra ($^{1}H$ NMR at 300 MHz and $^{13}C$ NMR at 75 MHz) were recorded on a Bruker Model Avance DMX 300 Spectrometer. MALDI-TOF mass spectra were recorded on Autoflex II TOF/TOF (Bruker Daltonics) in the positive reflector mode using external calibration with protein calibration standard I (Bruker Daltonics). Sinapic acid was used as matrix.

### 2. Methods

#### 2.1. Preparation of synthetic DNA

Linear double-stranded DNA encoding T7 promoter upstream of *lazA* ORF and its mutants were assembled by PCR from synthetic single-stranded DNA oligonucleotides using Taq polymerase. All PCR were performed in 10 mM Tris (pH 8.4), 50 mM KCl, 0.1% (v/v) Triton X-100, 2.5 mM $MgCl_2$, 250 μM each dNTP supplemented with 500 nM of appropriate primers and Taq DNA polymerase (standard PCR conditions). Three stage thermal cycling included a denaturing step at 95°C for 40 s, annealing at 52°C for 40 s, and extension at 72°C for 40 s. The list of all oligonucleotides and assembly schemes can be found in Tables S5 and S6.

For *lazA*$^{wt}$, forward and reverse primers were annealed and extended in the primer extension step. PCR solution (100 μL) containing 500 nM primers was incubated at 95°C for 60 s. Then, five cycles of annealing (52°C for 60 s) and extension (72°C for 60 s) were performed, and 1 μL of the product was used as a template for the next step. The first PCR was performed under the standard conditions with 5 cycles of amplification. The second PCR was performed on a 1000 μL scale using 5 μL of the first PCR product with 14 cycles of amplification. Correct assembly was confirmed by 3% agarose gel electrophoresis. PCR product was extracted by phenol/chloroform/isoamyl alcohol (25:24:1, saturated with 10 mM Tris (pH 8.0), 1 mM EDTA), and then by chloroform/isoamyl alcohol (24:1). Extracted DNA was precipitated with ethanol, washed with 70% ethanol in water (v/v), and dissolved in 100 μL of water for storage.

Most other templates were assembled using *lazA*$^{wt}$ or *lazA*$^{min}$ as a template in one or two steps. For single step mutagenesis, reactions were performed on a 200 μL scale with 1 μL of 1:100 diluted template DNA and 14 cycles of amplification. For two step procedures,

100 µL of PCR solution containing 0.5 µL of 1:10 diluted template DNA was amplified for 10 cycles, followed by a 14-cycle amplification of the PCR product from the first step (1 µL in 200 µL PCR solution). Analysis of amplification and DNA isolation were performed as above. These DNA templates were used for *in vitro* translation without concentration adjustment or further purification.

#### 2.2. In vitro translation

A transcription-coupled *in vitro* translation system was reconstituted as previously reported[1,2] by mixing purified ribosome, enzymes and translation factors. The final reaction mixture contained 50 mM HEPES-KOH (pH 7.6), 100 mM KOAc, 2 mM guanosine triphosphate (GTP), 2 mM adenosine triphosphate (ATP), 1 mM cytidine triphosphate (CTP), 1 mM uridine triphosphate (UTP), 20 mM creatine phosphate, 12 mM $Mg(OAc)_2$, 2 mM spermidine, 2 mM dithiothreitol (DTT), 1.5 mg/mL *E. coli* total tRNA (Roche), 1.2 µM ribosome, 0.6 µM methionyl-tRNA formyltransferase (MTF), 2.7 µM prokaryotic initiation factor-1 (IF1), 0.4 µM prokaryotic initiation factor-2 (IF2), 1.5 µM prokaryotic initiation factor-3 (IF3), 10 µM elongation factor thermo unstable (EF-Tu), 10 µM elongation factor thermo stable (EF-Ts), 0.26 µM elongation factor G (EF-G), 0.25 µM release factor 2 (RF2), 0.17 µM release factor 3 (RF3), 0.5 µM ribosome recycling factor (RRF), 4 µg/mL creatine kinase, 3 µg/mL myokinase, 0.1 µM pyrophosphatase, 0.1 µM nucleotide-diphosphatase kinase, 0.1 µM T7 RNA polymerase, 0.73 µM alanyl-tRNA synthetase (AlaRS), 0.03 µM ArgRS, 0.38 µM AsnRS, 0.13 µM AspRS, 0.02 µM CysRS, 0.06 µM GlnRS, 0.23 µM GluRS, 0.09 µM GlyRS, 0.02 µM HisRS, 0.4 µM IleRS, 0.04 µM LeuRS, 0.11 µM LysRS, 0.03 µM MetRS, 0.68 µM PheRS, 0.16 µM ProRS, 0.04 µM SerRS, 0.09 µM ThrRS, 0.03 µM TrpRS, 0.02 µM TyrRS, 0.02 µM ValRS, 500 µM each proteinogenic amino acid and 100 µM 10-formyltetrahydrofolate (10-HCO-H4 folate). Translation reactions were performed at 37°C for 50 to 60 min with 1 µL of *lazA* variant template DNA, for a total translation volume of 5 µL. For DNA templates encoding an amber stop codon, RF1 (1 µM final concentration) was additionally supplemented to the translation mixture.

#### 2.3. Enzymatic reactions

Enzymatic reactions were performed with 2.5 µL translation product (Methods 2.2). The substrate was placed on ice, and 12.5 µL of ice-cold enzyme mixture (see below) was added. The mixture was incubated in a 25 °C thermostatic incubator for 5 min to 17 h (specified on a case by case basis), and then transferred on ice. Ice cold methanol containing 30 mM IAA (16 µL) was added to quench the reaction and precipitate protein and nucleic acid. After 5 min on ice, the precipitate was separated by centrifugation (15300 g for 4 min at 25 °C), and the supernatant was carried forward for LC-MS analysis.

Full enzyme mixture consisted of 2.4 µM LazB, 2.4 µM LazC, 1.2 µM LazD, 1.2 µM LazE,

2.4 µM LazF, 1.2 µM *S. lividans* GluRS, 12 µM *S. lactacystinaeus* $tRNA^{Glu}$ in 60 mM Tris buffer (pH 8.0) supplemented with 12 mM $MgCl_2$, 6 mM ATP, and 1.2 mM DTT. Specific components were omitted to prepare partial enzyme mixes and to perform stepwise treatments as specified for each experiment individually. The "no enzyme" treatments were done identically to the enzymatic reaction, except the enzymes and $tRNA^{Glu}$ were omitted from the mixture.

#### 2.4. LC-MS and DDA MS/MS analysis

Reaction outcomes were analyzed using Waters Xevo G2-XS QTof instrument equipped with Acquity I-Class UPLC system. HPLC was done on an Acquity UPLC Peptide BEH C18 column (dimensions: 150 x 2.1 mm; pore size: 300Å ; particle size: 1.7 µm) using 0.1% (v/v) formic acid (**FA**) in water (solvent A) or 0.1% (v/v) FA in acetonitrile (solvent B) as a mobile phase unless noted otherwise. Analysis was performed at 60°C and 250 µL/min flow rate running the following gradient: 1% B for 2 min; 1 to 81% B over 20 min; 95% B for 2 min; 1% B for 6 min (total run time: 30 min; **HPLC method 1**). For time course experiments, a shallower gradient was used for better separation of linear intermediates: 1% B for 2 min; 1 to 25% B over 4 min; 25 to 41% B over 16 min; 41 to 81% B over 6 min; 1 to 25% B over 4 min; 95% B for 2 min; 1% B for 6 min (total run time: 36 min; **HPLC method 2**).

MS analysis was done in a positive polarity/high sensitivity mode with a 0.3 s scan time. Capillary voltage was set to 700 V; ESI source and desolvation temperatures were 120 and 400°C, respectively. Manufacturer-supplied GFB was used as a lockspray standard for continuous mass axis referencing, and the lockspray setup procedure was performed according to the manufacturer's instructions prior to every run.

For tandem mass spectrometry, a data-dependent acquisition (DDA) method was used. CID fragmentation was triggered in real time if detected ions met the following conditions: ion intensity exceeded $4 \cdot 10^4$ ions, and z=4. MS/MS spectra were acquired with a 2 s scan time, with parameterized collision energy values. In method 1, collision energies were set to ramp from 6-8 to 30-40 eV over the range of acquired *m/z* values (200 to 2000); in method 2, these values were 6-8 to 28-33 eV, and in method 3, 6-8 to 26-29 eV. Samples subject to MS/MS were reanalyzed three times to acquire MS/MS spectra for each method. All acquired spectra were considered during data analysis, but reported are the most informative ones for brevity. In select cases, individual MS/MS spectra were stacked to improve signal to noise ratio.

#### 2.5. LC-MS and DDA MS/MS data analysis

LC-MS data was analyzed with MassLynx v.4.1. Product distributions were quantified by integrating areas under LazA-derived peaks. To this end, individual compound areas were calculated by summing areas under extracted ion current (EIC) chromatogram peaks for complete charged series, i.e.:

$$A^{pep} = \sum_{z} A_z^{EIC}(pep)$$

and LC-MS yields were estimated as a fraction of total, i.e.:

$$y^{pep} = \frac{A^{pep}}{\sum A}$$

Due to potential differences in ionization efficiencies of analyzed peptides, we note that this analysis should be treated as semi-quantitative rather than strictly quantitative. For thiopeptides, yields were not calculated for the same reason. Reported are full ion intensity areas for thiopeptide peaks normalized over the total ion intensity of each sample.

$^{br}$EIC chromatograms were generated as previously reported[1] with *m/z* ±100 (±400 Da) tolerance window; for $^{nr}$EIC, the m/z tolerance window was set either to 1.0 (for linear intermediates) or 0.1 for thiopeptides.

In some cases, an inseparable by HPLC mixture of Thn and Thz-containing peptides formed. Because molecular weight of these peptides differs by only 2.01 Da, their mass spectra are also convolved, which precludes their direct quantification. To quantify the Thn/Thz ratio in such cases, isotope envelope deconvolution was performed. A mass spectrum integrated over both peaks (usually at $z$=4) was subjected to least square regression to optimize for the fraction of Thn using the generalized reduced gradient (nonlinear) optimization (Microsoft Excel, Solver) by fitting to a combination of two theoretical mass spectra: one for a Thn-containing peptide, and another for Thz. Theoretical spectra were computed with enviPat Web 2.4.[3] See also Fig. S15.

MS/MS assignments were done manually. For a given peptide, a series of possible PTM patterns was generated, and for each of these patterns, ladders of *b*- and *y*-ion series were generated. Calculated ion ladders were compared against experimental spectra, and the ion ladder leading to the best match was used to make spectral assignments.

#### 2.6. Calculation of reduction potentials for Oxz/Thz-containing tripeptides

Gaussian16 Rev. B.01 software package was used to perform DFT calculations, and the reduction potential computation followed previously established protocols.[4] Molecular structures in water phase were optimized in the PCM solvation model at the B3LYP level of theory using 6-311+G(d) basis set. For the reduction half reaction

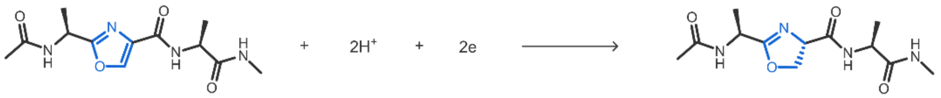

the free Gibbs energy change is related to the reduction potential:

$$\Delta G^{0'} = \Delta G^{0}_{Oxn} - (\Delta G^{0}_{Oxz} + 2\Delta G^{0'}_{H^+(aq)}) = -nFE^{0'}\text{ , where}$$

$$\Delta G^{0'}_{H^+(aq)} = \Delta G^{0}_{H^+(aq)} + 0.059\cdot pH$$

$$\Delta G^{0}_{H^+(aq)} = -264.6\ \text{kcal/mol}$$

Thus, the reduction potential can be calculated from

$$E^{0'}(pH) = \frac{\Delta G^{0}_{Oxn} - \Delta G^{0}_{Oxz} - 2\Delta G^{0'}_{H^+(aq)} - 0.059\cdot pH}{-nF}$$

Gibbs free energies at 298.15 K for oxidized and reduced forms were calculated as the sum of the energy in solution and the thermal correction of Gibbs free energy, and standard reduction potentials were calculated at pH 8. Optimized molecular geometries and $\Delta G^0$ values are summarized in section 3.7.

#### 2.7. Determination of the reduction potential of LazF-bound FMN

The reduction potential of LazF-bound FMN was carried out following the method of Massey with modifications.[5,6] A 1 mL mixture of 50 μM LazF, 15 μM phenosafranine and 2 μM benzyl viologen dichloride in 25 mM HEPES buffer (pH 8.0) containing 500 mM NaCl and 5% (v/v) glycerol was made anaerobic inside a nitrogen-flushed glove box. UV-Vis spectra were recorded using a Nanodrop 2000c spectrophotometer (Thermo Scientific) in a 1 cm pathlength cuvette at 25 °C. Stepwise chemical reduction was performed with 5 mM solution of sodium dithionite ($Na_2S_2O_4$) dissolved in the same buffer. To the protein/dye mixture, sodium dithionite (0.5–2 μl) was added once every 5 min, and a 300–800 nm absorption spectrum was recorded prior to the addition of the next aliquot. Titration continued until both phenosafranine and FMN were fully reduced (Fig. S45).

Absorption spectra for fully oxidized and fully reduced FMN/LazF and phenosafranine were recorded separately and were used to calculate extinction coefficients (Fig. S57). All acquired spectra were baseline-corrected at 700 nm. Reduction of phenosafranine was monitored at 540 nm, where spectral interference from FMN is negligible. For FMN, the ratio of oxidized to reduced forms was calculated at 457 nm after correcting for phenosafranine absorption. The reduction potential was calculated at pH 8 using phenosafranine as a reference ($E^{0'}_{pH=8} = -281.5$ mV) from a linear regression to the following equation:

$$\log_{10}\left(\frac{[ox(LazF)]}{[red(LazF)]}\right) = \log_{10}\left(\frac{[ox(dye)]}{[red(dye)]}\right) - \frac{nF}{2.3RT}\left(E^{0'}_{pH=8}(LazF) - E^{0'}_{pH=8}(dye)\right)$$

The reported $E^{0'}_{pH=8}$ (LazF) value is the average of three independent measurements ± one standard deviation.

### 3. Experimental data

#### 3.1. PTM assignment scheme

All studied PTM types had unique mass shifts due to IAA alkylation on unmodified Cys residues and selective oxazoline hydrolysis under the HPLC conditions. Efficient IAA alkylation from 50% methanol at 4°C proceeded quantitatively, simplifying assignment of PTM patterns in partially modified precursor peptides. Figure S1 summarizes mass shifts for studied PTM types.

With the exception of Oxn (see below), PTMs were chemically stable during HPLC-MS. After quenching the reactions, Dha (if unconjugated to azoline/azole) slowly underwent Michael addition to cysteine, β-mercaptoethanol or DTT included in the reaction mixture. These thiol adducts were treated as Dha for quantification purposes.

Oxn quantitatively hydrolyzed during our HPLC-MS conditions (0.1% FA additive at 60 °C), and reverted to the product whose mass was identical to unmodified Ser (Fig. S2b). Acidic hydrolysis of Oxn is a well-studied phenomenon, and has been shown before to quantitatively yield β-aminoesters over β-hydroxyamides (Fig. S2a).[7,8] These β-aminoester Oxn hydrolysis products behaved differently from the starting material Ser-containing peptides on HPLC, and had a characteristic retention time shift, eluting as much as 1 min earlier. Because other PTMs shifted peptide's retention time in the opposite direction, Oxn modification was assigned based on HPLC retention time (Fig. S2c, d). Changing the mobile phase additive from FA to acetic acid (0.1%) only partially alleviated hydrolysis, and resulted in a mixture of a β-aminoester and an intact Oxn (Fig. 2c). A side-by-side LC-MS and MALDI-TOF analysis of reaction mixtures confirmed the presence of an extra dehydration invisible by LC-MS (Fig. S2c, d). In contrast to Oxn, Thn proved to be stable during HPLC. In most cases, less than 2% hydrolysis product was observed.

To confirm that an early-eluting isomer of the starting material corresponds to *in situ* Oxn formation followed by its hydrolysis, we synthesized an authentic Oxn dipeptide and analyzed it by LC-MS as described in section 3.2.

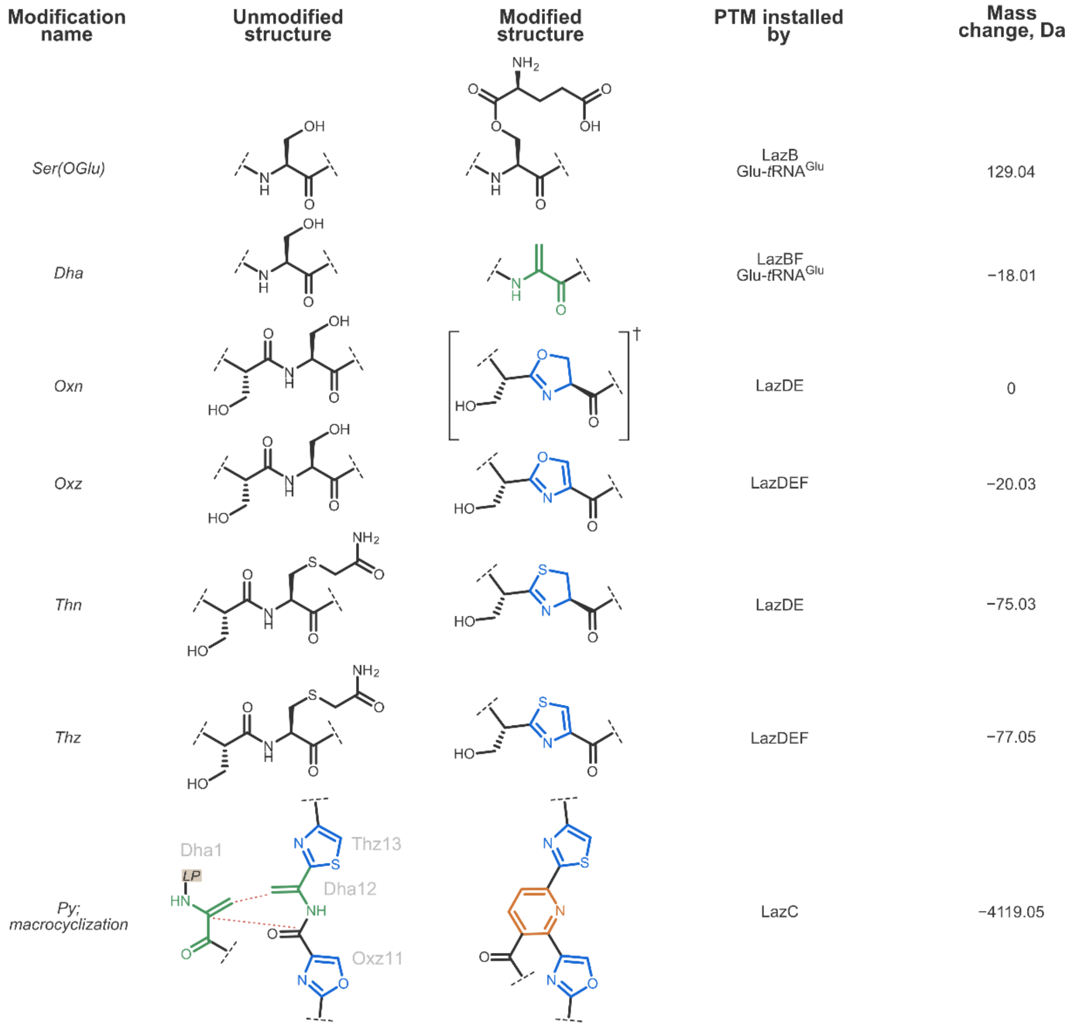

**Figure S1**. PTM assignment scheme. Every PTM had a unique mass shift, which simplified the product assignment process. †See text above for discussion of Oxn stability.

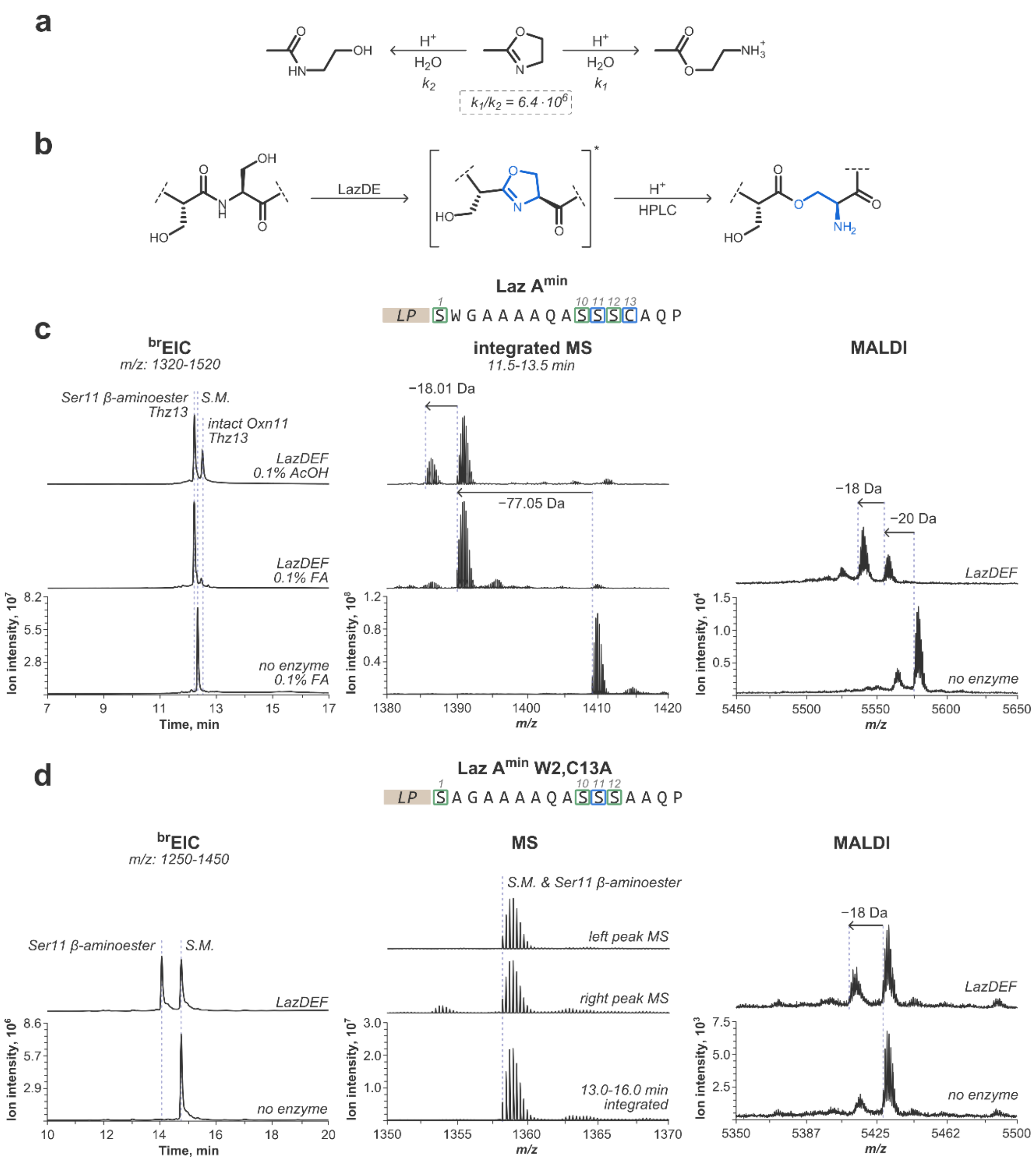
a
H+
H2O
k2
k1
k1/k2 = 6.4 · 10^6
b
LazDE
H+
HPLC
Laz A^min
LP S W G A A A A Q A S S S C A Q P
c
brEIC
m/z: 1320-1520
Ser11 β-aminoester
Thz13
S.M.
intact Oxn11
Thz13
LazDEF
0.1% AcOH
LazDEF
0.1% FA
no enzyme
0.1% FA
Ion intensity, 10^7
Time, min
integrated MS
11.5-13.5 min
−18.01 Da
−77.05 Da
Ion intensity, 10^8
m/z
MALDI
−18 Da
−20 Da
LazDEF
no enzyme
Ion intensity, 10^4
m/z
d
Laz A^min W2,C13A
LP S A G A A A A Q A S S S A A Q P
brEIC
m/z: 1250-1450
Ser11 β-aminoester
S.M.
LazDEF
no enzyme
Ion intensity, 10^6
Time, min
MS
S.M. & Ser11 β-aminoester
left peak MS
right peak MS
13.0-16.0 min
integrated
Ion intensity, 10^7
m/z
MALDI
−18 Da
LazDEF
no enzyme
Ion intensity, 10^3
m/z

**Figure S2**. Analysis of Oxn hydrolysis during HPLC-MS. a) A kinetic scheme of Oxn hydrolysis at low pH. Formation of a β-aminoester is favored by a factor of $10^6$. Data from ref. [7]. b) Formation and hydrolysis of Oxn during *in vitro* lactazole biosynthesis and the following LC-MS analysis. c) A typical example of Oxn hydrolysis. $LazA^{min}$ precursor peptide was *in vitro* translated and incubated with LazDEF for 2h at 25 °C. LCMS analysis using 0.1% FA as a mobile phase additive showed a single peak eluting earlier than the starting material (S.M.), and its molecular weight consistent with the formation of Thz13. Rerunning the same sample using 0.1% acetic acid as an additive resulted in two products, one of which had an extra dehydration and eluted after S.M. These products were assigned as β-aminoester at Ser11/Thz13 and intact Oxn11/Thz13. MALDI-TOF analysis of the same reaction mixture confirmed that Oxn11/Thz13 is the major product in the reaction mixture. d) Another illustration of Oxn hydrolysis. $LazA^{min}$ W2,C13A was incubated with LazDEF for 2h at 25 °C. LC-MS analysis results in two peaks with identical molecular weight, the S.M., and its early-eluting isomer. The early-eluting isomer was assigned as β-aminoester at Ser11 derived from Oxn11. MALDI-TOF analysis of the same reaction mixture confirmed that a mixture of Oxn11 and S.M. formed during the reaction.

#### 3.2. Chemical synthesis and LCMS analysis of Boc-Phe-Oxn-OMe

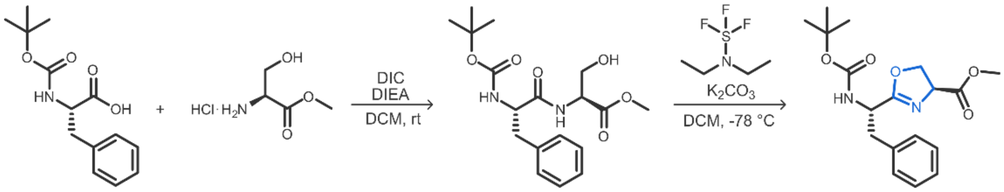

**Scheme S1**. Synthesis scheme for Boc-Phe-Oxn-OMe.

Boc-Phe-Ser-OMe dipeptide and Boc-Phe-Oxn-OMe were synthesized following an established procedure.[9] Briefly, diisopropylethylamine (0.3 ml) was added to a solution of Boc-$^{L}$Phe-COOH (420 mg) in DCM (2.5 ml). The mixture was cooled to 4 °C, and DIC (0.28 ml) was added dropwise. After 1h at 4 °C, $^{L}$Ser-OMe·HCl (0.26 g) was added in one portion, and the solution was stirred at room temperature overnight. Precipitate was filtered, and the solvent was removed under reduced pressure. The residue was redissolved in ethyl acetate and extracted with 10% citric acid, saturated $NaHCO_3$ and brine. The organic extract was dried over $MgSO_4$ and purified on a Biotage Isolera™ Spektra flash purification system equipped with a Biotage SNAP Ultra 10 g column using an ethyl acetate/hexane solvent system. The product was obtained as a white solid (336 mg; 58%).

In the second step, diethylaminosulfur trifluoride (39.7 µl) was added dropwise to a cold (–78 °C) solution of Boc-Phe-Ser-OMe (100 mg) in DCM (2.5 ml). After stirring for 1 h at -78 °C, anhydrous $K_2CO_3$ (57 mg) was added in one portion and the mixture was allowed to warm to room temperature. The reaction was poured into saturated $NaHCO_3$, and the biphasic mixture was extracted with DCM. The combined organic extracts were dried over $MgSO_4$, filtered, and concentrated under reduced pressure. Flash purification over $SiO_2$ as above afforded Boc-Phe-Oxn-OMe as an off-white solid (77 mg; 81%), whose NMR spectra were consistent with literature data.[9]

$^{1}$H NMR (300 MHz, $CDCl_3$): δ = 7.29–7.10 (m, 5 H, Ph), 5.17 (d, J= 8.0 Hz,1 H, NH), 4.69 (m, J= 9.8 Hz, 2H, CHCO2Me, CHBn), 4.59 (t, J= 8.3 Hz, 1H, OCHH), 4.44 (dd, J= 8.7, 10.5 Hz, 1H,OCHH), 3.74 (s, 3 H, CO2CH3), 3.14 (dd, J= 5.7, 13.6 Hz, 1 H, CHHPh), 3.01 (dd, J= 5.4, 13.6 Hz, 1 H, CHHPh), 1.40 [s, 9H,C(CH3)3] ppm.

$^{13}$C NMR (75 MHz, $CDCl_3$): δ = 170.9, 169.3, 154.9, 135.8, 129.6, 128.3, 126.8, 79.8, 70.0, 67.8, 52.7, 49.7, 38.8, 28.3.

Boc-Phe-Ser-OMe and Boc-Phe-Oxn-OMe were analyzed by LC-MS under standard conditions (0.1% FA as an additive; 60 °C; Fig. S5). Boc-Phe-Oxn-OMe quantitatively hydrolyzed to an early-eluting β-aminoester, confirming our reasoning from section 3.1.

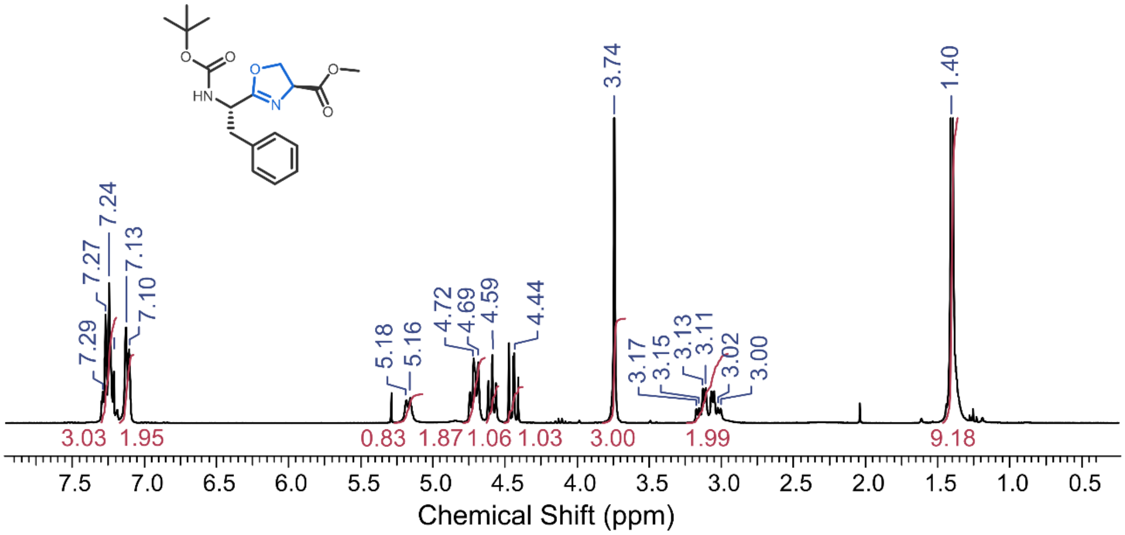

**Figure S3**. [1]H NMR spectrum (300 MHz; $CDCl_3$) of Boc-Phe-Oxn-OMe.

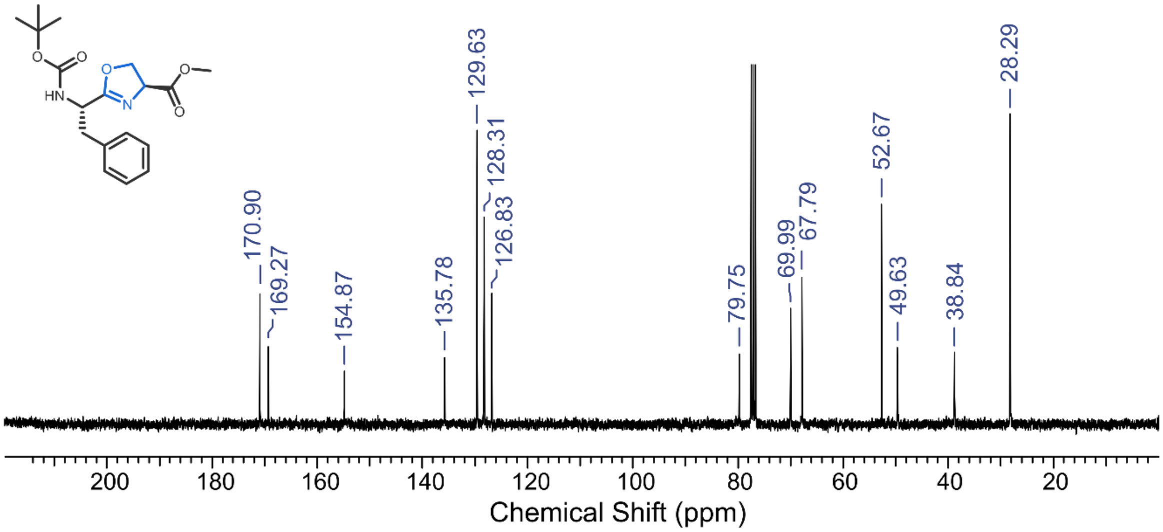

**Figure S4**. [13]C NMR spectrum (75 MHz; $CDCl_3$) of Boc-Phe-Oxn-OMe.

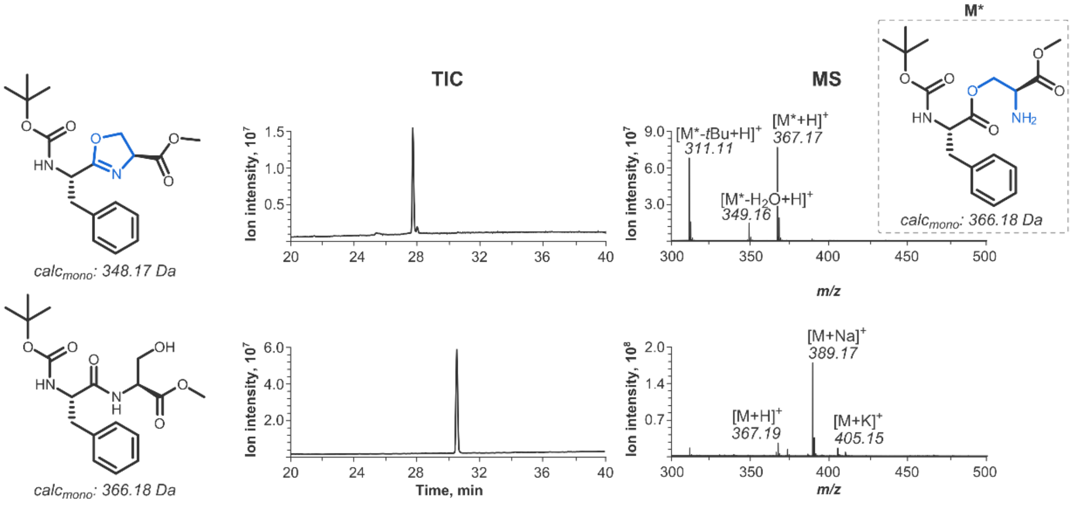

**Figure S5.** LC-MS analysis of Oxn hydrolysis using a synthetic Oxn standard. Boc-Phe-Oxn-OMe (above) and Boc-Phe-Ser-OMe (below) were synthesized (section 3.2) and analyzed by LC-MS under standard conditions (0.1% FA as an additive; C18 phase; 60 °C). Consistent with our results from LazA modification experiments, the Oxn product eluted earlier than the "starting material" Ser dipeptide, and had molecular weight consistent with the formation of a β-aminoester, supporting the occurrence of quantitative hydrolysis of Oxn under our LC-MS conditions.

#### 3.3. LazA$^{min}$ time course assignments

Assignments of PTM patterns on captured intermediates were done following the scheme outlined in section 3.1 and from MS/MS data. Table S1 summarizes observed intermediates and shunt products. Figures S6-S22 show data to support these annotations.

Formation of various lanthipeptides, stemming from a non-enzymatic cyclization of Dha1 to Cys13, was the major failure mode of biosynthesis. For LazA$^{min}$, these products comprised around 5% of total after a 60-min incubation with LazBCDEF/GluRS/tRNA$^{Glu}$. Assigning a peak as a lanthipeptide was done on the basis of its molecular weight, MS/MS data (which showed abrogation of fragmentation consistent with the formation of a macrocycle between residues 1 and 13), and steady accumulation of these products over the course of biosynthesis. In contrast to real intermediates, these products were biosynthetic "dead ends", and were never consumed even after extending reaction times.

**Table S1**. (see next page). Progress of LazA$^{min}$ biosynthesis in the FIT-Laz system. For each intermediate, *m/z* values correspond to the monoisotopic mass peaks. RT stands for HPLC retention time (method B); a.u. stands for arbitrary units (see also section 2.5).

| *m/z* | RT, min | assignment | notes | MS/MS confirmation | reaction time, min | | | | | |
|---|---|---|---|---|---|---|---|---|---|---|
| | | | | | 0 | 5 | 15 | 30 | 45 | 60 |
| 1026.26 | 15.41 | LP-$NH_2$ | Macrocyclization product | | 0.0 | 0.0 | 4.8 | 35.5 | 65.6 | 82.7 |
| 1371.41 | 17.92 | Dha1, 10, 12 Oxz11 Thz13 | Macrocyclization substrate | Fig. S14 | 0.0 | 0.0 | 1.3 | 4.5 | 1.1 | 1.3 |
| 1375.90 | 17.60 | Dha10, 12 Oxz11 Thz13 | | Fig. S13 | 0.0 | 0.4 | 1.6 | 1.6 | 0.8 | 0.0 |
| 1380.90 1381.41 | 15.85 | Dha1, 12 Oxn11 Thn13/Thz13 | 32% Thz 68% Thn Fig. S15b | Fig. S12 for Oxn-product | 0.0 | 0.0 | 3.1 | 4.0 | 2.0 | 1.0 |
| 1385.41 1385.91 | 15.59 | Oxn11 Dha12 Thn13/Thz13 | 29% Thz 71% Thn Fig. S15a | Fig. S11 for Oxn-product | 0.0 | 0.3 | 11.2 | 20.4 | 13.1 | 5.9 |
| 1390.41 | 15.25 | Oxn11 Thn13 | | Fig. S10 | 0.0 | 1.5 | 15.2 | 9.3 | 4.0 | 0.7 |
| 1390.41 | 15.88 | Dha1 to Cys13 Lanthipeptide | isomer 1; shunt product | | 0.0 | 1.7 | 3.0 | 2.5 | 2.8 | 2.8 |
| 1390.41 | 16.07 | | isomer 2; shunt product | | 0.0 | 0.1 | 1.2 | 2.0 | 2.3 | 2.2 |
| 1404.68 | 16.20 | Dha1 | | Fig. S9 | 0.0 | 4.6 | 12.7 | 3.6 | 1.0 | 0.3 |
| 1409.18 | 15.14 | Oxn11 | | | 0.0 | 1.1 | 3.5 | 2.0 | 0.8 | 0.2 |
| 1409.18 | 15.83 | S.M. | | Fig. S8 | 100.0 | 88.9 | 38.2 | 8.4 | 3.6 | 1.3 |
| | | minor products, combined | | | 0.0 | 1.3 | 4.5 | 6.0 | 3.1 | 1.6 |
| | | | | | thiopeptide intensity, a. u. | | | | | |
| 682.27 | 9.28 | thiopeptide | | | 0.0 | 0.0 | 1.4 | 12.2 | 22.3 | 27.4 |

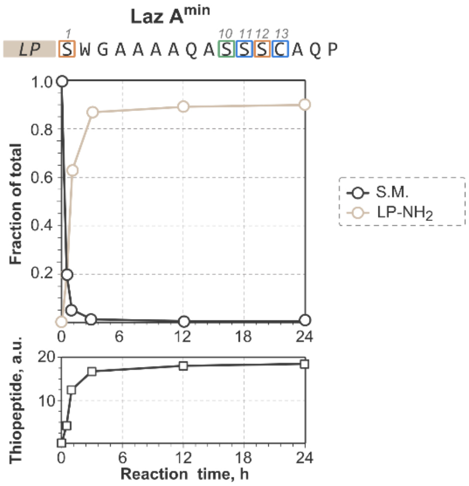

**Figure S6**. Preliminary time course experiment. Translation-derived precursor peptide, LazA$^{min}$, was incubated with LazBCDEF/GluRS/tRNA$^{Glu}$ for specified time and the outcomes were analyzed by LC-MS. Consumption of the starting material (S.M.), and the final step of biosynthesis, i.e. the macrocyclization reaction leading to the formation of the thiopeptide (lactazole S4-C7A) and LP-$NH_2$, were monitored to estimate the overall biosynthesis rate. These data suggest the LazA$^{min}$ maturation is complete within 3 h.

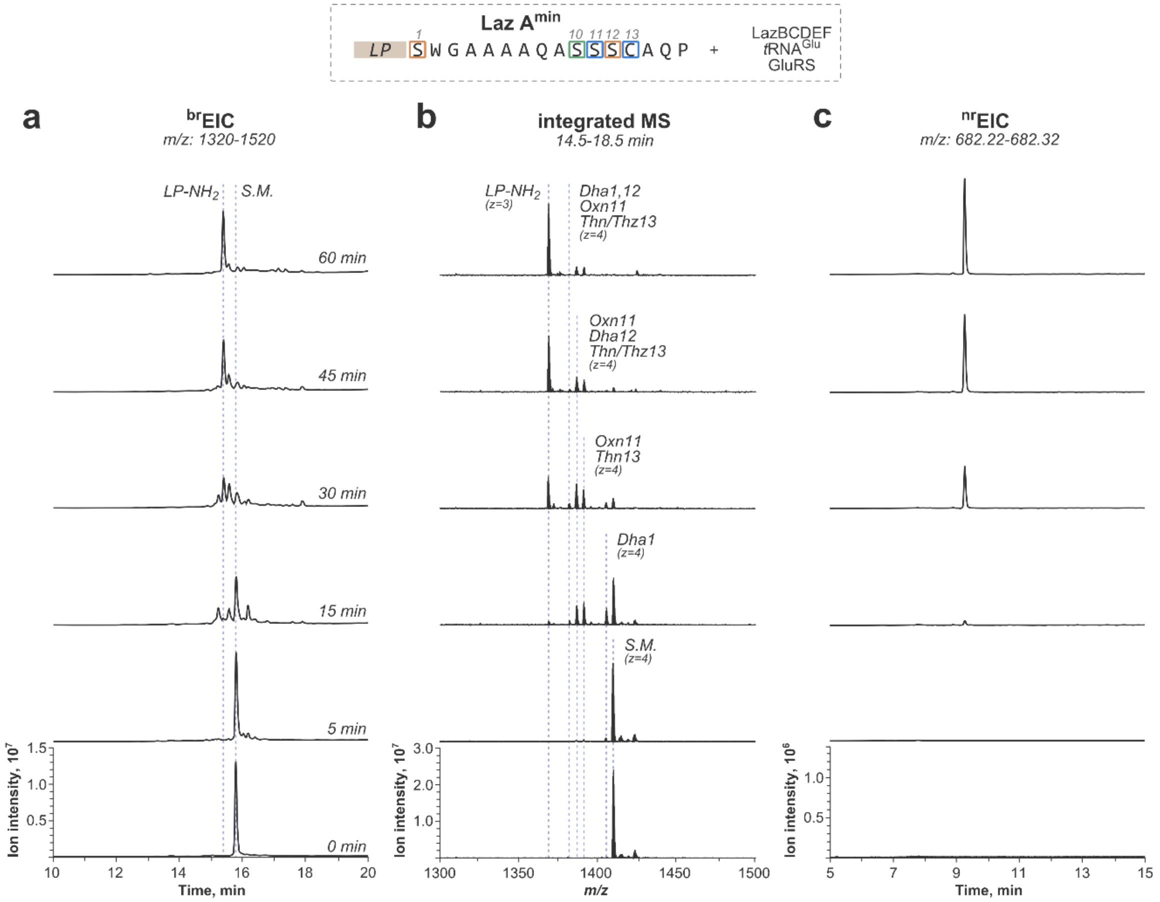

**Figure S7**. Progression of LazA$^{min}$ maturation in the FIT-Laz system. Translation-derived precursor peptide, LazA$^{min}$, was incubated with LazBCDEF/GluRS/tRNA$^{Glu}$ for specified time and the outcomes were analyzed by LC-MS. a) Overview $^{br}$EIC chromatograms visualizing accumulation and consumption of linear intermediates as they are modified from LazA$^{min}$ (0 min) to LP-$NH_2$ (60 min). b) Integrated mass spectra corresponding to chromatograms from panel a) with annotations for individual intermediates forming during the process. c) $^{nr}$EIC chromatograms for the thiopeptide (lactazole S4-C7A) accumulating alongside LP-$NH_2$. Y-axes are scaled internally for each panel.

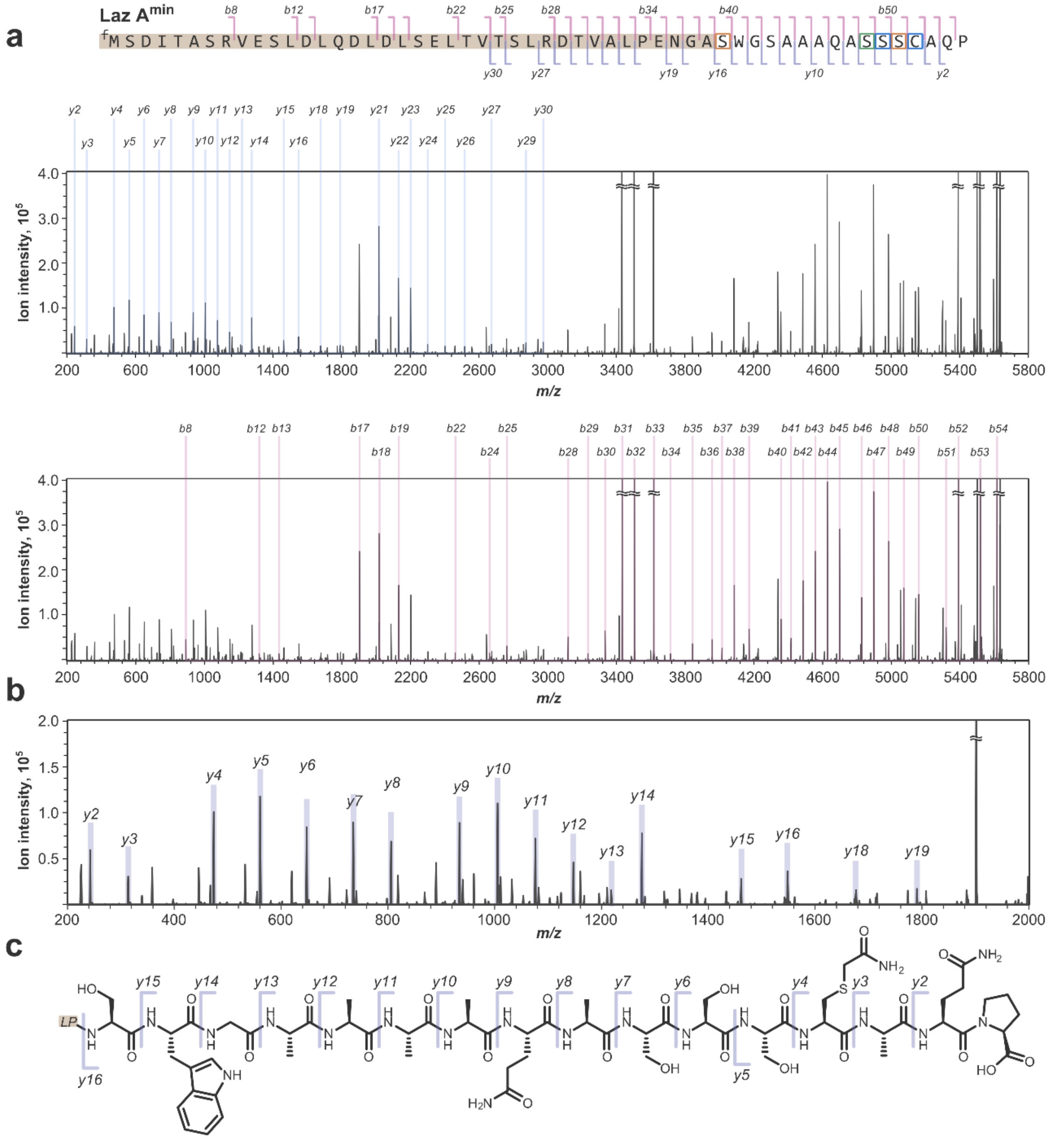

**Figure S8**. Annotated MS/MS spectrum for unmodified LazA$^{min}$. a) Charge-deconvoluted CID fragmentation spectrum for the $z$=4 precursor ion ($m/z$ 1409.18; mono$_{calc}$: 5632.68 Da; mono$_{obs}$: 5632.68 Da) obtained with collision energies ramped from 20.8 to 24.8 eV and displaying *y*- and *b*-ion assignments; stable molecule losses ($H_2O$, $NH_3$, CO, etc.) and double fragmentation assignments are omitted for clarity. "fM" stands for formyl-methionine. b) Zoomed-in low molecular weight ($m/z$ 200-2000) section of the spectrum showing *y*-ion fragmentations inside the CP sequence; full *y*-ion ladder is observed. Stable molecule losses and *b*-ions and are omitted for clarity. c) Chemical structure of LazA$^{min}$ CP with mapped *y*-ion annotations.

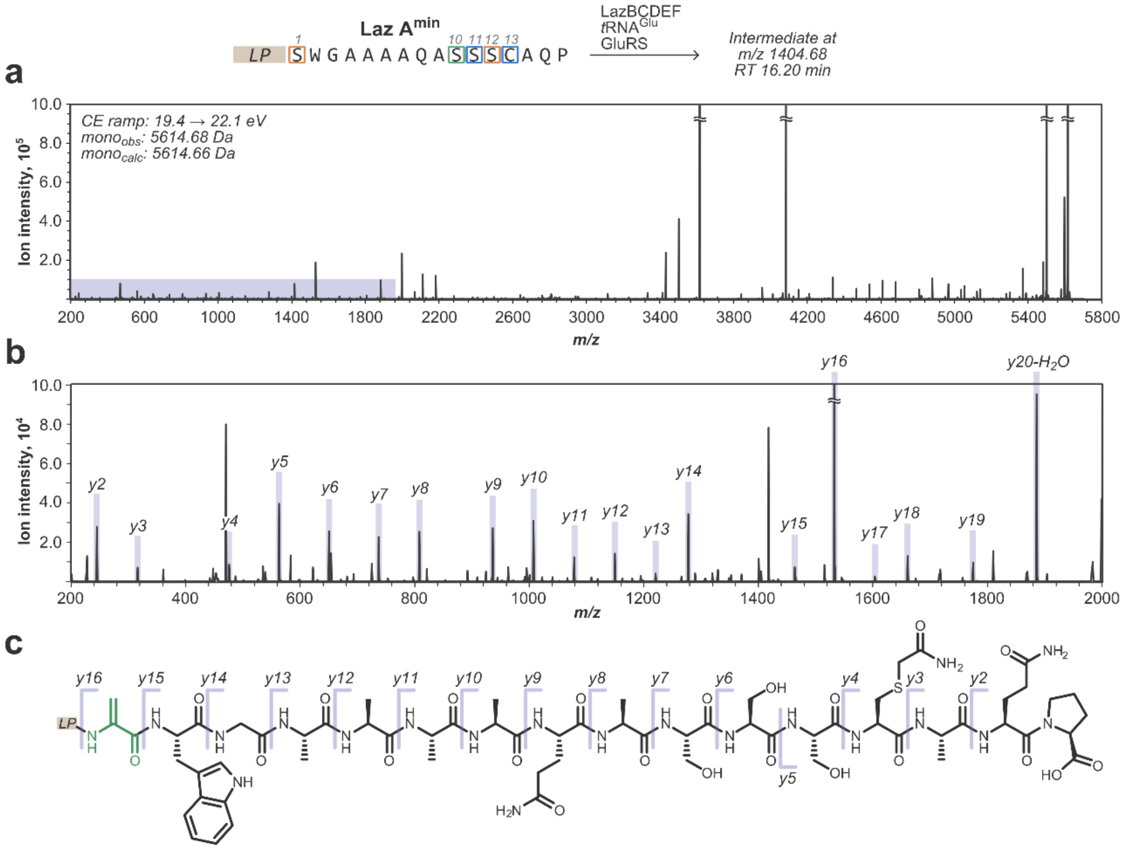

**Figure S9**. Annotated MS/MS spectrum for Dha1 LazA$^{min}$ intermediate captured after treating LazA$^{min}$ with LazBCDEF/GluRS/tRNA$^{Glu}$. a) Full charge-deconvoluted CID fragmentation spectrum for the *z*=4 precursor ion (*m/z* 1404.68; mono$_{calc}$: 5614.66 Da; mono$_{obs}$: 5614.68 Da) obtained with collision energies ramped from 19.4 to 22.1 eV. b) A zoomed-in fraction of the spectrum corresponding to the shaded area from panel a) with spectral assignments; *y*-ions are annotated; most *b*-ions, stable molecule losses ($H_2O$, $NH_3$, CO, etc.) and double fragmentation events are omitted for clarity. c) Chemical structure of Dha1 LazA$^{min}$ CP with mapped *y*-ion annotations. High quality spectrum enabled unambiguous localization of the dehydration event to Ser1.

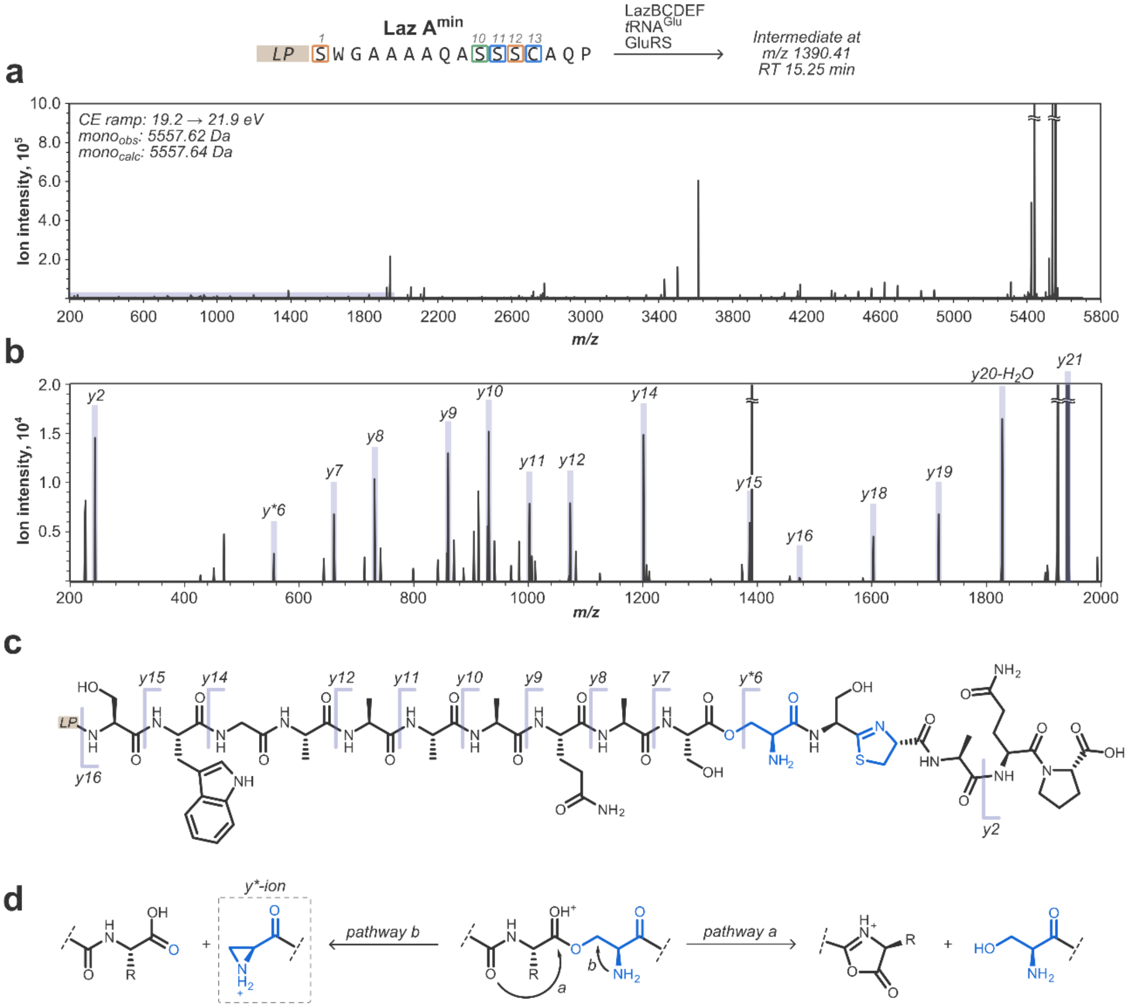

**Figure S10**. Annotated MS/MS spectrum for Oxn11/Thn13 LazA$^{min}$ intermediate captured after treating LazA$^{min}$ with LazBCDEF/GluRS/tRNA$^{Glu}$. a) Full charge-deconvoluted CID fragmentation spectrum for the $z$=4 precursor ion ($m/z$ 1390.41; mono$_{calc}$: 5557.64 Da; mono$_{obs}$: 5557.62 Da) obtained with collision energies ramped from 19.2 to 21.9 eV. b) A zoomed-in fraction of the spectrum corresponding to the shaded area from panel a) with spectral assignments; *y*-ions are annotated; most *b*-ions, stable molecule losses ($H_2O$, $NH_3$, CO, etc.) and double fragmentation events are omitted for clarity. c) Chemical structure of Oxn11/Thn13 LazA$^{min}$ CP with mapped *y*-ion annotations. No IAA-adduct no Cys13 in combination with an almost complete *y*-ion ladder facilitates the assignment as Oxn11/Thn13. Formation of $y^{*}6$ ion support the presence of an Oxn-derived β-aminoester. d) A plausible explanation for the observed fragmentation of Ser11 β-aminoester; β-fragmentations similar to the proposed pathway *b* have been previously reported.[10]

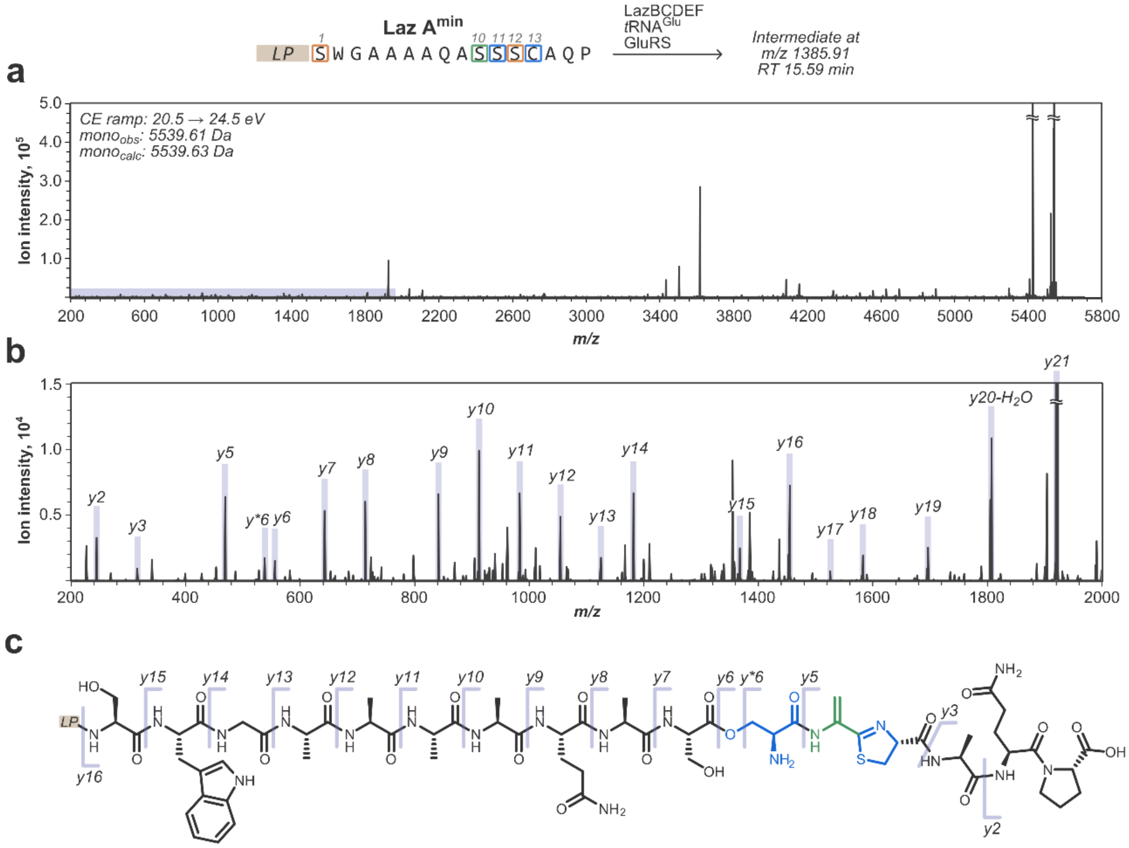

**Figure S11**. Annotated MS/MS spectrum for Oxn11-Dha12-Thn13 LazA$^{min}$ intermediate captured after treating LazA$^{min}$ with LazBCDEF/GluRS/tRNA$^{Glu}$. a) Full charge-deconvoluted CID fragmentation spectrum for the *z*=4 precursor ion (*m/z* 1385.91; mono$_{calc}$: 5539.63 Da; mono$_{obs}$: 5539.61 Da) obtained with collision energies ramped from 20.5 to 24.5 eV. b) A zoomed-in fraction of the spectrum corresponding to the shaded area from panel a) with spectral assignments; *y*-ions are annotated; most *b*-ions, stable molecule losses ($H_2O$, $NH_3$, CO, etc.) and double fragmentation events are omitted for clarity. c) Chemical structure of Oxn11/Thn13 LazA$^{min}$ CP with mapped *y*-ion annotations; *y5* localizes Dha to position12, and supports the assigned Oxn11-Dha12-Thn13 PTM pattern. For notes regarding formation of the *y*$^*$*6* ion, refer to Fig. S10.

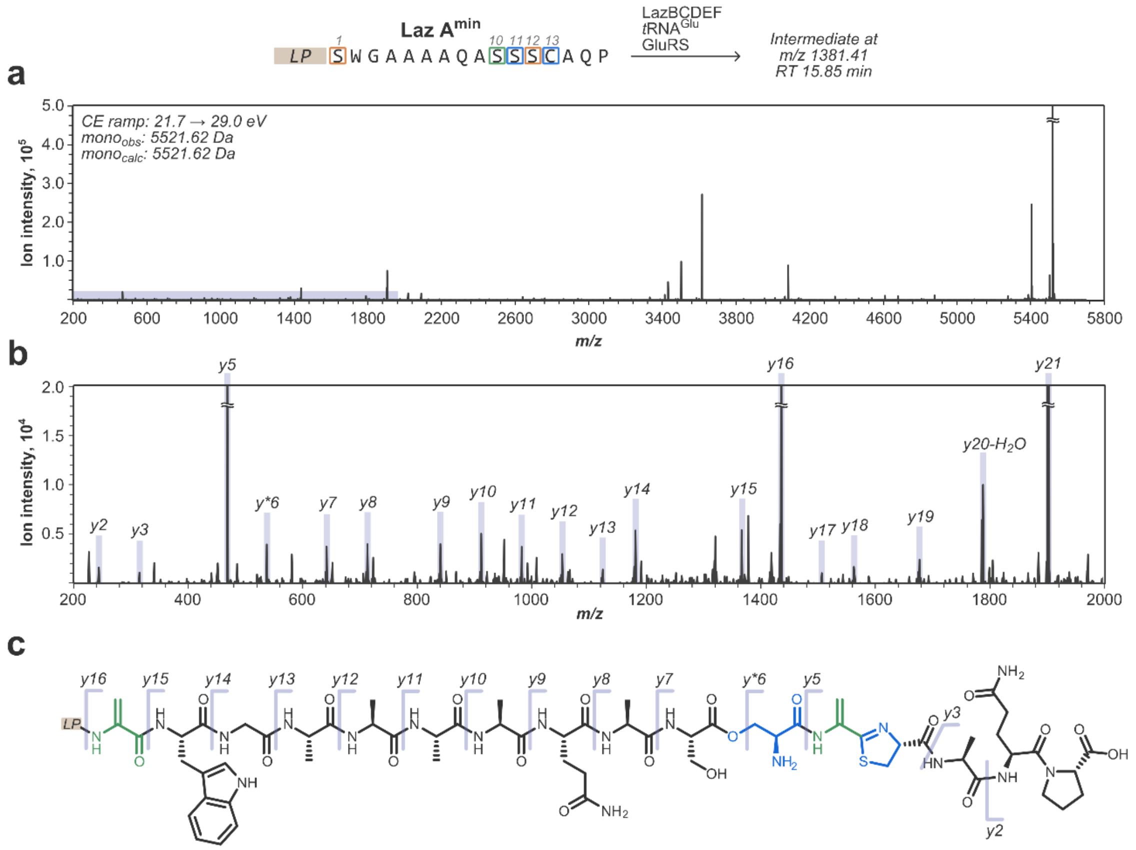

**Figure S12**. Annotated MS/MS spectrum for Dha1/Oxn11-Dha12-Thn13 LazA$^{min}$ intermediate captured after treating LazA$^{min}$ with LazBCDEF/GluRS/tRNA$^{Glu}$. a) Full charge-deconvoluted CID fragmentation spectrum for the *z*=4 precursor ion (*m/z* 1381.41; mono$_{calc}$: 5521.62 Da; mono$_{obs}$: 5521.62 Da) obtained with collision energies ramped from 21.7 to 29.0 eV. b) A zoomed-in fraction of the spectrum corresponding to the shaded area from panel a) with spectral assignments; *y*-ions are annotated; most *b*-ions, stable molecule losses ($H_2O$, $NH_3$, CO, etc.) and double fragmentation events are omitted for clarity. c) Chemical structure of Dha1/Oxn11/Thn13 LazA$^{min}$ CP with mapped *y*-ion annotations; *y5* localizes the first Dha to position 12, and the *y15/16* pair indicates that the second Dha is in position 1. For notes regarding formation of the *y*$^{*}$*6* ion, refer to Fig. S10.

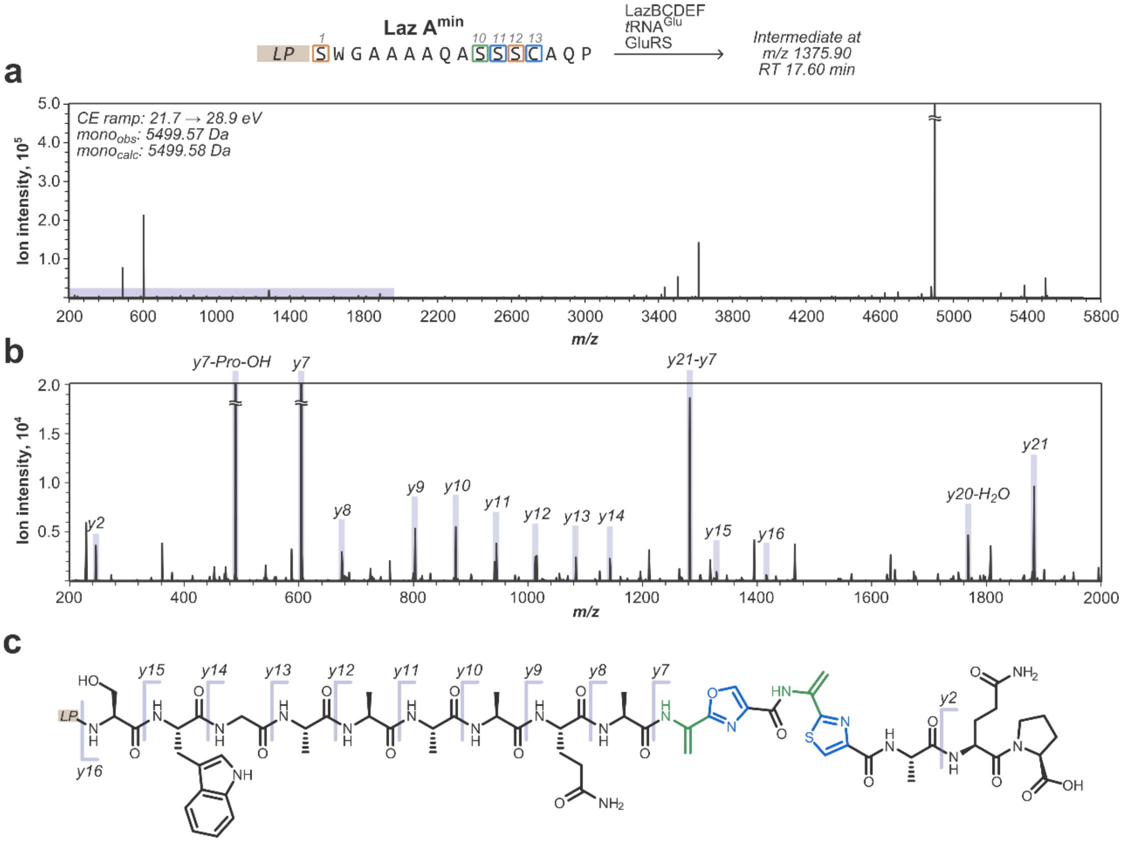

**Figure S13**. Annotated MS/MS spectrum for Dha10-Oxz11-Dha12-Thz13 LazA$^{min}$ intermediate captured after treating LazA$^{min}$ with LazBCDEF/GluRS/tRNA$^{Glu}$. a) Full charge-deconvoluted CID fragmentation spectrum for the *z*=4 precursor ion (*m/z* 1375.90; mono$_{calc}$: 5499.58 Da; mono$_{obs}$: 5499.57 Da) obtained with collision energies ramped from 21.7 to 28.9 eV. b) A zoomed-in fraction of the spectrum corresponding to the shaded area from panel a) with spectral assignments; *y*-ions are annotated; most *b*-ions, stable molecule losses ($H_2O$, $NH_3$, CO, etc.) and double fragmentation events are omitted for clarity. c) Chemical structure of Dha10-Oxz11-Dha12-Thz13 LazA$^{min}$ CP with mapped *y*-ion annotations; a prominent and characteristic *y7* ion supports the Dha10-Oxz11-Dha12-Thz13 assignment.

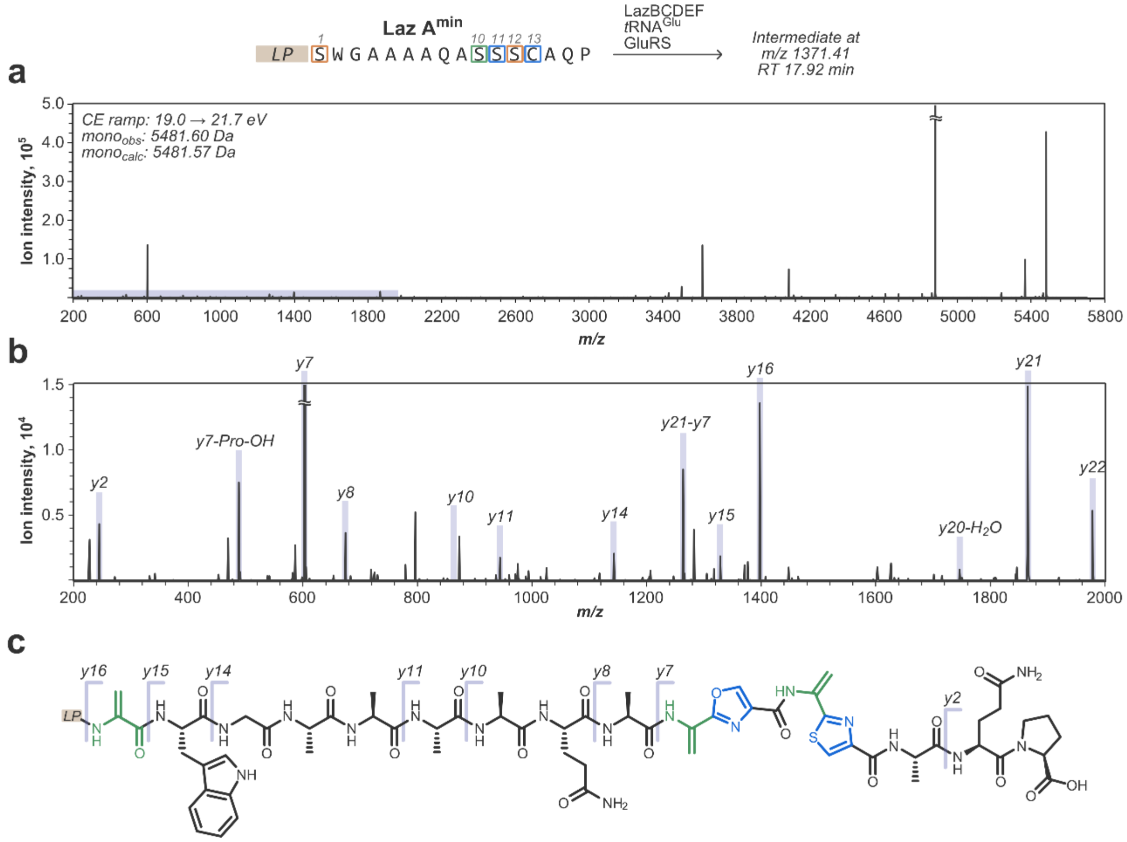

**Figure S14**. Annotated MS/MS spectrum for the macrocyclization precursor peptide, i.e. Dha1/Dha10-Oxz11-Dha12-Thz13 LazA$^{min}$ intermediate, captured after treating LazA$^{min}$ with LazBCDEF/GluRS/tRNA$^{Glu}$. a) Full charge-deconvoluted CID fragmentation spectrum for the $z$=4 precursor ion ($m/z$ 1371.41; mono$_{calc}$: 5481.57 Da; mono$_{obs}$: 5481.60 Da) obtained with collision energies ramped from 19.0 to 21.7 eV. b) A zoomed-in fraction of the spectrum corresponding to the shaded area from panel a) with spectral assignments; *y*-ions are annotated; most *b*-ions, most stable molecule losses ($H_2O$, $NH_3$, CO, etc.) and double fragmentation events are omitted for clarity. c) Chemical structure of Dha1/Dha10-Oxz11-Dha12-Thz13 LazA$^{min}$ CP with mapped *y*-ion annotations; a prominent and characteristic *y7* ion supports the Dha10-Oxz11-Dha12-Thz13 assignment, and the *y15/16* pair indicates the presence of Dha in position 1.

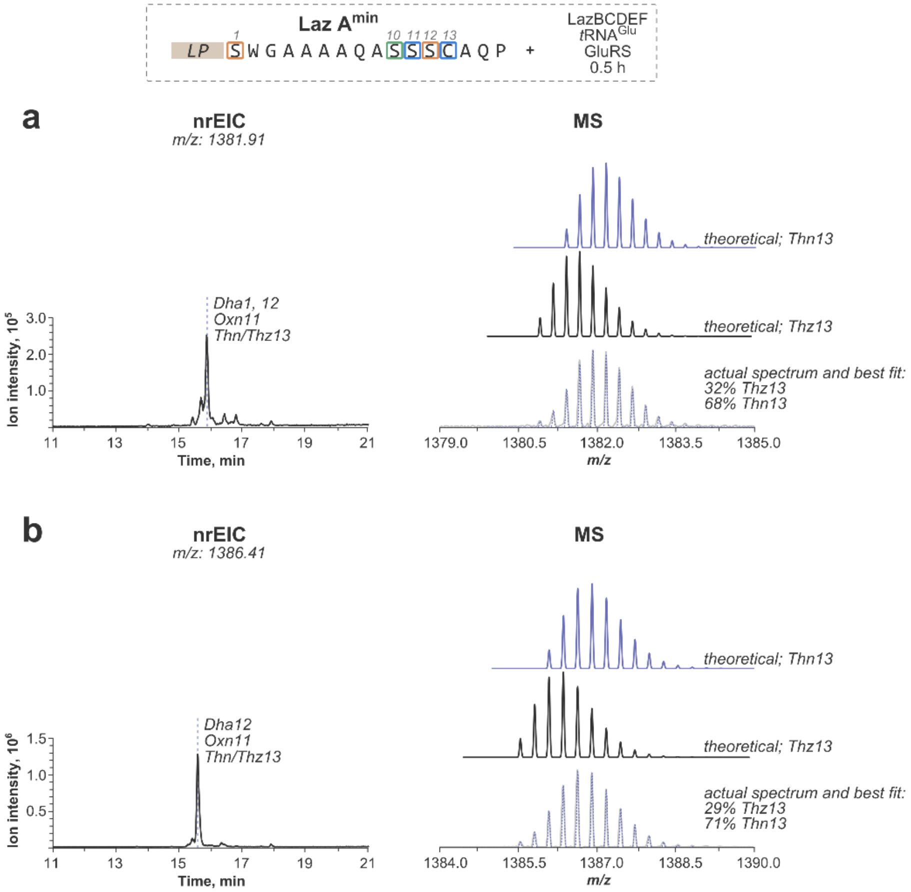

**Figure S15**. Isotope envelope deconvolution enables quantification of Thn/Thz ratio for some biosynthetic intermediates. Certain peptides, for instance, Dha1/Oxn11-Dha12-Thn13 LazA$^{min}$ and Dha1/Oxn11-Dha12-Thz13 LazA$^{min}$, co-eluted by HPLC. Because molecular weight of these peptides differs by only 2.01 Da, their mass spectra considerably overlap, which precludes their direct quantification. In such cases, isotope envelope deconvolution was performed as described in section 2.5. a) Quantification of the Thn/Thz ratio for Dha1/Oxn11-Dha12-Thn13/Thz13 LazA$^{min}$ intermediate. A $^{nr}$EIC chromatogram corresponding to the intermediate in question and a zoomed-in mass spectrum (z=4; grey solid line) integrated over the entire peak are shown. Theoretical mass spectra for the individual components (Thn13 and Thz13) used for fitting as well as the best fit (blue dotted line) are also displayed. After a 30-min incubation with LazBCDEF/GluRS/tRNA$^{Glu}$, Dha1/Oxn11-Dha12-Thn13/Thz13 LazA$^{min}$ intermediate contains approximately 32%Thz. b) Analogously to panel a), quantification of the Thn/Thz ratio for Oxn11-Dha12-Thn13/Thz13 LazA$^{min}$ intermediate.

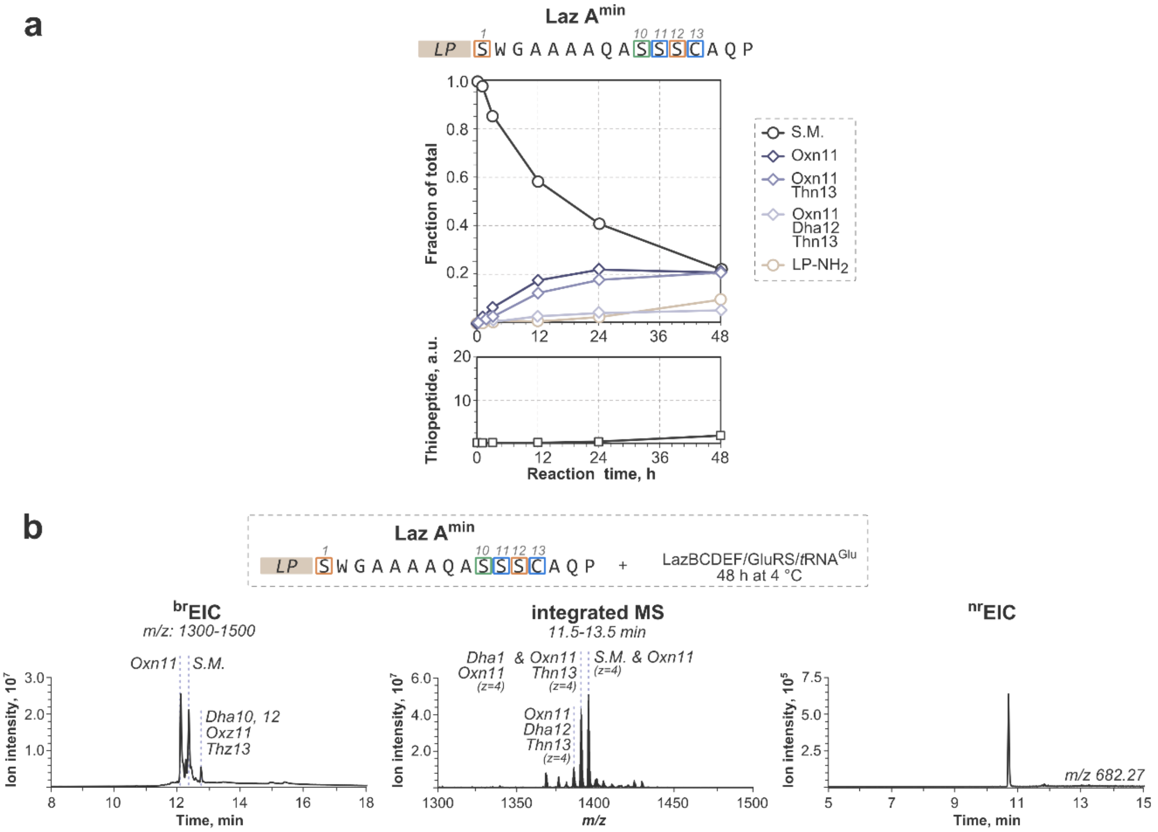

**Figure S16**. Treatment of LazA$^{min}$ by Laz enzymes at 4 °C slowly accumulates early stage intermediates. a) Quantification of the time course experiment. LazA$^{min}$ was treated with LazBCDEF/GluRS/tRNA$^{Glu}$ at 4 °C for 0.5, 1, 3, 12, 24 and 48 h, after which the reaction was quenched and analyzed by LC-MS. PTM assignments and quantification of intermediates were done as described in sections 3.1 and 2.5, respectively. After 48 h, unmodified LazA$^{min}$ remains alongside early stage intermediates (primarily, Oxn11, Oxn11/Thn13 and Oxn11-Dha12-Thn13) and small amounts of LP-NH$_2$ and the thiopeptide. b) LC-MS chromatograms ($^{br}$EIC to visualize linear LazA forms, and $^{nr}$EIC for the produced thiopeptide) and integrated mass spectrum showing the product distribution after the final, 48-h time point.

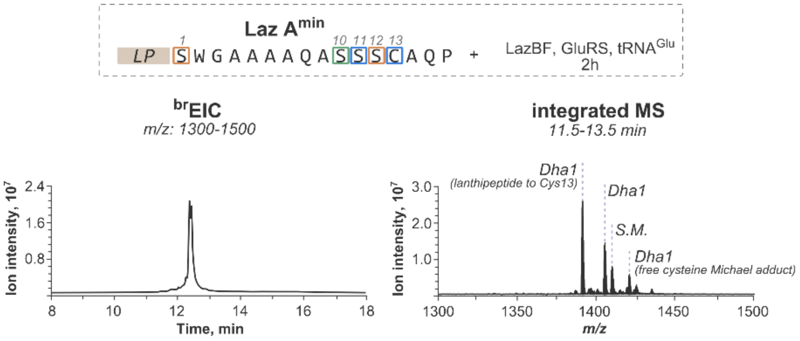

**Figure S17**. LazBF-catalyzed formation of Dha1 in LazA$^{min}$ is independent of azole formation and is kinetically competent. LazA$^{min}$ substrate was accessed with the FIT system, incubated with LazBF/GluRS/tRNA$^{Glu}$ for 2 h, and analyzed by LC-MS. Displayed are a $^{br}$EIC chromatogram and an MS spectrum integrated over the entire peak. LazA$^{min}$ is almost completely consumed after 2 h, and no more than a single dehydration takes place. Several products form due to non-enzymatic Michael addition of various thiols to Dha1.

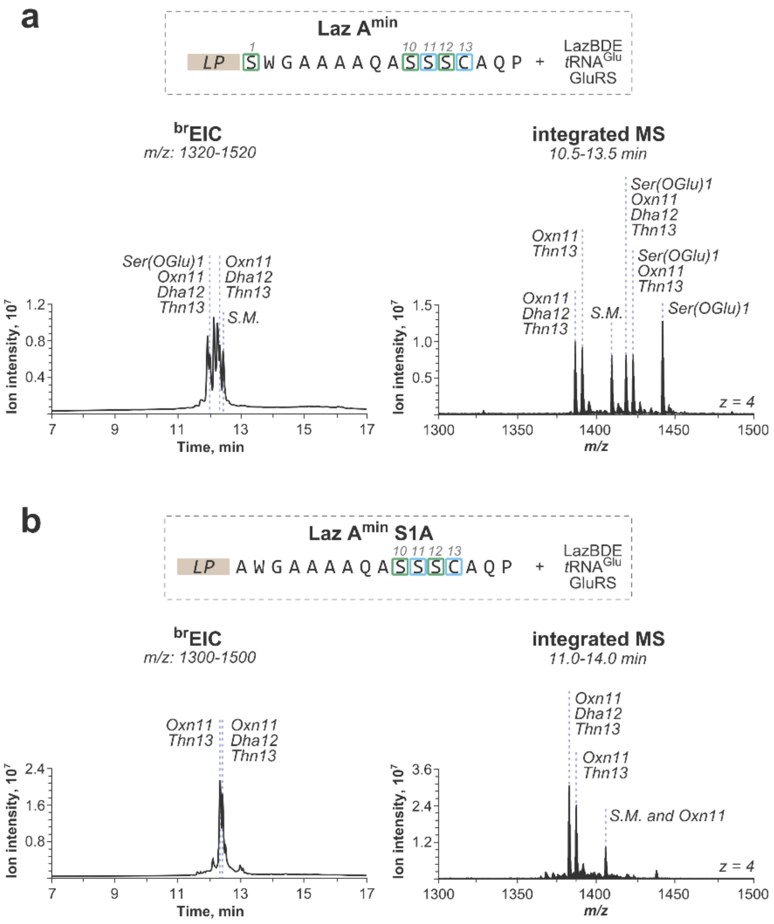

**Figure S18**. LazB can install Dha in an azoline-dependent manner. LazA variants (LazA$^{min}$ in panel a) and LazA$^{min}$ S1A in panel b)) were incubated with the enzyme mix lacking the dehydrogenase activity (LazBDE/GluRS/tRNA$^{Glu}$) for 1 h, and analyzed by LC-MS. Displayed are $^{br}$EIC chromatograms showing the overall product distributions and MS spectra integrated over product peaks. In LazA$^{min}$ case, a complex mixture of products formed. To simplify the annotation, the S1A mutation was introduced, and in that case, after 1 h the Oxn11-Dha12-Thn13 product formed, indicating that LazB can modify Ser adjacent to azolines. Curiously, Dha (as opposed to Ser(OGlu)) formed even in the absence of LazF for Ser12, but not for Ser1 in LazA$^{min}$, suggesting that glutamate elimination from Ser(OGlu) adjacent to a Thn can occur spontaneously.

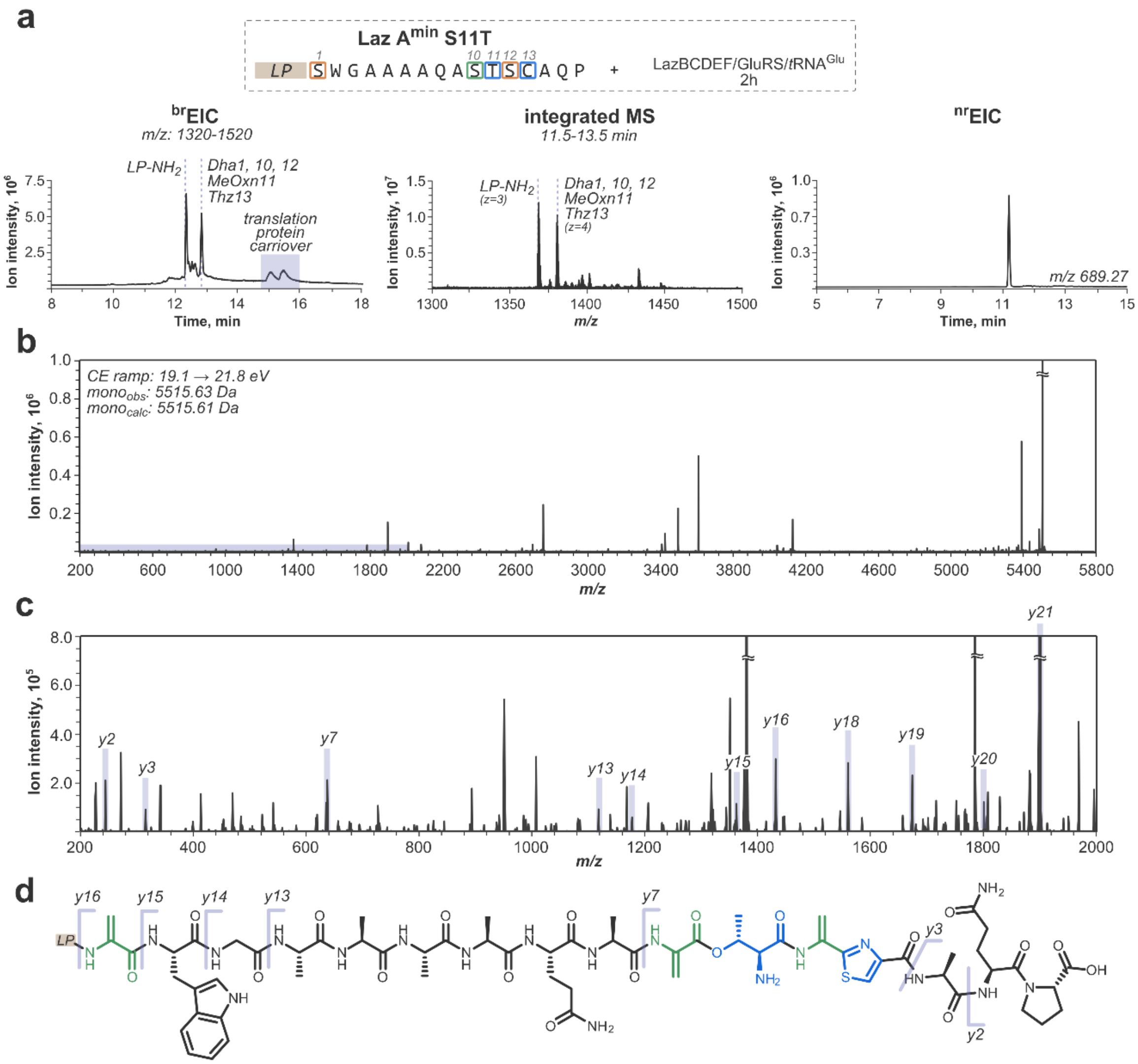

**Figure S19**. Observation of the elusive Dha10-Oxn11-Dha12-Thz13 intermediate. This peptide accumulated together with the thiopeptide/LP-$NH_2$ pair when LazA$^{min}$ S11T mutant was treated LazBCDEF/GluRS/tRNA$^{Glu}$ for 2 h. a) LC-MS chromatograms ($^{br}$EIC to visualize linear LazA forms, and $^{nr}$EIC for the produced thiopeptide) and integrated mass spectrum demonstrating the accumulation of the Dha10-Oxn11-Dha12-Thz13 intermediate. b) Full charge-deconvoluted CID fragmentation spectrum for the *z*=4 precursor ion (*m/z* 1379.92; mono$_{calc}$: 5515.63 Da; mono$_{obs}$: 5515.61 Da) obtained with collision energies ramped from 19.1 to 21.8 eV. c) A zoomed-in fraction of the spectrum corresponding to the shaded area from panel a) with spectral assignments; *y*-ions are annotated; most *b*-ions, stable molecule losses ($H_2O$, $NH_3$, CO, etc.) and double fragmentation events are omitted for clarity. d) Chemical structure of Dha1/Dha10-MeOxn11-Dha12-Thz13 LazA$^{min}$ CP with mapped *y*-ion annotations; the *y3*/*y7* and the *y15/y16* ion pairs support the assignment.

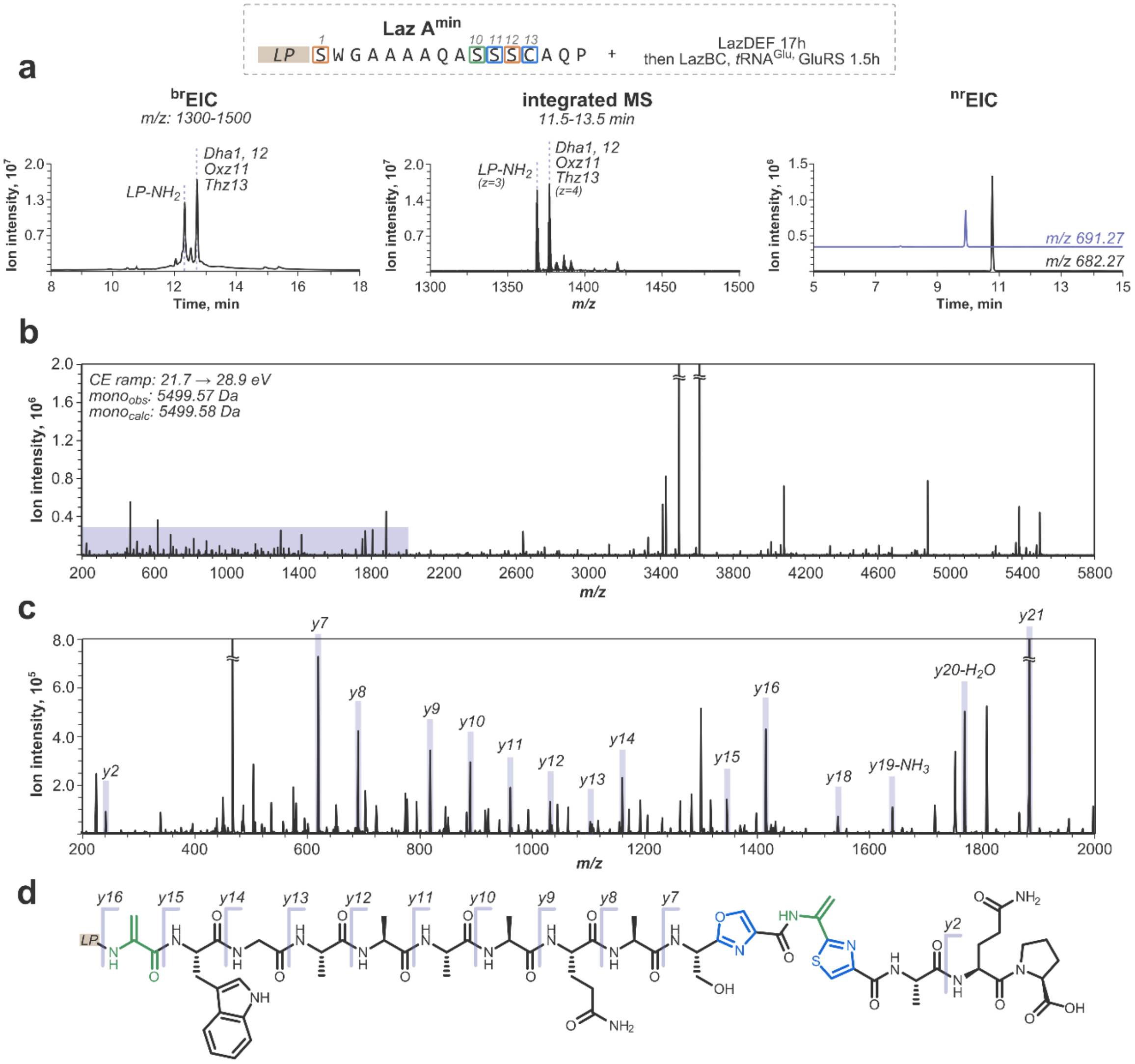

**Figure S20**. Premature oxidation of Oxn11 slows down biosynthesis and leads to a mixture of thiopeptides. To form Oxz11 prior to Dha installation, *in vitro* translated LazA$^{min}$ precursor peptide was incubated with LazDEF for 17 h, which cleanly yielded the Oxz11/Thz13 intermediate (refer to Fig. 3). Treating this product with LazBC/GluRS/tRNA$^{Glu}$ for 1.5 h resulted in the Dha1/Oxz11-Dha12-Thz13 peptide, indicating that Ser10 dehydration is slow when Oxn11 is prematurely oxidized. Additionally, a mixture of thiopeptides formed. Together, these data point to the critical role of LazF during lactazole assembly. a) LC-MS chromatograms ($^{br}$EIC to visualize linear LazA forms, and $^{nr}$EIC for the produced thiopeptides) and integrated mass spectrum demonstrating accumulation of the Dha1/Oxz11-Dha12-Thz13 intermediate. The second thiopeptide (*m/z* 691.27) corresponds to the underdehydrated product, i.e. Ser10-lactazole S4-C7A. b) Full charge-deconvoluted CID fragmentation spectrum for the *z*=4 precursor ion (*m/z* 1375.90; mono$_{calc}$: 5499.58 Da; mono$_{obs}$: 5499.57 Da) obtained with collision energies ramped from 21.7 to 28.9 eV. c) A zoomed-in fraction of the spectrum corresponding to the shaded area from panel a) with spectral assignments; *y*-ions are annotated; most *b*-ions, stable molecule losses ($H_2O$, $NH_3$, CO, etc.) and double fragmentation events are omitted for clarity. d) Chemical structure of Dha1/Oxz11-Dha12-Thz13 LazA$^{min}$ CP with mapped *y*-ion annotations; the *y2*/*y7* and the *y15/y16* ion pairs support the assignment.

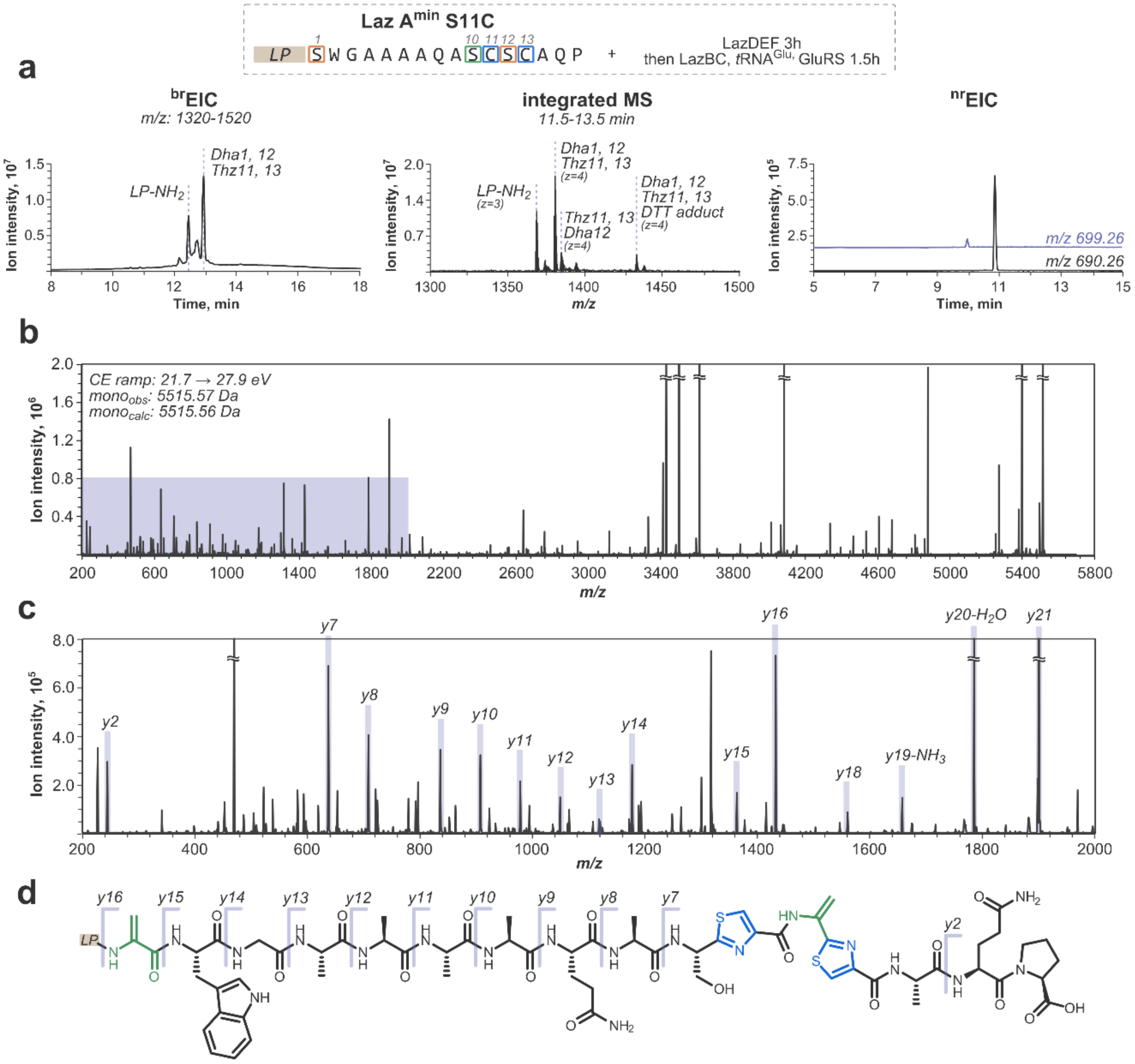

**Figure S21**. Similar to Fig. S20, maturation of LazA$^{min}$ S11C is slow. *In vitro* translated LazA$^{min}$ S11C was incubated with LazDEF for 3 h and then with LazBC/GluRS/tRNA$^{Glu}$ for 1.5 h, after which the Dha1/Thz11-Dha12-Thz13 peptide accumulated. Incubating LazA$^{min}$ S11C with the full enzyme set led to a similar outcome, except the key intermediate was accompanied by a number of lanthipeptide shunt products (data not shown). a) LC-MS chromatograms ($^{br}$EIC to visualize linear LazA forms, and $^{nr}$EIC for the produced thiopeptides) and integrated mass spectrum demonstrating the accumulation of the Dha1/Thz11-Dha12-Thz13 intermediate. The second thiopeptide (*m/z* 699.26) corresponds to the underdehydrated product, i.e. Ser10 lactazole S4-C7A/S11C. b) Full charge-deconvoluted CID fragmentation spectrum for the *z*=4 precursor ion (*m/z* 1379.90; mono$_{calc}$: 5515.56 Da; mono$_{obs}$: 5515.57 Da) obtained with collision energies ramped from 21.7 to 27.9 eV. c) A zoomed-in fraction of the spectrum corresponding to the shaded area from panel a) with spectral assignments; *y*-ions are annotated; most *b*-ions, stable molecule losses ($H_2O$, $NH_3$, CO, etc.) and double fragmentation events are omitted for clarity. d) Chemical structure of Dha1/Thz11-Dha12-Thz13 LazA$^{min}$ S11C CP with mapped *y*-ion annotations; the *y2*/*y7* and the *y15/y16* ion pairs support the assignment.

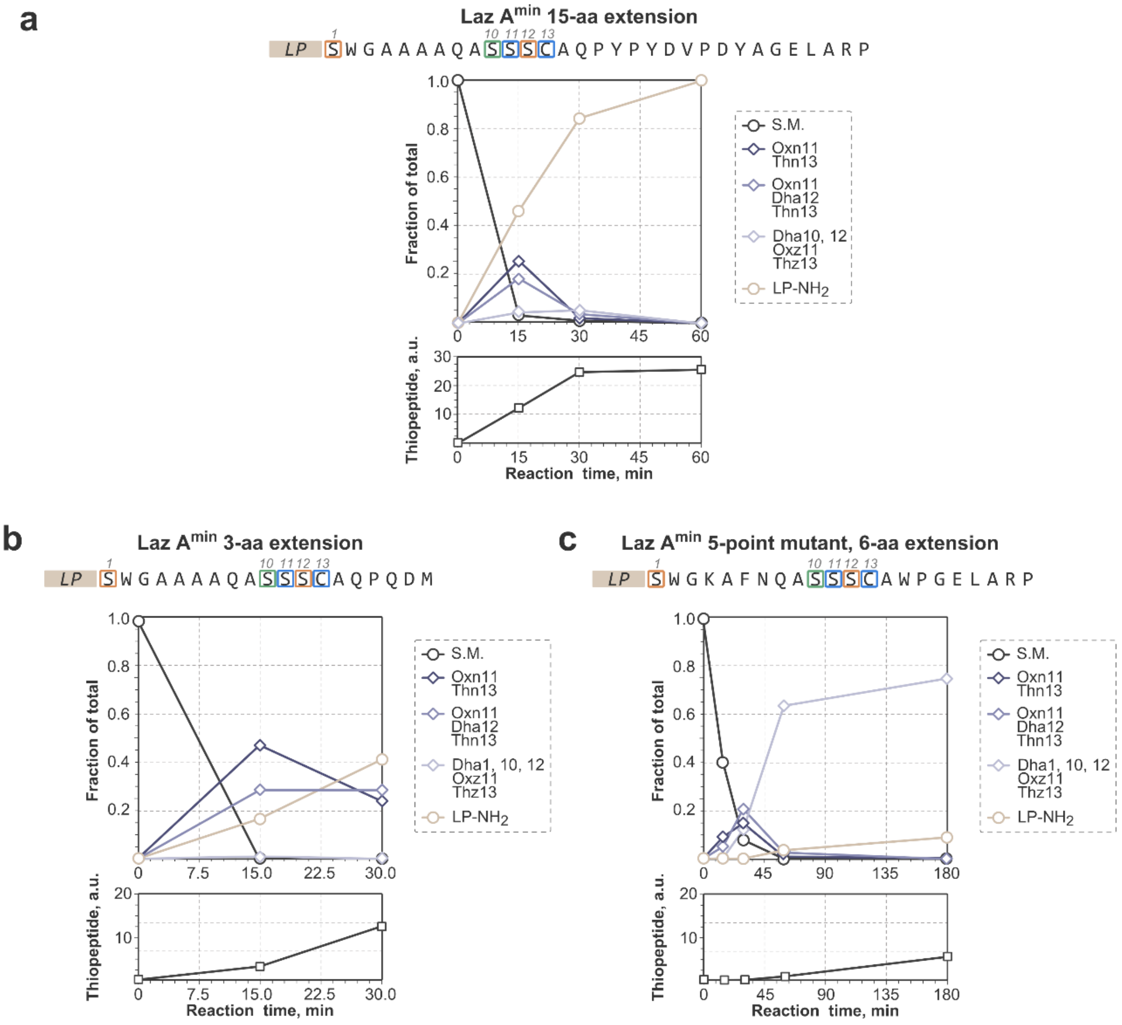

**Figure S22**. Maturation of LazA$^{min}$ variants proceeds via the same pathway as LazA$^{min}$. a) Time course analysis of LazA$^{min}$ 15-aa extension peptide maturation. Precursor peptide was treated with LazBCDEF/GluRS/tRNA$^{Glu}$ for 15, 30 and 60 min, and the outcomes were analyzed by LC-MS. PTM assignments and quantification of intermediates were done as described in sections 3.1 and 2.5, respectively. In this case, biosynthesis proceeds both faster and more efficiently than LazA$^{min}$, but the observed intermediates are the same. b) Time course analysis of LazA$^{min}$ 3-aa extension peptide maturation as in panel a), except the 60-min time point was not acquired. For this peptide, biosynthesis follows the LazA$^{min}$ pathway at a comparable rate. c) Time course analysis of LazA$^{min}$ 5-point mutant/6-aa extension peptide maturation as in panel a), except the time course was extended to 3 h. Maturation of this substrate proceeds substantially slower than LazA$^{min}$ due to accumulation of fully modified linear precursor peptide (Dha1/Dha10-Oxz11-Dha12-Thz13), which indicates that LazC-catalyzed macrocyclization is the rate limiting step for this mutant.

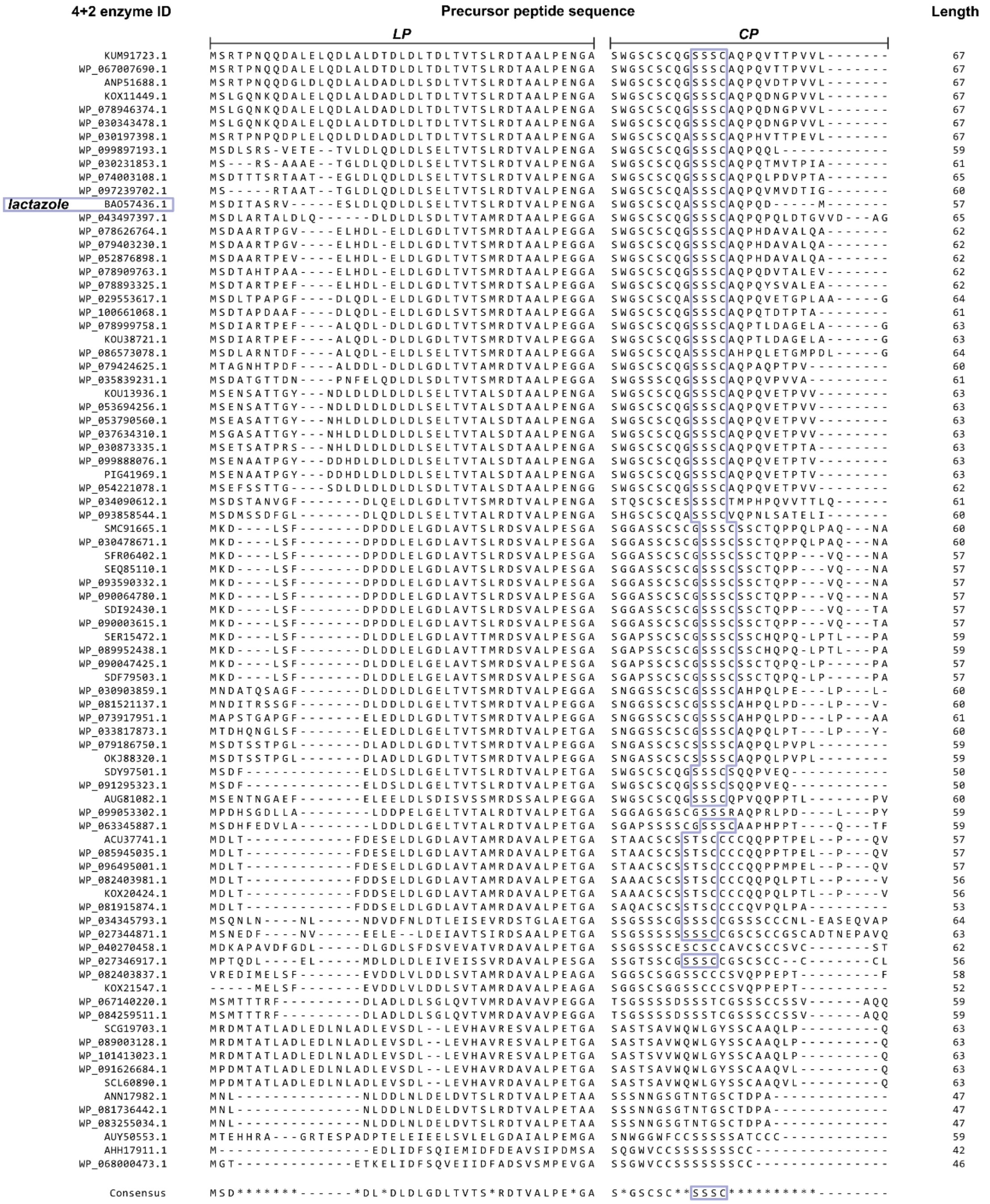

**Figure S23**. Sequence alignment of predicted lactazole-like thiopeptide precursor peptides. Based on the sequence similarity between structural genes, 251 lactazole-like BGCs[11] were narrowed down to 83 members closest to *laz* BGC. CP sequences of 60 of these precursor peptides include a Ser-Ser-Ser-Cys motif similar to lactazole, and additional 6 peptides have a Ser-Thr-Ser-Cys motif. Sequence alignment was done in CLC Genomics Workbench 12.0 (Qiagen Bioinformatics) with the following parameters: gap open cost = 30; gap extension cost = 3.

#### 3.4. LazA$^{aux}$ time course assignments

Assignments of PTM patterns on captured intermediates were done following the scheme outlined in section 3.1 and from MS/MS data. See also section 3.3 for assignment of shunt products. Table S2 summarizes observed intermediates and shunt products. Figures S24-31 show data to support these annotations.

**Table S2**. Progress of LazA$^{aux}$ biosynthesis in the FIT-Laz system. For each intermediate, *m/z* values correspond to the monoisotopic mass peaks. RT stands for HPLC retention time (method B).

| ***m/z*** | **RT, min** | **assignment** | **notes** | **MS/MS confirmation** | **reaction time, min** | | | | |
|---|---|---|---|---|---|---|---|---|---|
| | | | | | **0** | **5** | **15** | **30** | **60** |
| 1375.39 | 17.94 | Dha1, 4, 6<br>Thz5, 7 | Fully modified peptide | | 0.0 | 0.0 | 0.3 | 0.9 | 3.7 |
| 1432.65 | 16.62 | Dha1, 6<br>Thz5, 7<br>DTT-adduct | Lactazole A modification pattern | | 0.0 | 0.0 | 0.0 | 5.8 | 19.3 |
| 1379.90 | 17.35 | Dha1, 6<br>Thz5, 7 | | Fig. S31 | 0.0 | 5.8 | 40.2 | 55.4 | 63.4 |
| 1384.40 | 17.01 | Thz5, 7<br>Dha6 | | Fig. S30 | 0.0 | 54.0 | 47.2 | 28.1 | 5.3 |
| 1389.40 | 15.97 | Thz5<br>Thn7 | | Fig. S29 | 0.0 | 7.8 | 0.0 | 0.0 | 0.0 |
| 1404.15 | 16.39 | Thn5<br>Dha6 | | Fig. S28 | 0.0 | 3.0 | 0.2 | 0.1 | 0.0 |
| 1408.66 | 16.09 | Dha1 to Cys5/7<br>Lanthipeptide | shunt product | | 0.0 | 4.1 | 7.3 | 6.9 | 5.6 |
| 1408.67 | 15.69 | Thn5 | | Fig. S27 | 0.0 | 8.8 | 0.4 | 0.0 | 0.0 |
| 1422.91 | 15.90 | Dha1 | | Fig. S26 | 0.0 | 3.9 | 0.0 | 0.0 | 0.0 |
| 1427.42 | 15.54 | S.M. | | Fig. S25 | 100.0 | 9.1 | 1.9 | 0.7 | 0.5 |
| | | minor products | | | 0.0 | 3.5 | 2.6 | 2.2 | 2.1 |

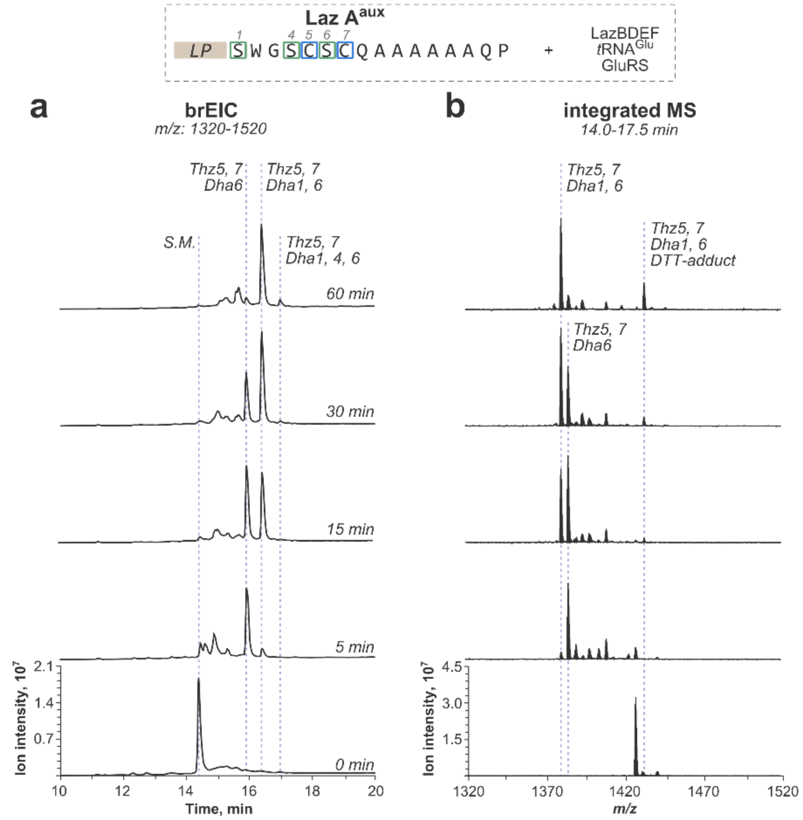

**Figure S24**. Progression of PTM installation for LazA$^{aux}$. Translation-derived substrate, LazA$^{aux}$, was incubated with LazBDEF/GluRS/tRNA$^{Glu}$ for specified time and the outcomes were analyzed by LC-MS. a) Overview $^{br}$EIC chromatograms visualizing accumulation and consumption of linear forms. b) Integrated mass spectra corresponding to chromatograms from panel a) with annotations for individual intermediates forming during the process. Y-axes are scaled internally for each panel.

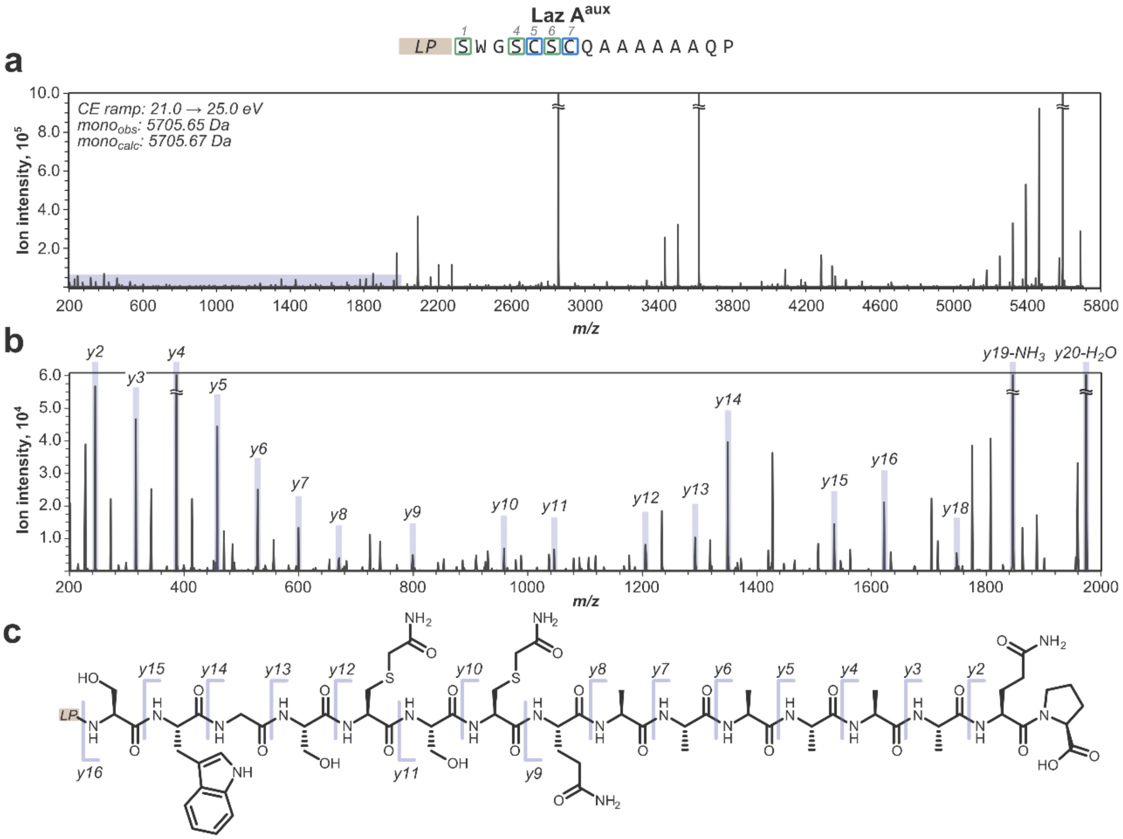

**Figure S25**. Annotated MS/MS spectrum for the unmodified peptide, i.e. LazA$^{aux}$. a) Full charge-deconvoluted CID fragmentation spectrum for the $z$=4 precursor ion ($m/z$ 1427.42; mono$_{calc}$: 5705.67 Da; mono$_{obs}$: 5705.65 Da) obtained with collision energies ramped from 21.0 to 25.0 eV. b) A zoomed-in fraction of the spectrum corresponding to the shaded area from panel a) with spectral assignments; *y*-ions are annotated; most *b*-ions, stable molecule losses ($H_2O$, $NH_3$, CO, etc.) and double fragmentation events are omitted for clarity. c) Chemical structure of LazA$^{aux}$ CP with mapped *y*-ion annotations; full *y*-ion ladder is observed.

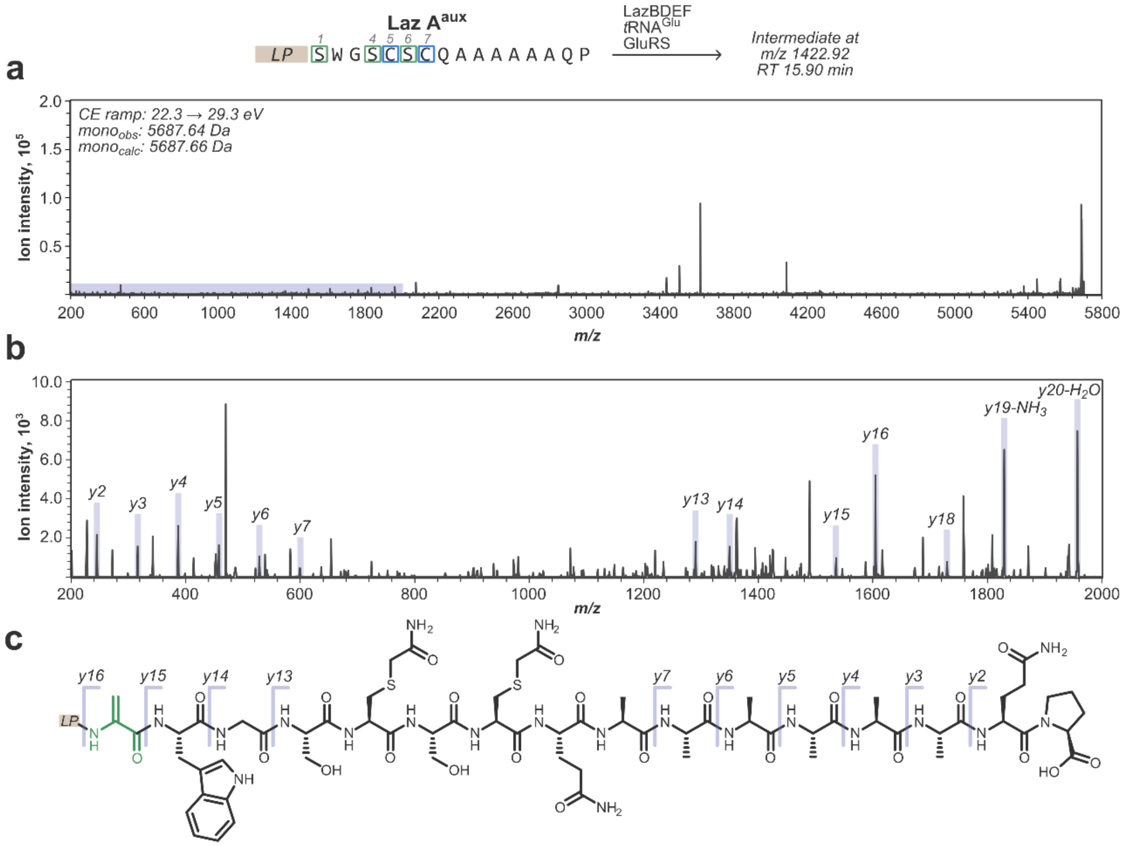

**Figure S26**. Annotated MS/MS spectrum for Dha1 LazA$^{aux}$ intermediate captured after treating LazA$^{aux}$ with LazBDEF/GluRS/tRNA$^{Glu}$. a) Full charge-deconvoluted CID fragmentation spectrum for the $z$=4 precursor ion ($m/z$ 1422.92; mono$_{calc}$: 5687.66 Da; mono$_{obs}$: 5687.64 Da) obtained with collision energies ramped from 22.3 to 29.3 eV. b) A zoomed-in fraction of the spectrum corresponding to the shaded area from panel a) with spectral assignments; *y*-ions are annotated; most *b*-ions, stable molecule losses ($H_2O$, $NH_3$, CO, etc.) and double fragmentation events are omitted for clarity. c) Chemical structure of Dha1 LazA$^{min}$ CP with mapped *y*-ion annotations. The *y15/y16* pair of ions localizes the dehydration event to Ser1.

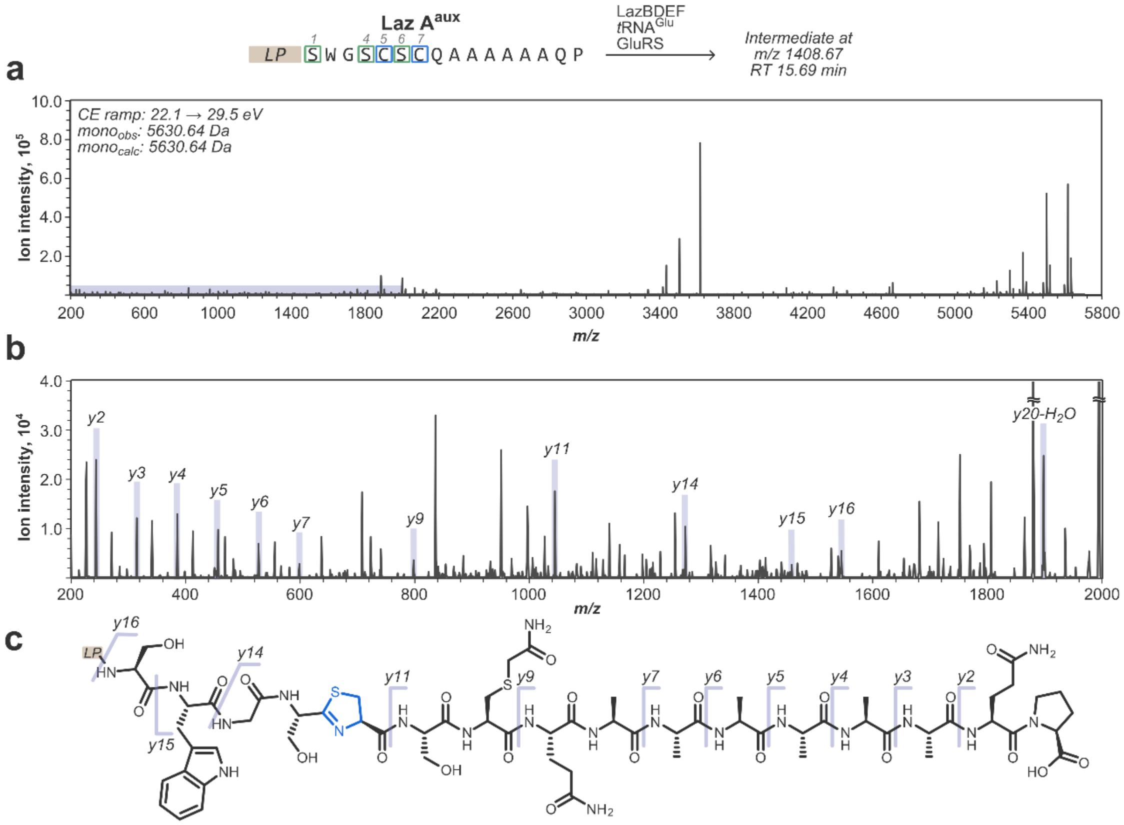

**Figure S27**. Annotated MS/MS spectrum for Thn5 LazA$^{aux}$ intermediate captured after treating LazA$^{aux}$ with LazBDEF/GluRS/tRNA$^{Glu}$. a) Full charge-deconvoluted CID fragmentation spectrum for the *z*=4 precursor ion (*m/z* 1408.67; mono$_{calc}$: 5630.64 Da; mono$_{obs}$: 5630.64 Da) obtained with collision energies ramped from 22.1 to 29.5 eV. b) A zoomed-in fraction of the spectrum corresponding to the shaded area from panel a) with spectral assignments; *y*-ions are annotated; most *b*-ions, stable molecule losses ($H_2O$, $NH_3$, CO, etc.) and double fragmentation events are omitted for clarity. c) Chemical structure of Thn5 LazA$^{min}$ CP with mapped *y*-ion annotations. The *y9/y11* ion pair indicates that Cys7 remains intact, implying that the cyclodehydration event took place at Cys5.

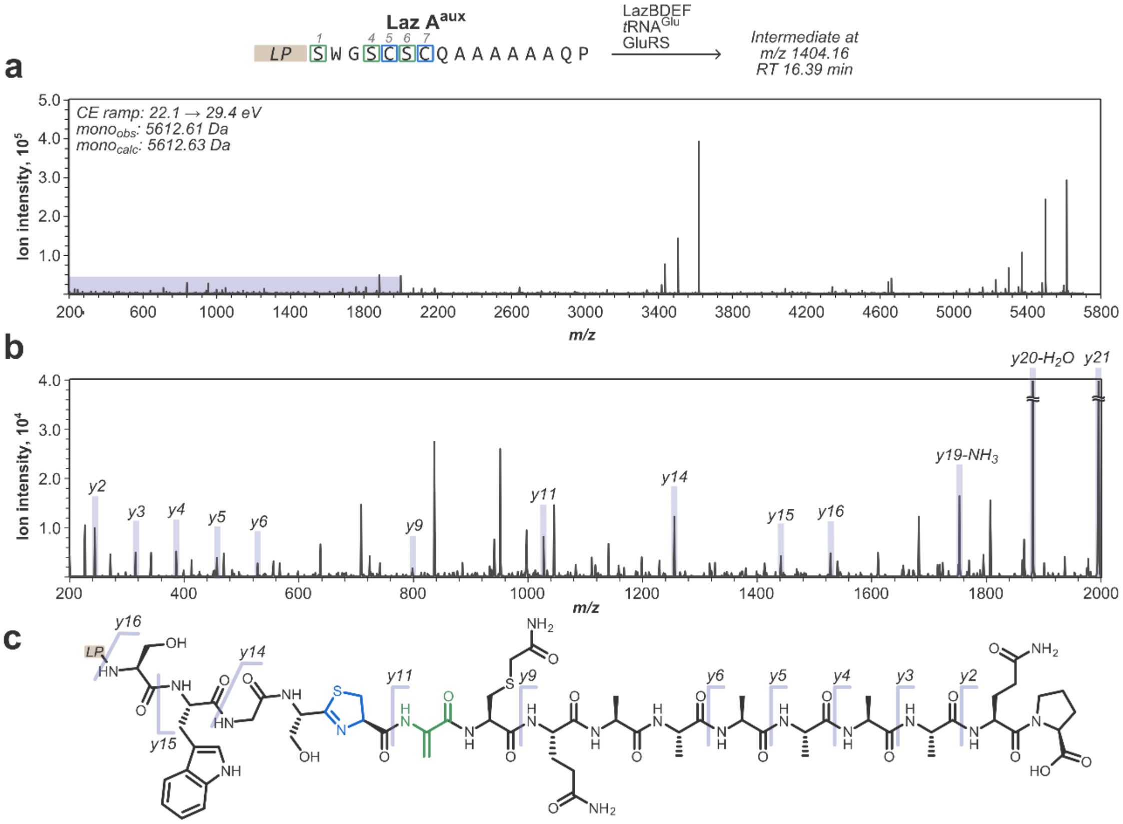

**Figure S28**. Annotated MS/MS spectrum for Thn5-Dha6 LazA$^{aux}$ intermediate captured after treating LazA$^{aux}$ with LazBDEF/GluRS/tRNA$^{Glu}$. a) Full charge-deconvoluted CID fragmentation spectrum for the $z$=4 precursor ion ($m/z$ 1404.16; mono$_{calc}$: 5612.63 Da; mono$_{obs}$: 5612.61 Da) obtained with collision energies ramped from 22.1 to 29.4 eV. b) A zoomed-in fraction of the spectrum corresponding to the shaded area from panel a) with spectral assignments; *y*-ions are annotated; most *b*-ions, stable molecule losses ($H_2O$, $NH_3$, CO, etc.) and double fragmentation events are omitted for clarity. c) Chemical structure of Thn5-Dha6 LazA$^{min}$ CP with mapped *y*-ion annotations. The *y9/y11* ion pair indicates that Cys7 remains intact (implying that the cyclodehydration event took place at Cys5), and supports formation of Dha at Ser6. The *y15/y16* ion pair further indicates that Ser1 remains intact.

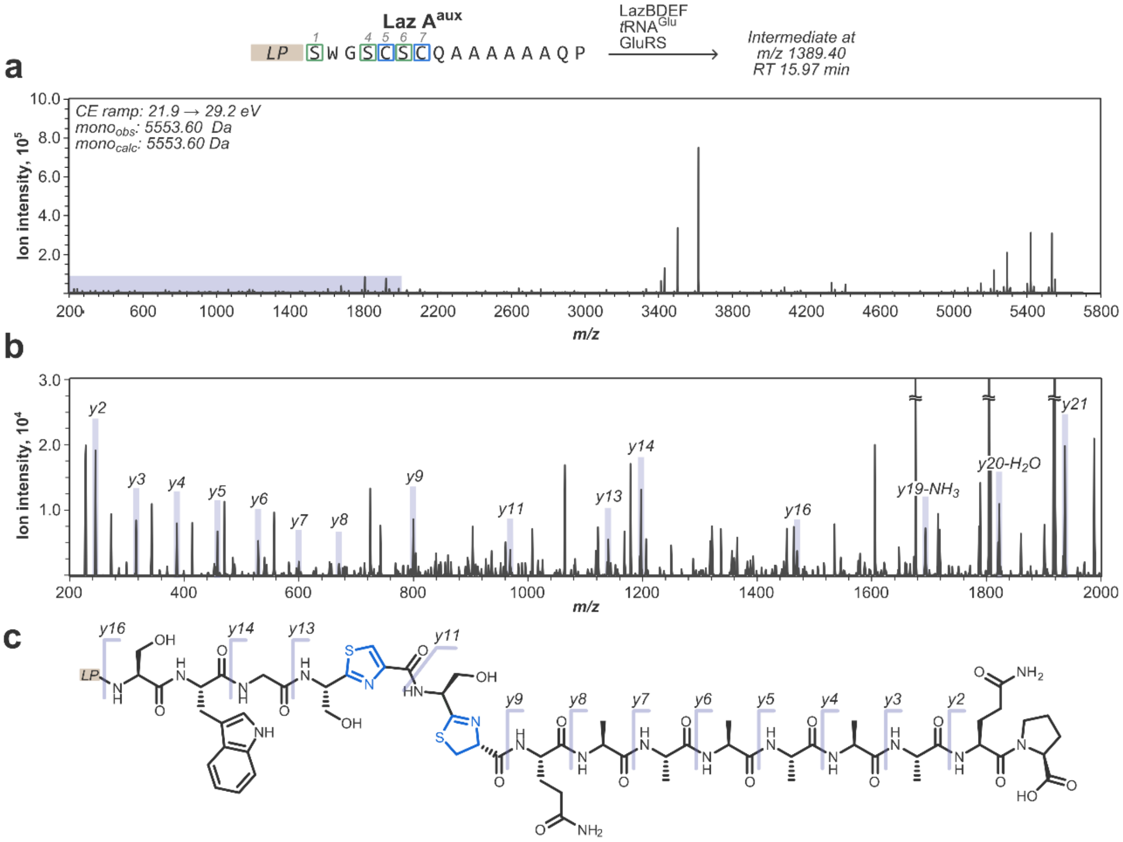

**Figure S29**. Annotated MS/MS spectrum for Thz5/Thn7 LazA$^{aux}$ intermediate captured after treating LazA$^{aux}$ with LazBDEF/GluRS/tRNA$^{Glu}$. a) Full charge-deconvoluted CID fragmentation spectrum for the $z$=4 precursor ion ($m/z$ 1389.40; mono$_{calc}$: 5553.60 Da; mono$_{obs}$: 5553.60 Da) obtained with collision energies ramped from 21.9 to 29.2 eV. b) A zoomed-in fraction of the spectrum corresponding to the shaded area from panel a) with spectral assignments; *y*-ions are annotated; most *b*-ions, stable molecule losses ($H_2O$, $NH_3$, CO, etc.) and double fragmentation events are omitted for clarity. c) Chemical structure of Thz5/Thn7 LazA$^{min}$ CP with mapped *y*-ion annotations. The *y9/y11* ion pair supports formation of Thn at Cys7, and the *y11/y13* pair indicates that Thz formed at Cys5.

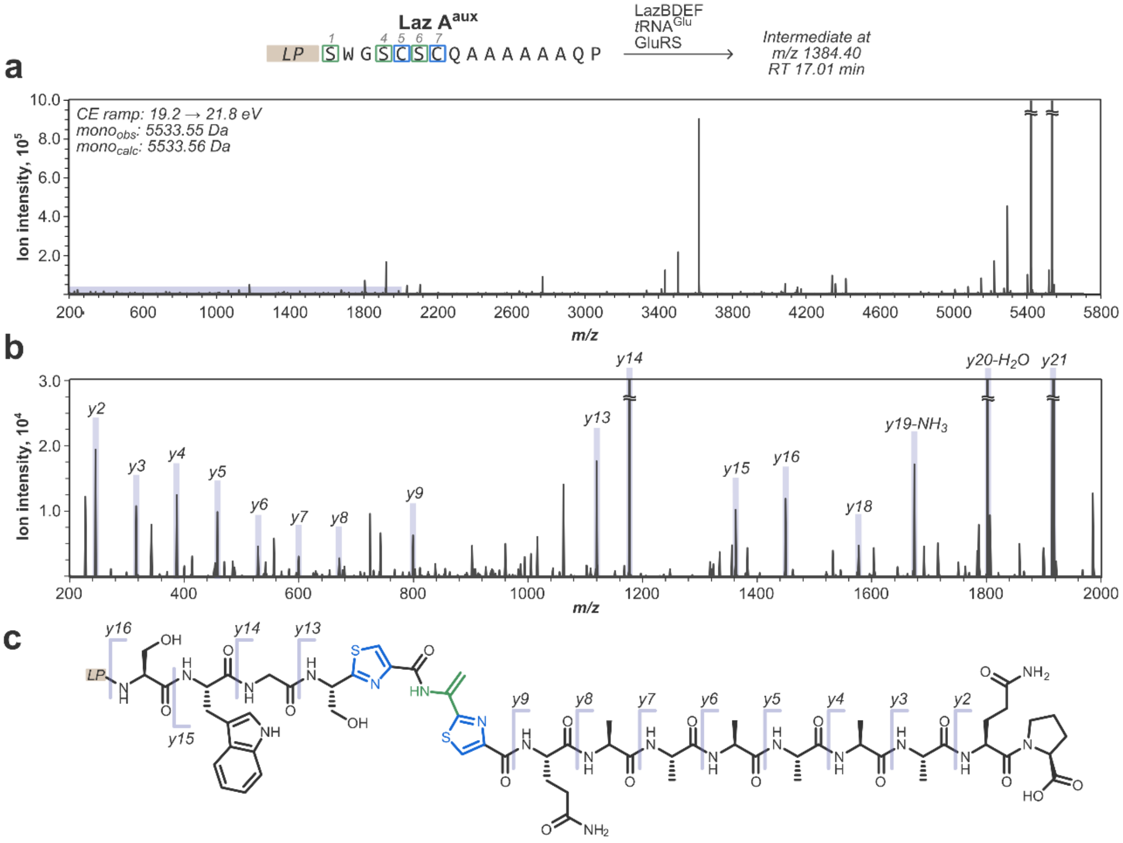

**Figure S30**. Annotated MS/MS spectrum for Thz5-Dha6-Thz7 LazA$^{aux}$ intermediate captured after treating LazA$^{aux}$ with LazBDEF/GluRS/tRNA$^{Glu}$. a) Full charge-deconvoluted CID fragmentation spectrum for the *z*=4 precursor ion (*m/z* 1384.40; mono$_{calc}$: 5533.56 Da; mono$_{obs}$: 5533.55 Da) obtained with collision energies ramped from 19.2 to 21.8 eV. b) A zoomed-in fraction of the spectrum corresponding to the shaded area from panel a) with spectral assignments; *y*-ions are annotated; most *b*-ions, stable molecule losses ($H_2O$, $NH_3$, CO, etc.) and double fragmentation events are omitted for clarity. c) Chemical structure of Thz5-Dha6-Thz7 LazA$^{min}$ CP with mapped *y*-ion annotations. The *y9/y13* ion pair supports the assignment, and the *y15/y16* additionally indicates that Ser1 remains unmodified.

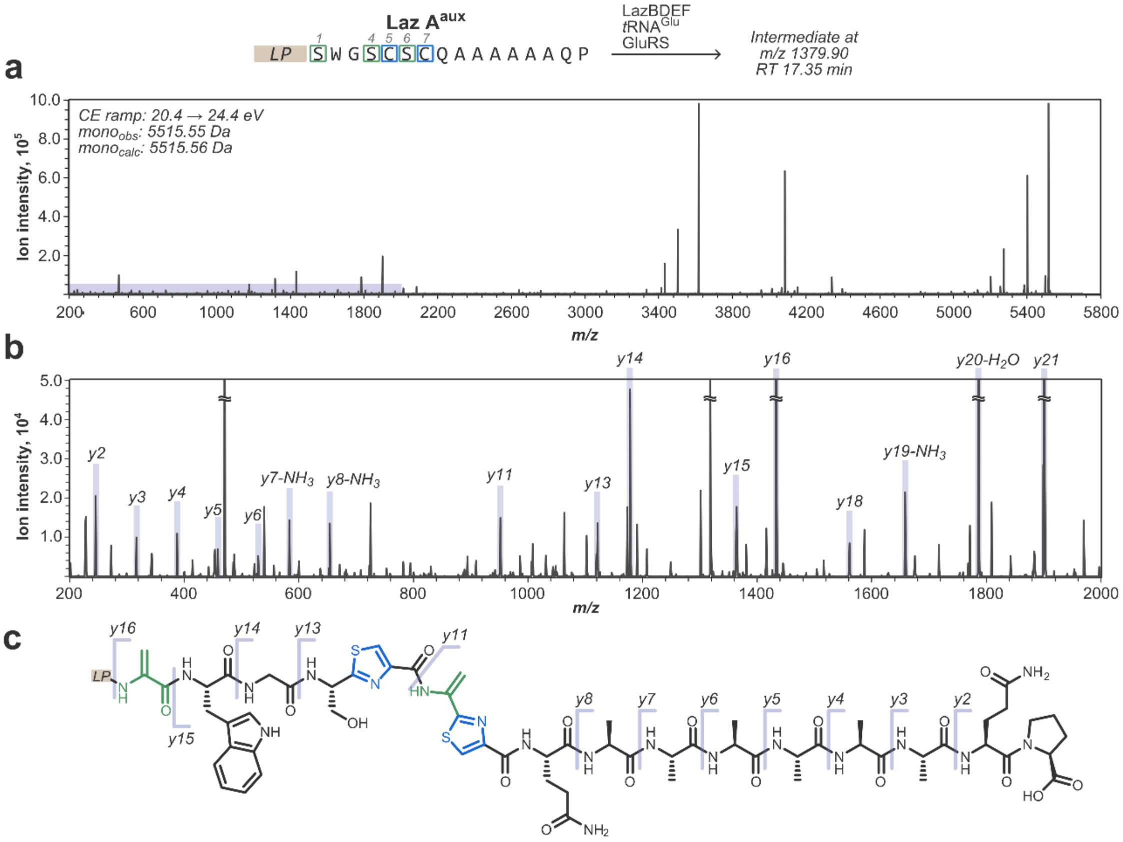

**Figure S31**. Annotated MS/MS spectrum for Dha1/Thz5-Dha6-Thz7 LazA$^{aux}$ intermediate captured after treating LazA$^{aux}$ with LazBDEF/GluRS/tRNA$^{Glu}$. a) Full charge-deconvoluted CID fragmentation spectrum for the $z$=4 precursor ion ($m/z$ 1379.90; mono$_{calc}$: 5515.56 Da; mono$_{obs}$: 5515.56 Da) obtained with collision energies ramped from 20.4 to 24.4 eV. b) A zoomed-in fraction of the spectrum corresponding to the shaded area from panel a) with spectral assignments; *y*-ions are annotated; most *b*-ions, stable molecule losses ($H_2O$, $NH_3$, CO, etc.) and double fragmentation events are omitted for clarity. c) Chemical structure of Dha1/Thz5-Dha6-Thz7 LazA$^{min}$ CP with mapped *y*-ion annotations. Ions *y8-NH$_3$, y11, y13, y15 and y16* collectively support the assignment.

#### 3.5. LazA$^{wt}$ time course assignments

Assignments of PTM patterns on captured intermediates were done following the scheme outlined in section 3.1 and from MS/MS data. See also section 3.3 for assignment of shunt products. Table S3 summarizes observed intermediates and shunt products. Figures S32-S41 show data to support these annotations.

**Table S3**. Progress of LazA$^{wt}$ biosynthesis in the FIT-Laz system. For each intermediate, *m/z* values correspond to the monoisotopic mass peaks. RT stands for HPLC retention time (method B).

| ***m/z*** | **RT, min** | **assignment** | **notes** | **MS/MS assignment** | **reaction time, min** | | | | | |
|---|---|---|---|---|---|---|---|---|---|---|
| | | | | | **0** | **5** | **15** | **30** | **45** | **60** |
| 1026.26 | 15.37 | LP-$NH_2$ | Macrocyclization product | | 0.0 | 1.7 | 40.6 | 56.7 | 70.8 | 73.7 |
| 1381.38 | 18.1 | Dha1, 6, 10, 12<br>Thz5, 7, 13<br>Oxz11 | Macrocyclization substrate | | 0.0 | 0.0 | 1.4 | 0.7 | 0.3 | 0.2 |
| 1386.38 | 17.42 | Dha1, 6, 10, 12<br>Thz5, 7<br>Oxz11<br>Dha1 to Cys13<br>Lanthipeptide | shunt product | | 0.0 | 0.0 | 2.7 | 5.6 | 4.8 | 5.8 |
| 1385.38 | 17.76 | Dha6, 10, 12<br>Thz5, 7, 13<br>Oxz11 | | Fig. S41 | 0.0 | 0.0 | 0.7 | 1.2 | 0.9 | 0.0 |
| 1390.88 | 17.17 | Dha1, 6, 12<br>Thz5, 7<br>Oxn11<br>Thn13 | | | 0.0 | 0.0 | 7.4 | 10.7 | 7.5 | 3.7 |
| 1395.38 | 16.69 | Thz5, 7<br>Dha6, 12<br>Oxn11<br>Thn13 | | Fig. S40 | 0.0 | 0.0 | 5.9 | 2.6 | 0.0 | 0.0 |

| | | | | | | | | | | |
|---|---|---|---|---|---|---|---|---|---|---|
| 1399.9 | 17.25 | Thz5, 7<br>Dha1, 6<br>Dha1 to Cys13<br>Lanthipeptide | isomer 1; shunt product | | 0.0 | 0.0 | 3.2 | 4.0 | 4.1 | 5.0 |
| 1399.9 | 17.42 | | isomer 2; shunt product | | 0.0 | 0.0 | 4.5 | 5.1 | 5.5 | 7.0 |
| 1400.38 | 17.67 | Thz5<br>Thn7, 13<br>Dha12 | | Fig. S39 | 0.0 | 0.0 | 8.8 | 7.7 | 4.1 | 2.2 |
| 1414.14 | 16.92 | Thz5, 7<br>Dha6, 12 | | Fig. S38 | 0.0 | 4.7 | 6.0 | 1.0 | 0.0 | 0.0 |
| 1418.65 | 16.47 | Thz5, 7<br>Dha6 | | Fig. S37 | 0.0 | 41.6 | 13.0 | 1.2 | 0.0 | 0.6 |
| 1423.65 | 15.61 | Thz5<br>Thn7 | | Fig. S36 | 0.0 | 10.5 | 0.9 | 0.0 | 0.0 | 0.0 |
| 1438.41 | 15.9 | Thn5<br>Dha6 | | Fig. S35 | 0.0 | 5.5 | 0.0 | 0.0 | 0.0 | 0.0 |
| 1442.91 | 15.36 | Thn5 | | Fig. S34 | 0.0 | 11.2 | 0.0 | 0.0 | 0.0 | 0.0 |
| 1457.17 | 15.63 | Dha1 | | | 0.0 | 3.3 | 0.0 | 0.0 | 0.0 | 0.0 |
| 1461.68 | 15.26 | S.M. | | Fig. S33 | 100.0 | 14.4 | 1.6 | 0.5 | 0.6 | 0.3 |
| | | minor products, combined | | | 0.0 | 7.1 | 3.1 | 2.9 | 1.4 | 1.5 |
| | | | | | thiopeptide intensity, a. u. | | | | | |
| 701.20 | 12.81 | thiopeptide | | | 0.0 | 0.2 | 10.1 | 11.0 | 12.9 | 12.4 |

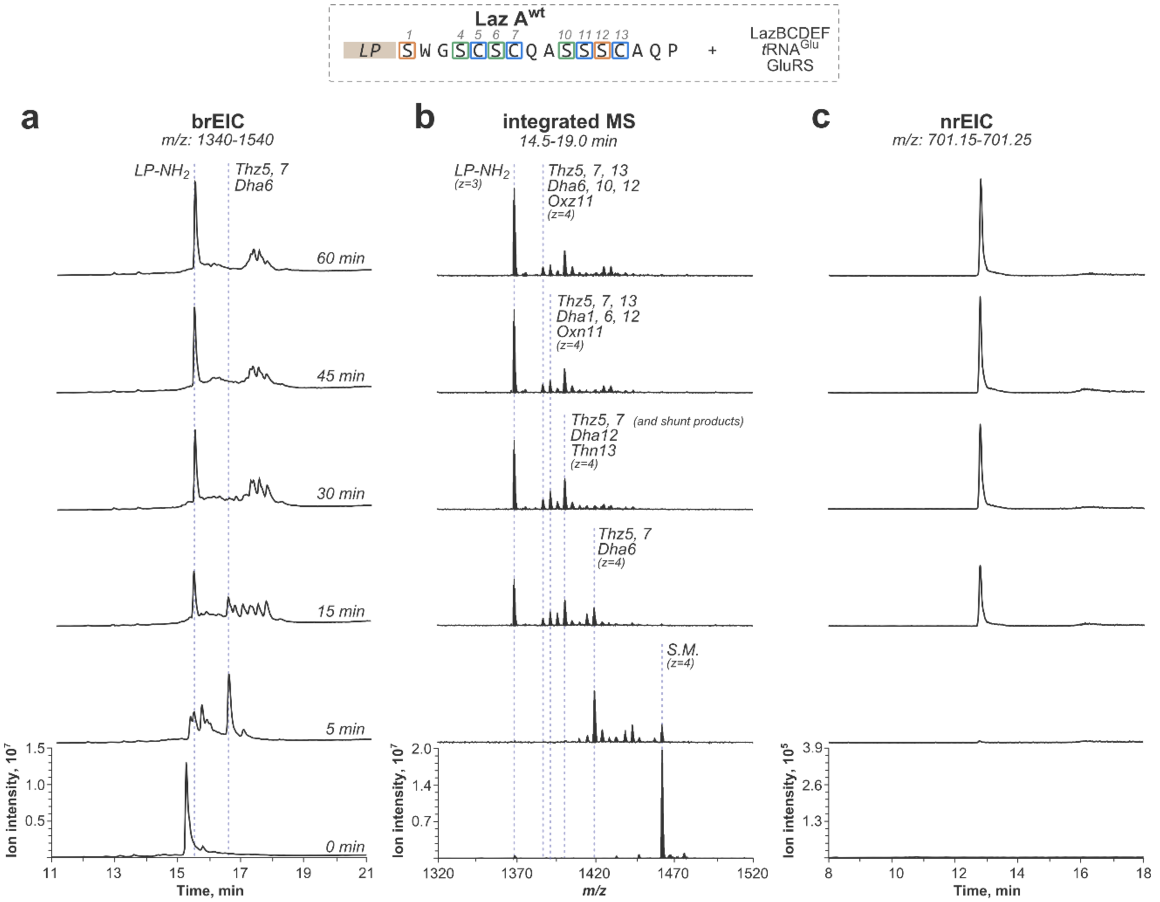

**Figure S32**. Progression of lactazole A biosynthesis in the FIT-Laz system. Translation-derived precursor peptide, LazA$^{wt}$, was incubated with LazBCDEF/GluRS/tRNA$^{Glu}$ for specified time and the outcomes were analyzed by LC-MS. a) Overview $^{br}$EIC chromatograms visualizing accumulation and consumption of linear intermediates as they are modified from LazA$^{wt}$ (0 min) to LP-NH$_2$ (60 min). b) Integrated mass spectra corresponding to chromatograms from panel a) with annotations for select individual intermediates forming during the process. c) $^{nr}$EIC chromatograms for lactazole A accumulating alongside LP-NH$_2$. Y-axes are scaled internally for each panel.

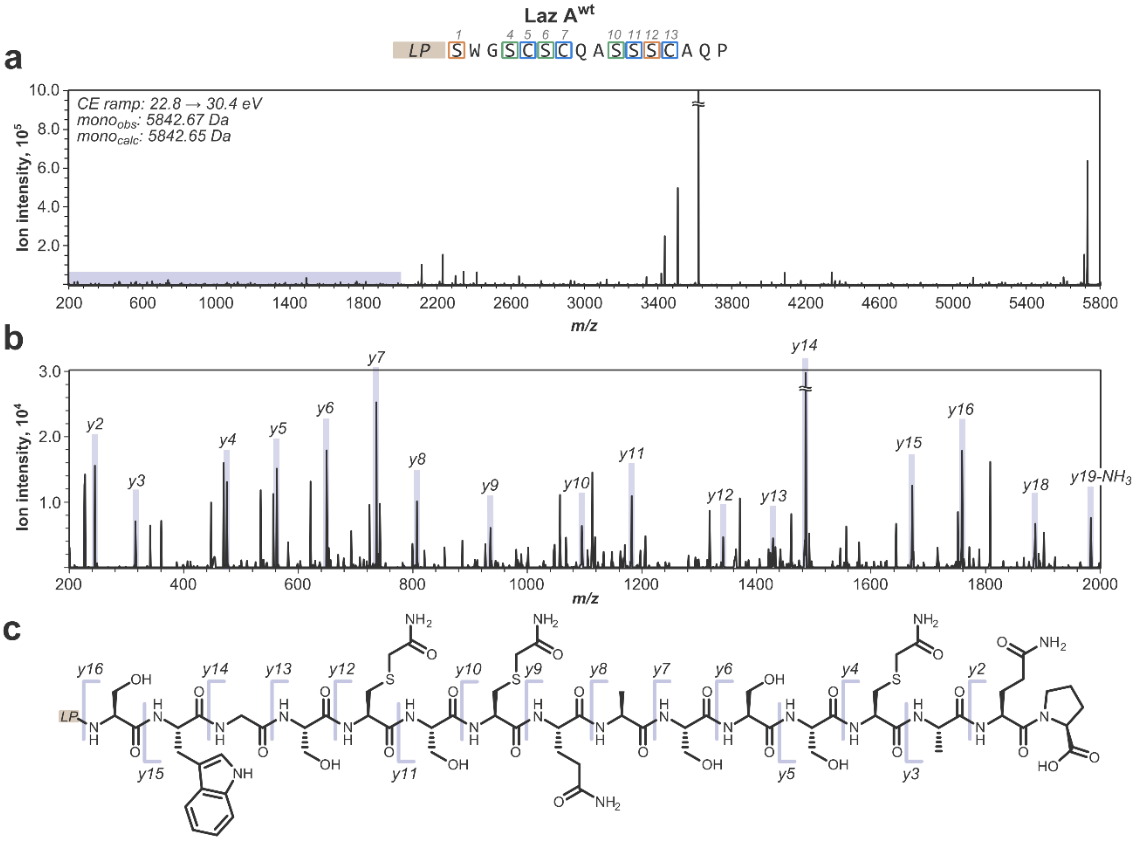

**Figure S33**. Annotated MS/MS spectrum for the unmodified peptide, i.e. LazA[wt]. a) Full charge-deconvoluted CID fragmentation spectrum for the $z$=4 precursor ion ($m/z$ 1461.68; $mono_{calc}$: 5842.65 Da; $mono_{obs}$: 5842.67 Da) obtained with collision energies ramped from 22.8 to 30.4 eV. b) A zoomed-in fraction of the spectrum corresponding to the shaded area from panel a) with spectral assignments; $y$-ions are annotated; most $b$-ions, stable molecule losses ($H_2O$, $NH_3$, CO, etc.) and double fragmentation events are omitted for clarity. c) Chemical structure of LazA[wt] CP with mapped $y$-ion annotations; full $y$-ion ladder is observed.

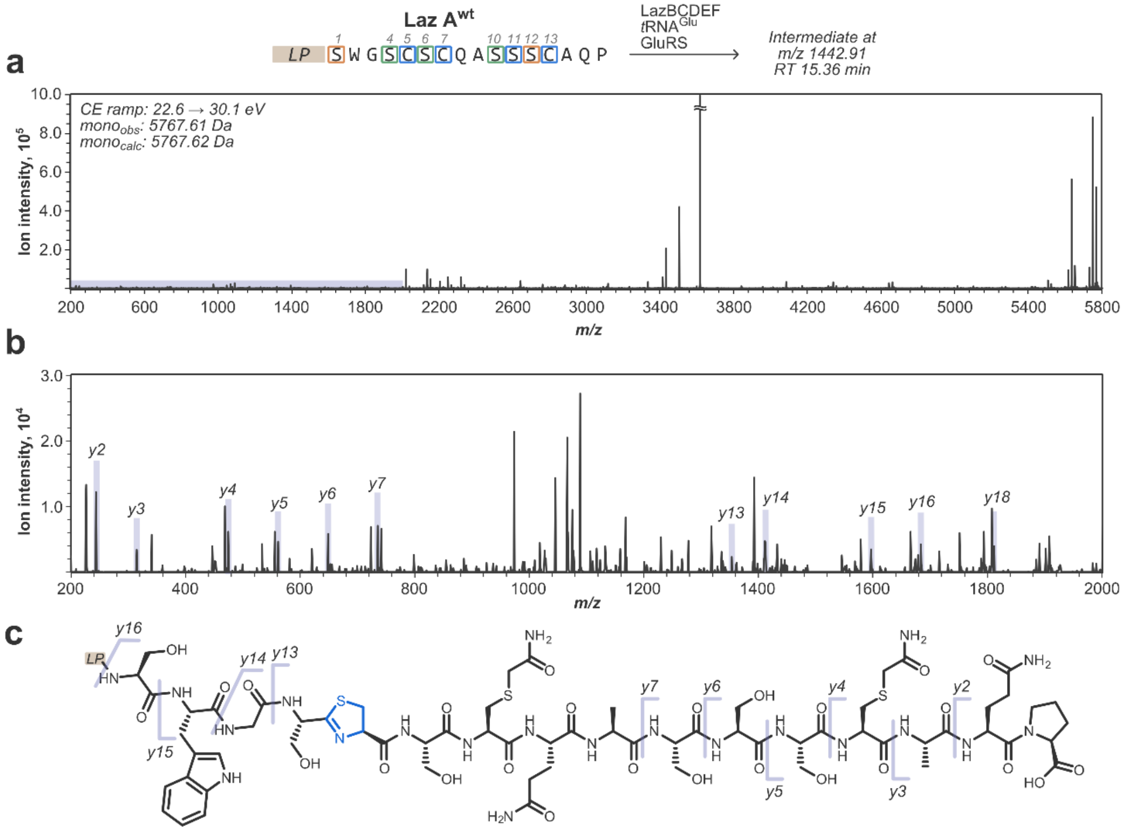

**Figure S34**. Annotated MS/MS spectrum for Thn5 LazA$^{wt}$ intermediate captured after treating LazA$^{wt}$ with LazBCDEF/GluRS/tRNA$^{Glu}$. a) Full charge-deconvoluted CID fragmentation spectrum for the $z$=4 precursor ion (*m/z* 1442.91; mono$_{calc}$: 5767.62 Da; mono$_{obs}$: 5767.61 Da) obtained with collision energies ramped from 22.6 to 30.1 eV. b) A zoomed-in fraction of the spectrum corresponding to the shaded area from panel a) with spectral assignments; *y*-ions are annotated; *most b-ions, stable molecule* losses ($H_2O$, $NH_3$, CO, etc.) and double fragmentation events are omitted for clarity. c) Chemical structure of Thn5 LazA$^{wt}$ CP with mapped *y*-ion annotations. The *y3/y4* ion pair indicates that Cys13 is unmodified. No fragmentation between Cys5 and Cys7 is observed, making unambiguous localization of Thn position impossible; assignment was made by analogy to LazA$^{aux}$.

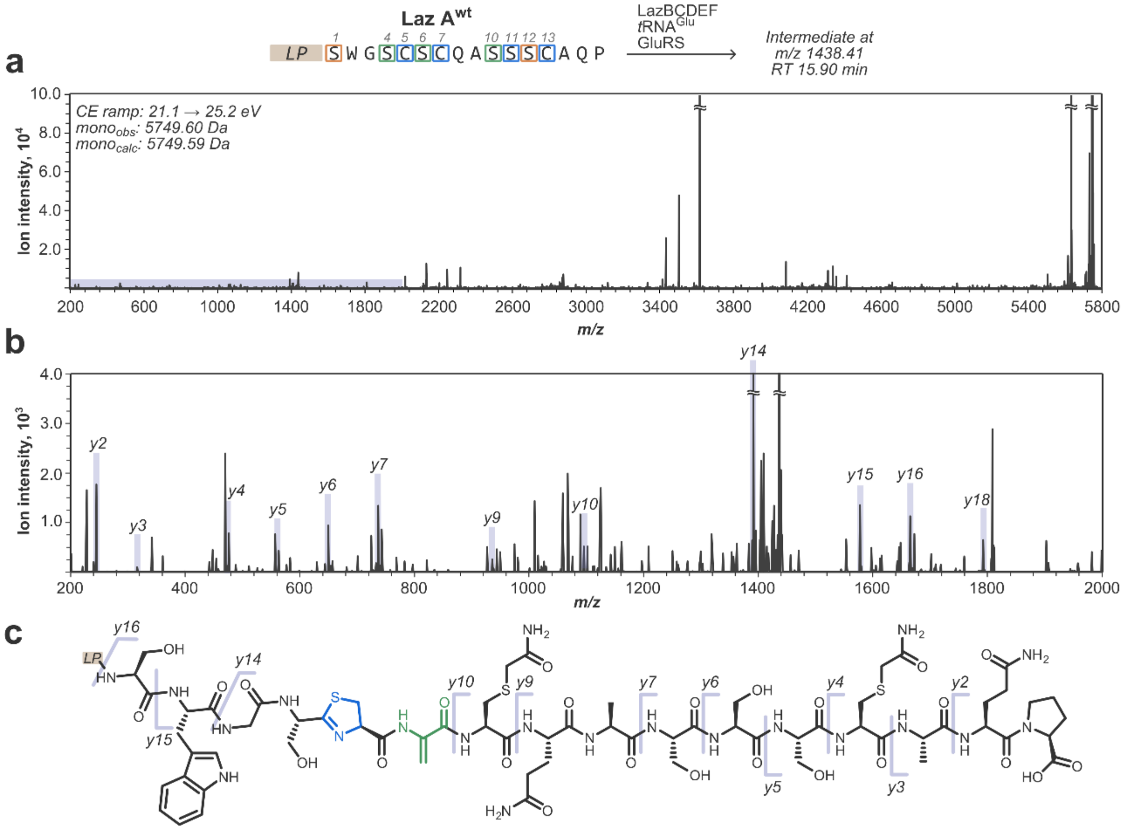

**Figure S35**. Annotated MS/MS spectrum for Thn5-Dha6 LazA^wt intermediate captured after treating LazA^wt with LazBCDEF/GluRS/tRNA^Glu. a) Full charge-deconvoluted CID fragmentation spectrum for the $z$=4 precursor ion (*m/z* 1438.41; $mono_{calc}$: 5749.59 Da; $mono_{obs}$: 5749.60 Da) obtained with collision energies ramped from 21.1 to 25.2 eV. b) A zoomed-in fraction of the spectrum corresponding to the shaded area from panel a) with spectral assignments; *y*-ions are annotated; most *b*-ions, stable molecule losses ($H_2O$, $NH_3$, CO, etc.) and double fragmentation events are omitted for clarity. c) Chemical structure of Thn5-Dha6 LazA^wt CP with mapped *y*-ion annotations. The *y3/y4* and *y9/y10* ion pairs suggest that Thn is localized at Cys5. Fragments *y10* and *y15/y16* support the assignment of Dha at Ser6.

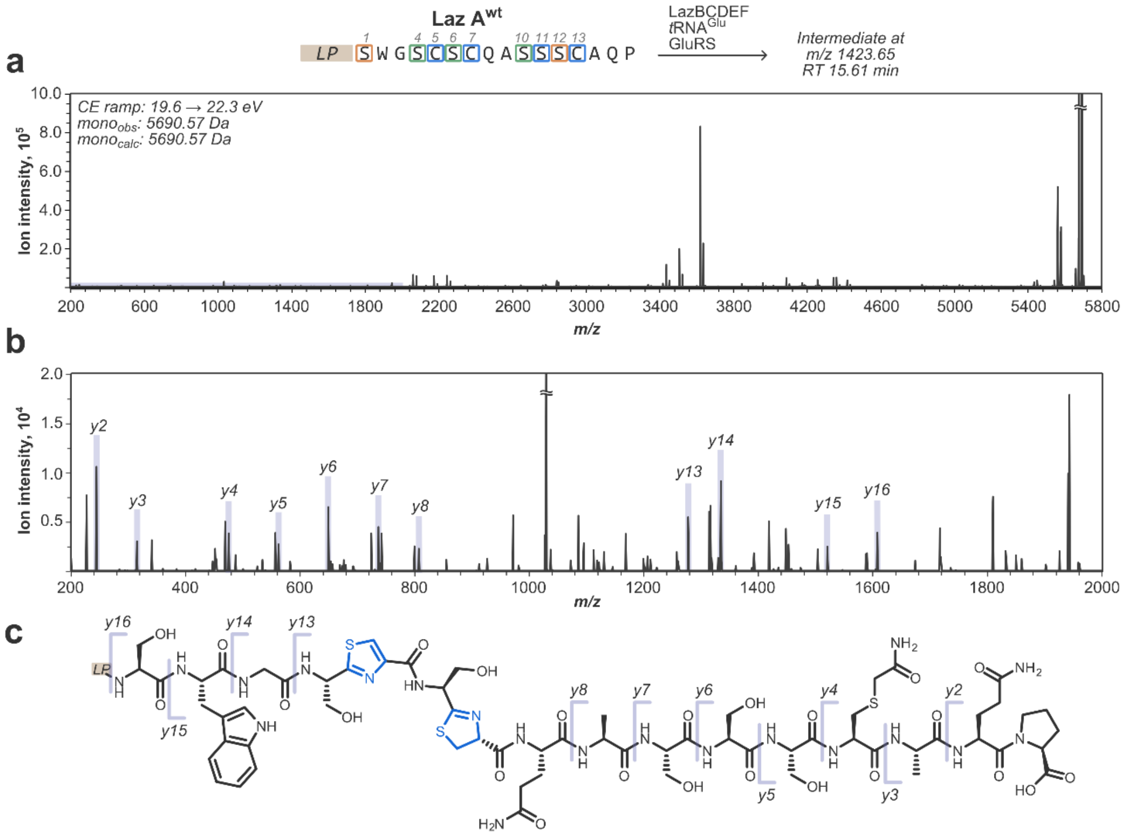

**Figure S36**. Annotated MS/MS spectrum for Thz5/Thn7 LazA$^{wt}$ intermediate captured after treating LazA$^{wt}$ with LazBCDEF/GluRS/tRNA$^{Glu}$. a) Full charge-deconvoluted CID fragmentation spectrum for the *z*=4 precursor ion (*m/z* 1423.65; mono$_{calc}$: 5690.57 Da; mono$_{obs}$: 5690.57 Da) obtained with collision energies ramped from 19.6 to 22.3 eV. b) A zoomed-in fraction of the spectrum corresponding to the shaded area from panel a) with spectral assignments; *y*-ions are annotated; most *b*-ions, stable molecule losses ($H_2O$, $NH_3$, CO, etc.) and double fragmentation events are omitted for clarity. c) Chemical structure of Thz5/Thn7 LazA$^{wt}$ CP with mapped *y*-ion annotations. The *y3/y4* ion pair indicates that Cys13 is unmodified. No fragmentation between Cys5 and Cys7 is observed, making unambiguous localization of Thz vs Thn position impossible; assignment was made by analogy to LazA$^{aux}$.

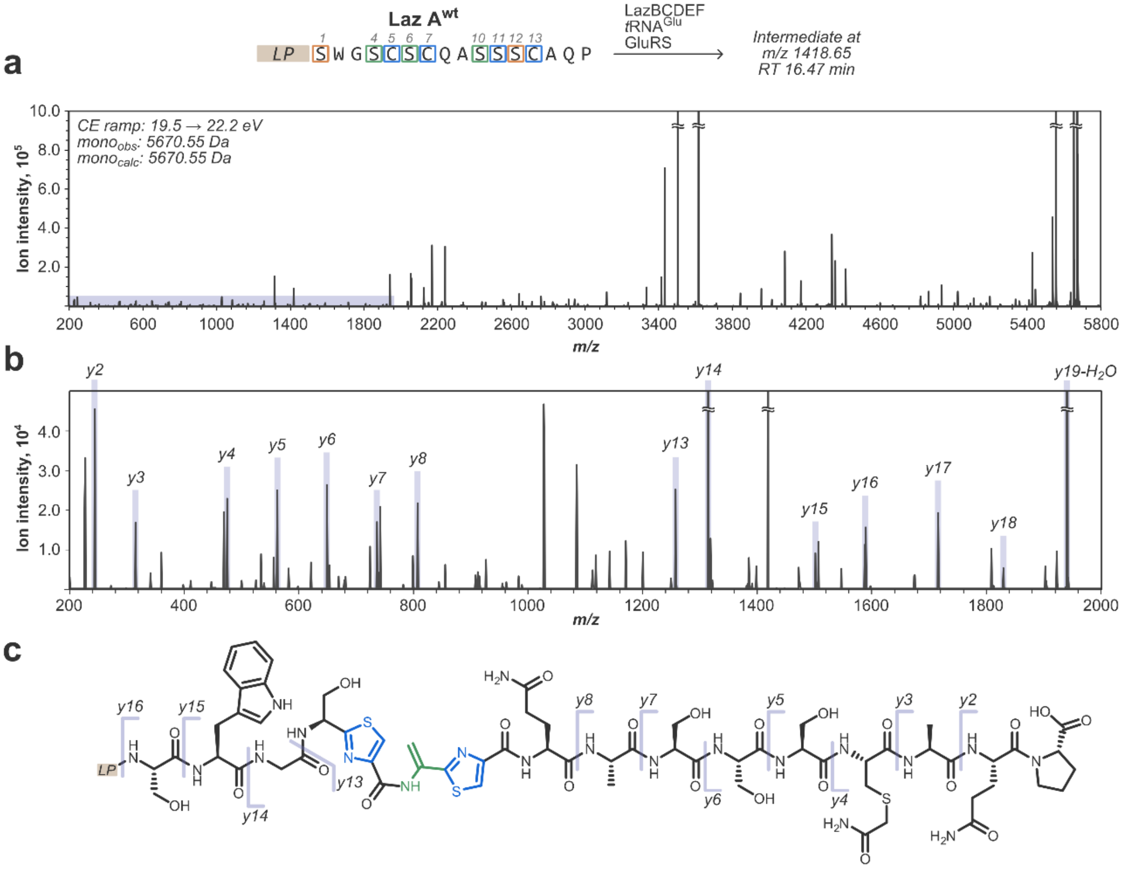

**Figure S37**. Annotated MS/MS spectrum for Ser4-Thz5-Dha6-Thz7 LazA$^{wt}$ intermediate captured after treating LazA$^{wt}$ with LazBCDEF/GluRS/tRNA$^{Glu}$. a) Full charge-deconvoluted CID fragmentation spectrum for the *z*=4 precursor ion (*m/z* 1418.65; mono$_{calc}$: 5670.55 Da; mono$_{obs}$: 5670.55 Da) obtained with collision energies ramped from 19.5 to 22.2 eV. b) A zoomed-in fraction of the spectrum corresponding to the shaded area from panel a) with spectral assignments; *y*-ions are annotated; most *b*-ions, stable molecule losses ($H_2O$, $NH_3$, CO, etc.) and double fragmentation events are omitted for clarity. c) Chemical structure of Thz5-Dha6-Thz7 LazA$^{wt}$ CP with mapped *y*-ion annotations. The *y8/y13* ion pair supports the assignment.

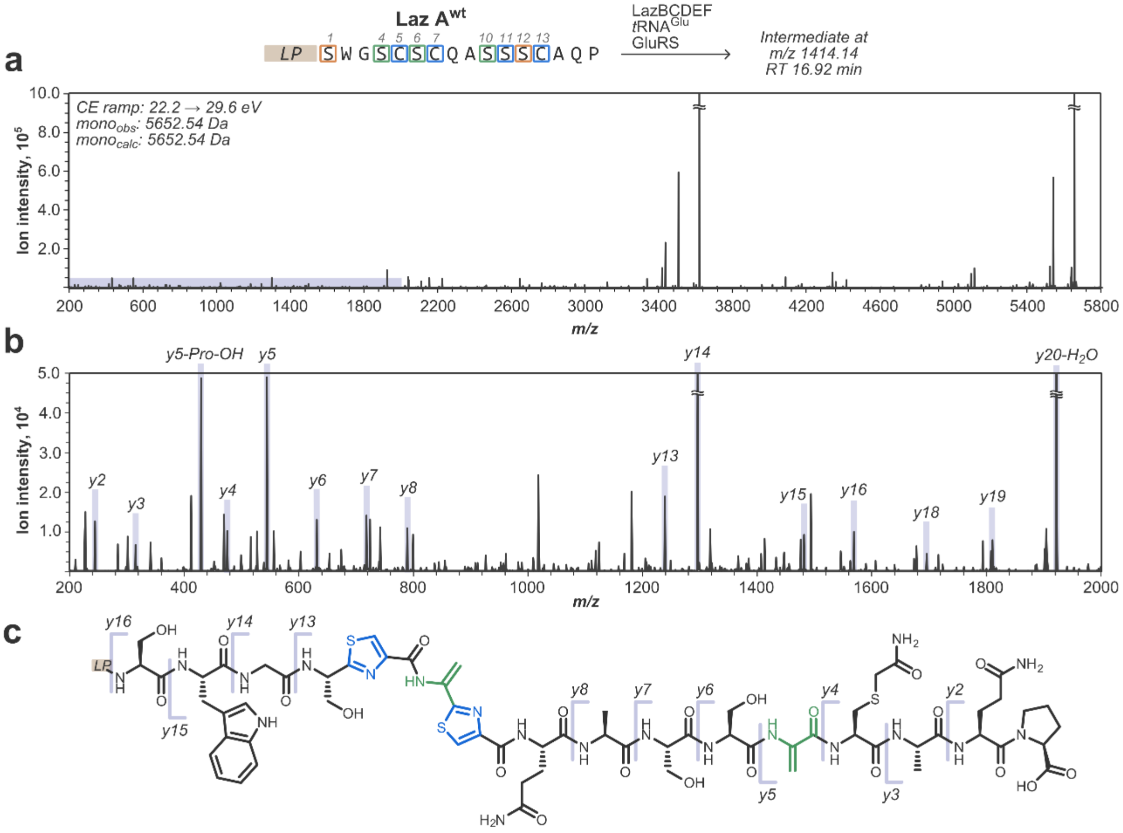

**Figure S38**. Annotated MS/MS spectrum for Thz5-Dha6-Thz7/Dha12 LazA$^{wt}$ intermediate captured after treating LazA$^{wt}$ with LazBCDEF/GluRS/tRNA$^{Glu}$. a) Full charge-deconvoluted CID fragmentation spectrum for the *z*=4 precursor ion (*m/z* 1414.14; mono$_{calc}$: 5652.54 Da; mono$_{obs}$: 5652.54 Da) obtained with collision energies ramped from 22.2 to 29.6 eV. b) A zoomed-in fraction of the spectrum corresponding to the shaded area from panel a) with spectral assignments; *y*-ions are annotated; most *b*-ions, stable molecule losses ($H_2O$, $NH_3$, CO, etc.) and double fragmentation events are omitted for clarity. c) Chemical structure of Thz5-Dha6-Thz7/Dha12 LazA$^{wt}$ CP with mapped *y*-ion annotations. An intense *y5* ion and its derivatives support the Dha12 assignment. PTMs at residues 5-7 have a fragmentation pattern similar to Fig. S36.

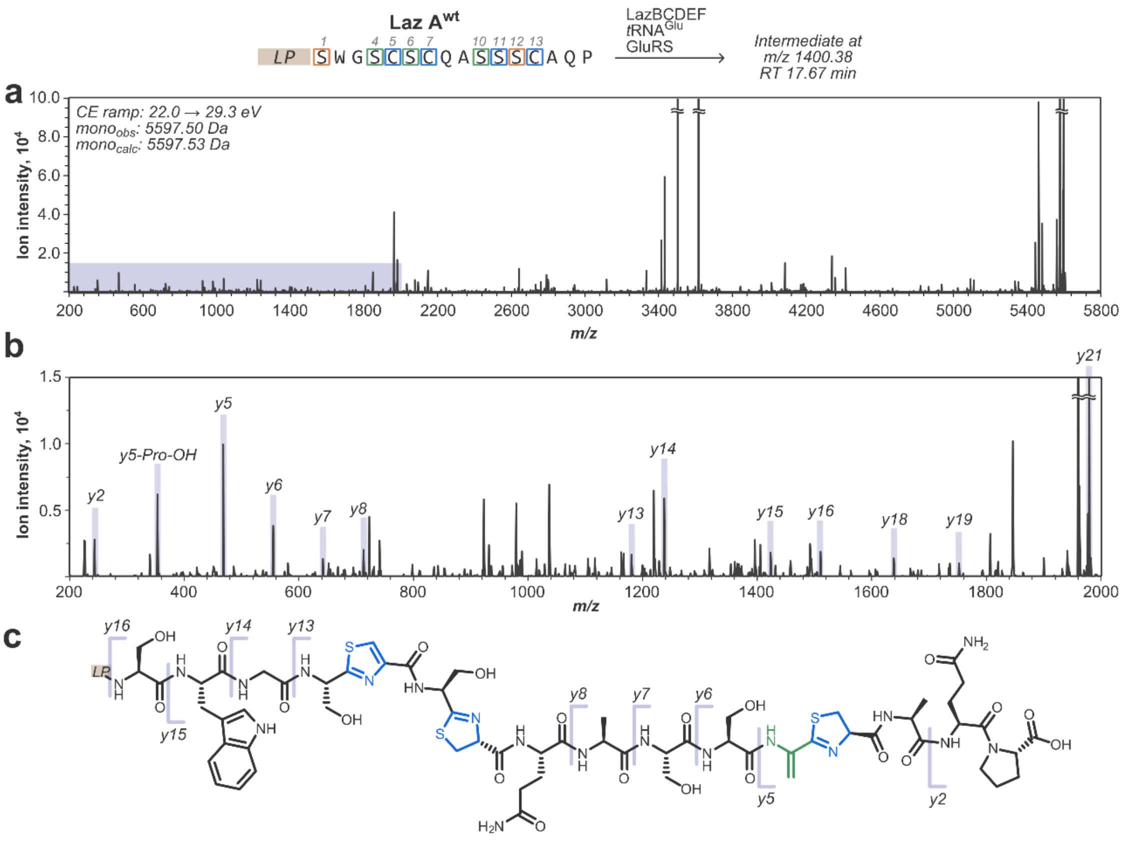

**Figure S39**. Annotated MS/MS spectrum for Thz5/Thn7/Dha12-Thn13 LazA[wt] intermediate captured after treating LazA[wt] with LazBCDEF/GluRS/tRNA[Glu]. a) Full charge-deconvoluted CID fragmentation spectrum for the *z*=4 precursor ion (*m/z* 1400.38; $mono_{calc}$: 5597.53 Da; $mono_{obs}$: 5597.50 Da) obtained with collision energies ramped from 22.0 to 29.3 eV. b) A zoomed-in fraction of the spectrum corresponding to the shaded area from panel a) with spectral assignments; *y*-ions are annotated; most *b*-ions, stable molecule losses ($H_2O$, $NH_3$, CO, etc.) and double fragmentation events are omitted for clarity. c) Chemical structure of Thz5/Thn7/Dha12-Thn13 LazA[wt] CP with mapped *y*-ion annotations. An intense *y5* ion and its derivatives support the Dha12-Thn13 assignment; PTM pattern around residues 5-7 was assigned by analogy to Fig. S36. This intermediate is the only observed case of modification at residues 10-13 taking place prior to the installation of PTMs at amino acids 4-7.

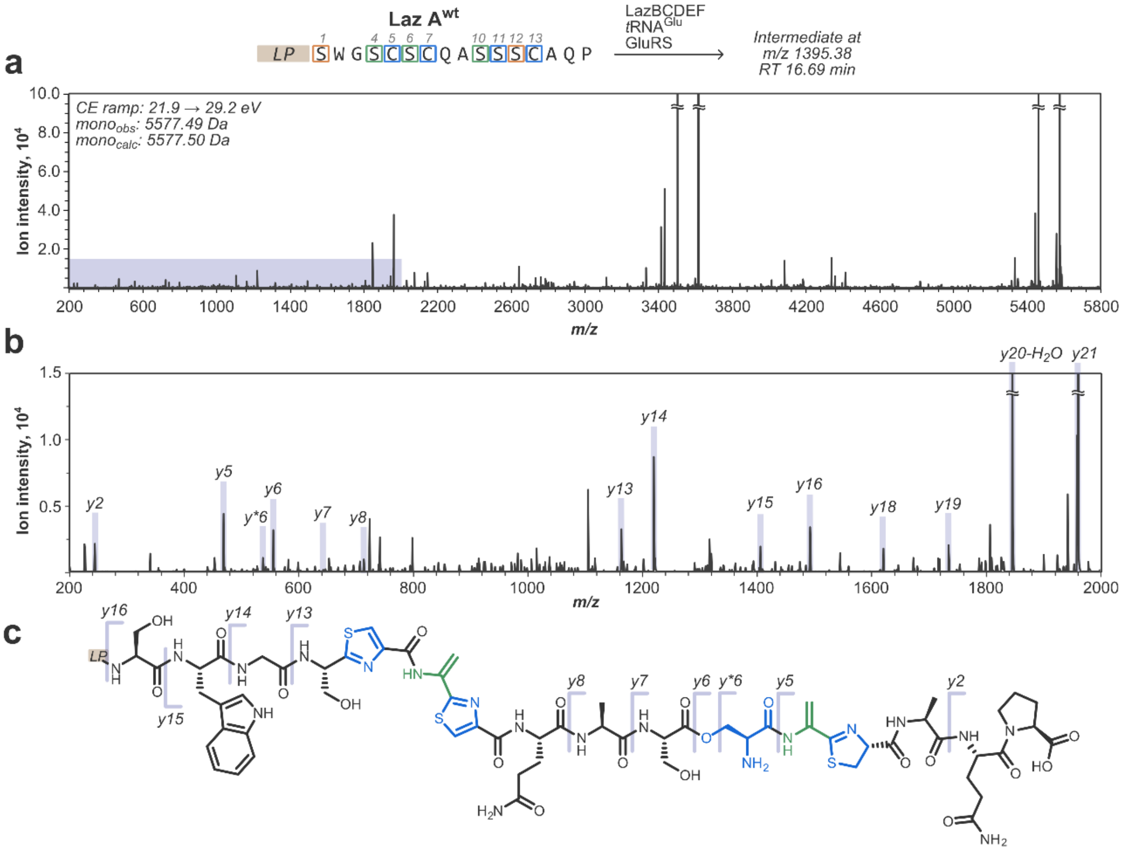

**Figure S40**. Annotated MS/MS spectrum for Thz5-Dha6-Thz7/Oxn11-Dha12-Thn13 LazA$^{wt}$ intermediate captured after treating LazA$^{wt}$ with LazBCDEF/GluRS/tRNA$^{Glu}$. a) Full charge-deconvoluted CID fragmentation spectrum for the $z$=4 precursor ion ($m/z$ 1395.38; mono$_{calc}$: 5577.50 Da; mono$_{obs}$: 5577.49 Da) obtained with collision energies ramped from 21.9 to 29.2 eV. b) A zoomed-in fraction of the spectrum corresponding to the shaded area from panel a) with spectral assignments; *y*-ions are annotated; most *b*-ions, stable molecule losses ($H_2O$, $NH_3$, CO, etc.) and double fragmentation events are omitted for clarity. c) Chemical structure of Thz5-Dha6-Thz7/Oxn11-Dha12-Thn13 LazA$^{wt}$ CP with mapped *y*-ion annotations. The *y5* ion supports the Dha12-Thn13 assignment; he *y8/y13* ion pair confirms PTMs at residues 5-7. For notes regarding formation of the *y*$^{*}$*6* ion, refer to Fig. S10.

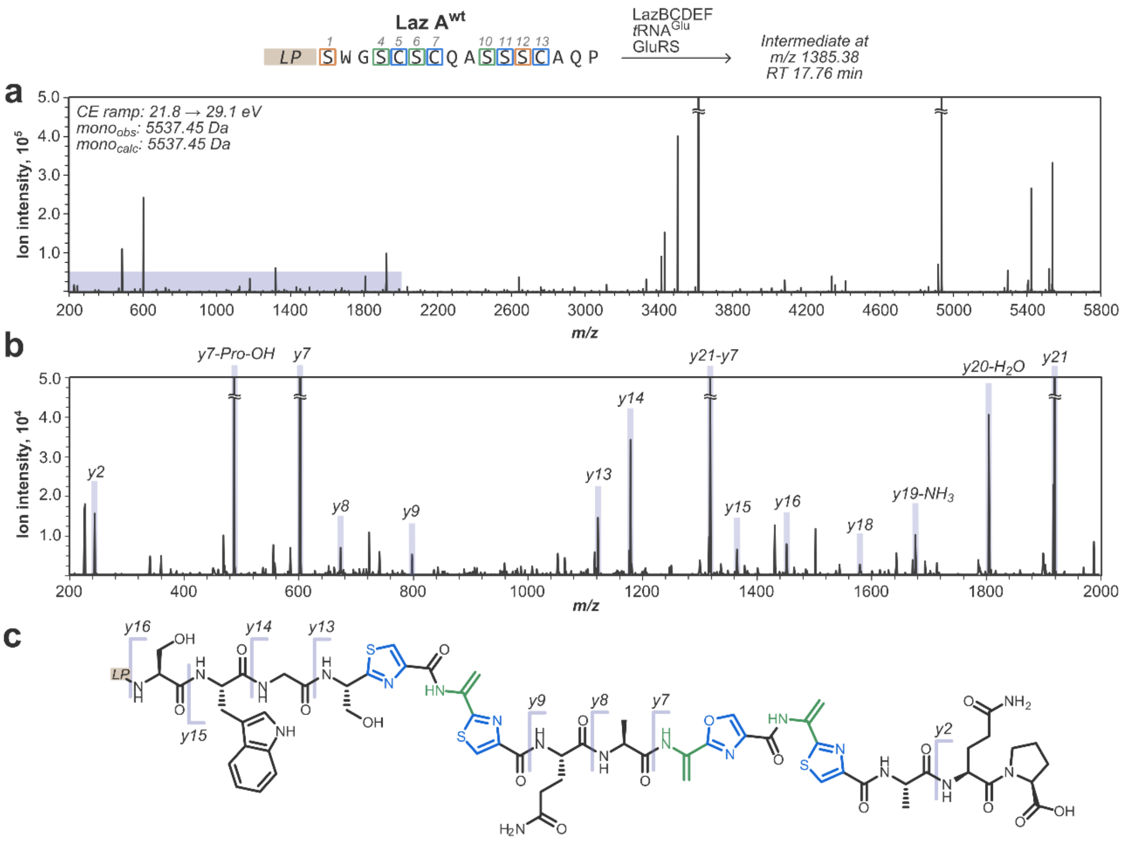

**Figure S41**. Annotated MS/MS spectrum for Thz5-Dha6-Thz7/Dha10-Oxz11-Dha12-Thz13 LazA$^{wt}$ intermediate captured after treating LazA$^{wt}$ with LazBCDEF/GluRS/tRNA$^{Glu}$. a) Full charge-deconvoluted CID fragmentation spectrum for the $z$=4 precursor ion ($m/z$ 1385.38; mono$_{calc}$: 5537.45 Da; mono$_{obs}$: 5537.45 Da) obtained with collision energies ramped from 21.8 to 29.1 eV. b) A zoomed-in fraction of the spectrum corresponding to the shaded area from panel a) with spectral assignments; $y$-ions are annotated; *most b-ions, stable molecule* losses ($H_2O$, $NH_3$, CO, etc.) and double fragmentation events are omitted for clarity. c) Chemical structure of Thz5-Dha6-Thz7/Dha10-Oxz11-Dha12-Thz13 LazA$^{wt}$ CP with mapped $y$-ion annotations. The *y7* ion supports the proposed pattern of modifications in residues 10-13; the *y9/y13* ion pair confirms PTMs at residues 5-7, and the *y15/16* ions confirm that Ser1 remains unmodified.

#### 3.6. Substrate specificity study

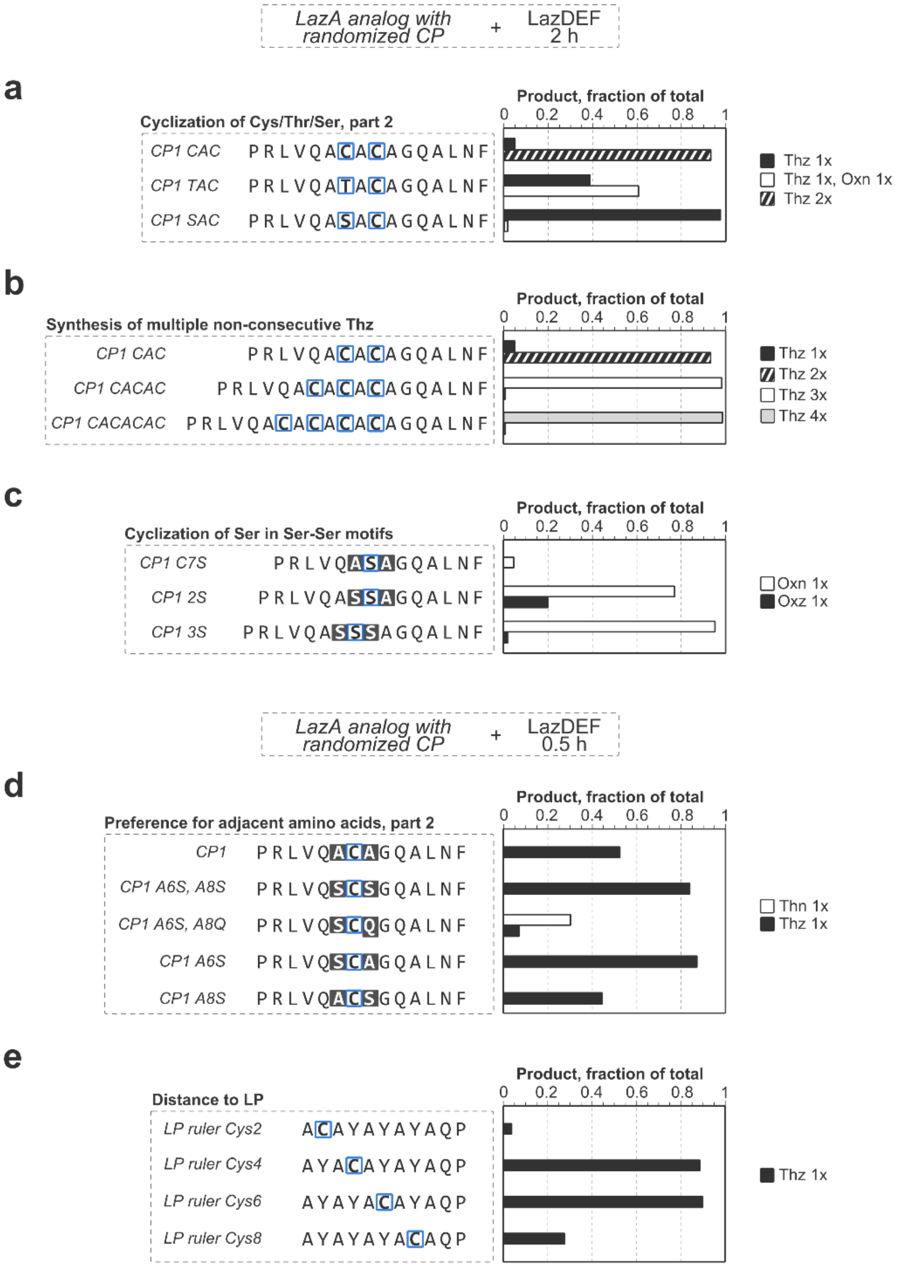

**Figure S42**. Supplementary data for LazDE specificity study. See also Fig. 5. LazA$^{CP1}$ variants were treated by LazDEF for 2 h or 0.5 h, and the outcomes were analyzed by LC-MS. a) The extent of Cys/Thr/Ser heterocyclization does not change when a Thz PTM is present in position +2 (compare with Fig. 5a). b) In contrast to heterocyclization of consecutive Cys (Fig. 5a), formation of multiple Thz in sequences with non-consecutive Cys is efficient. c) LazDE displays strong preference for a Ser residue in position –1. Whereas Ala-Ser-containing CPs remain essentially unmodified after a 2 h incubation, Ser-Ser motifs are excellent substrates. These results explain the preference of LazDE for Ser11 in $LazA^{wt}$ and $LazA^{min}$ over other Ser residues. d) $LazA^{CP1}$ variants with grafted "native" tripeptide sequences were treated by LazDEF for 0.5 h to better differentiate the influence of local sequence environments. The results were analyzed by LC-MS and quantified as described in section 2.5. These data suggest that i) Ser in position –1 is preferred over Ala, and ii) LazF-mediated dehydrogenation of Thn inside the Ala-Thn-Gln motif is slow. e) The effect of the distance between the cyclizable amino acid and the LP was investigated using 4 LazA variants. The peptides were translated in the FIT system and treated with LazDEF for 0.5 h. The results suggest that the enzyme acts in the N- to C-terminal direction and requires a short spacer sequence between the LP and the modifiable residue for efficient heterocyclization.

**a**

| | Sequence | LazBF GluRS tRNA$^{Glu}$ | LazDEF | LazBDEF GluRS tRNA$^{Glu}$ | Azoline/azole-dependent Dha formation |
|---|---|---|---|---|---|
| Laz A$^{min}$ S10A | LP SWGAAAAQAASSCAQP | *Dha1* | *Oxn11* *Thz13* | *Dha1, 12* *Oxn11* *Thz13* | *Ser12*: + |
| Laz A$^{min}$ S11A | LP SWGAAAAQASASCAQP | *Dha1* | *Thz13* | *Dha1* *Thz13* | *Ser10*: – *Ser12*: – |
| Laz A$^{min}$ S12A | LP SWGAAAAQASSACAQP | *Dha1* | *mixture of products*† | *mixture of products*† | *Ser10*: – |
| Laz A$^{min}$ C13A | LP SWGAAAAQASSSAAQP | *Dha1* | *Oxn11* | *Dha1* *Oxn11* | *Ser10*: – *Ser12*: – |

**b**

| | Sequence | LazBF GluRS tRNA$^{Glu}$ | LazDEF | LazBDEF GluRS tRNA$^{Glu}$ | |
|---|---|---|---|---|---|
| Laz A$^{aux}$ S4A | LP SWGACSCQAAAAAAQP | *Dha1* | *Thz5, 7* | *Dha1, 6* *Thz5, 7* | *Ser6*: + |
| Laz A$^{aux}$ C5A | LP SWGSASCQAAAAAAQP | *Dha1* | *Thz7* | *Dha1* *Thz7* | *Ser4*: – *Ser6*: – |
| Laz A$^{aux}$ S6A | LP SWGSCACQAAAAAAQP | *Dha1* | *Thz5, 7* | *Dha1* *Thz5, 7* | *Ser4*: – |
| Laz A$^{aux}$ C7A | LP SWGSCSAQAAAAAAQP | *Dha1* | *Thz5* | *Dha1, 6* *Thz5* | *Ser4*: – *Ser6*: + |

**c**

| | Sequence | LazBDEF GluRS tRNA$^{Glu}$ | |
|---|---|---|---|
| Laz A$^{CP2}$ SC | LP MVNLASCAGGAPQ | *Thz7* | *Ser6*: – |
| Laz A$^{CP2}$ CS | LP MVNLACSAGGAPQ | *mixture of products*§ | *Ser7*: + |
| Laz A$^{CP2}$ SSC | LP MVNLASSCAGGAPQ | *Oxn6* *Dha7* *Thz8* | *Ser7*: + |
| Laz A$^{CP2}$ SSSC | LP MVNLASSSCAGGAPQ | *Dha6, 8* *Oxz7* *Thz9* | *Ser6*: + *Ser8*: + |
| Laz A$^{CP2}$ SASC | LP MVNLASASCAGGAPQ | *Thz9* | *Ser6*: – *Ser8*: – |
| Laz A$^{CP2}$ SCSC | LP MVNLASCSCAGGAPQ | *Dha6, 8* *Thz7, 9* | *Ser6*: + *Ser8*: + |

**Figure S43**. Analysis of azoline/azole-dependent dehydration by LazBF in single point Ala mutants of LazA$^{min}$ (panel a), LazA$^{aux}$ (panel b) or LazA variants with randomized CP (LazA$^{CP2}$, panel c). The peptides were treated with LazBF/GluRS/tRNA$^{Glu}$, LazDEF or LazBDEF/GluRS/tRNA$^{Glu}$ for 2 h and the outcomes were analyzed by LC-MS. Displayed are major observed products for each reaction. Collectively, these data indicate that LazBF prefers Ser in Thz-Ser motifs (LazA$^{aux}$ C7A), whereas dehydration in Ser-Thz (LazA$^{aux}$ C5A) or Oxn-Ser (LazA$^{min}$ C13A) is slow. See also Fig. S44. †: For product distribution, see also Fig. 3b, c. §: For product distribution, see also Fig. S44b.

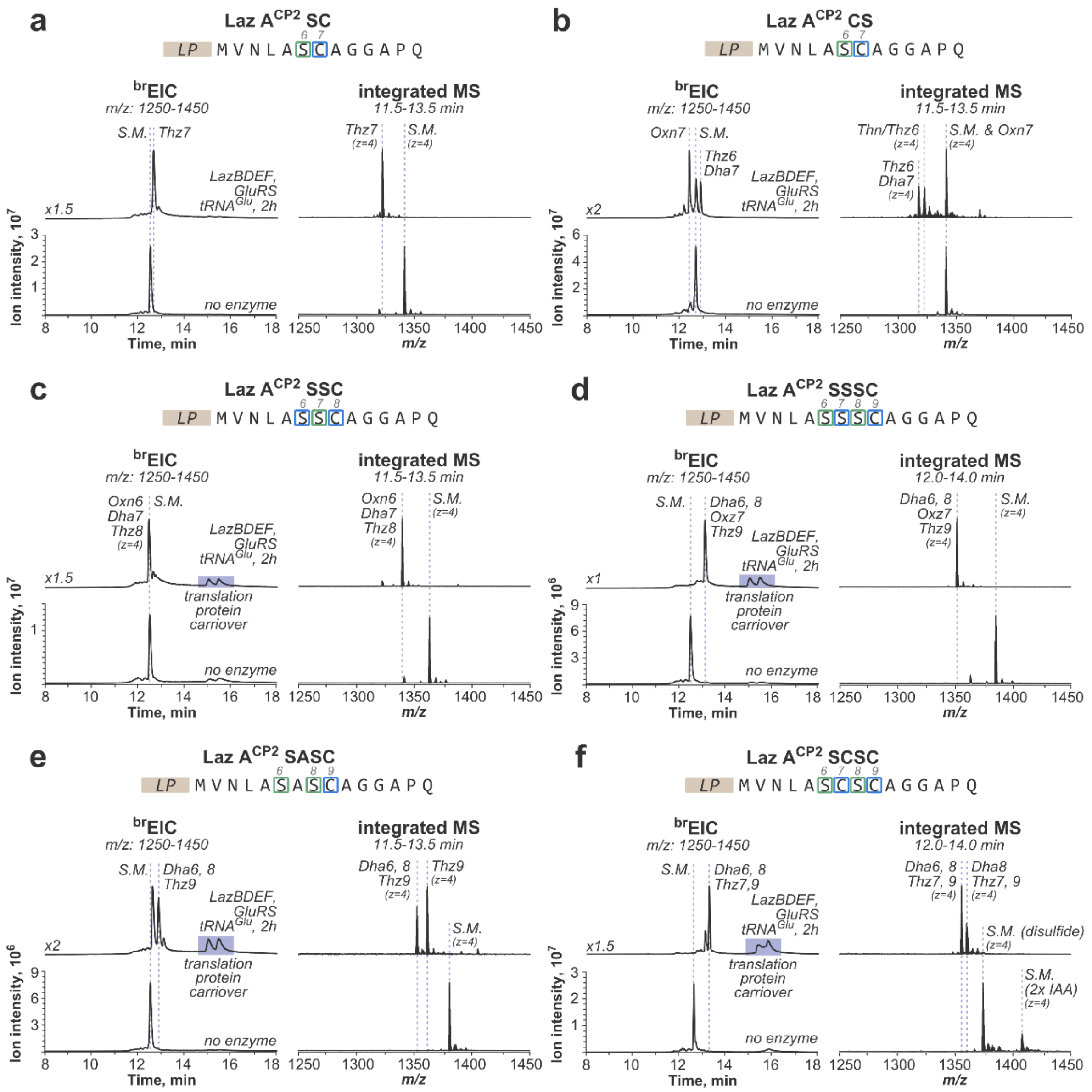

a
Laz A$^{CP2}$ SC
LP MVNLASCAGGAPQ
brEIC
m/z: 1250-1450
integrated MS
11.5-13.5 min
S.M. Thz7
LazBDEF, GluRS tRNA$^{Glu}$, 2h
no enzyme
b
Laz A$^{CP2}$ CS
Oxn7 S.M.
Thz6 Dha7
Thn/Thz6
S.M. & Oxn7
c
Laz A$^{CP2}$ SSC
Oxn6 Dha7 Thz8
translation protein carriover
d
Laz A$^{CP2}$ SSSC
12.0-14.0 min
Dha6, 8 Oxz7 Thz9
e
Laz A$^{CP2}$ SASC
Dha6, 8 Thz9
f
Laz A$^{CP2}$ SCSC
Dha6, 8 Thz7,9
Dha8 Thz7, 9
S.M. (disulfide)
S.M. (2x IAA)
Ion intensity, $10^7$
Ion intensity, $10^6$
Time, min
m/z

**Figure S44**. Analysis of azoline/azole dependent Ser dehydration by LazBF in LazA variants with randomized CP ($LazA^{CP2}$). In each case (panels a-f), the peptides were treated with LazBDEF/GluRS/$tRNA^{Glu}$ for 2 h and reaction outcomes were analyzed by LC-MS. Displayed are $^{br}$EIC chromatograms and integrated mass spectra showing the overall product distribution. These data are consistent with the results from time course study and the single point LazA mutant study (Fig. S44). LazBF prefers to dehydrate Ser in Thz-Ser rather than Ser-Thz motifs (panels a vs. b). Oxn-Ser-Thz (panel c) is also a competent substrate, which is consistent with the data for $LazA^{min}$ S10A (Fig. 3). The Ser-Ser-Ser-Cys motif (panel d) is cleanly modified to the lactazole-like Dha-Oxz-Dha-Thz pattern even when placed inside a randomized CP, highlighting the "local" action of Laz enzymes, and suggesting that diverse thiopeptides should be accessible with FIT-Laz. Modification of the Ser-Cys-Ser-Cys motif (panel f) proceeds relatively fast compared to $LazA^{aux}$ and $LazA^{min}$ S11C (Fig. S21), but still slower than Ser-Ser-Ser-Cys, confirming again, that Ser-Thz-Dha-Thz is a poor substrate for LazBF. Together, these data indicate that despite its local action, LazBF has a sophisticated recognition mode around azoline/azole PTMs.

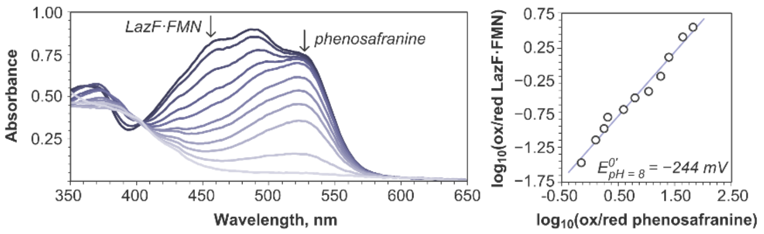

**Figure S45**. Determination of the reduction potential of LazF/FMN following the Massey method (S.I. 2.7). Anaerobic titration of LazF/phenosafranine mixture with a reducing agent ($Na_2S_2O_4$) allows referencing $E^{0'}$(LazF) with respect to the dye ($E^{0'}_{pH = 8}$ = –281.5 mV).

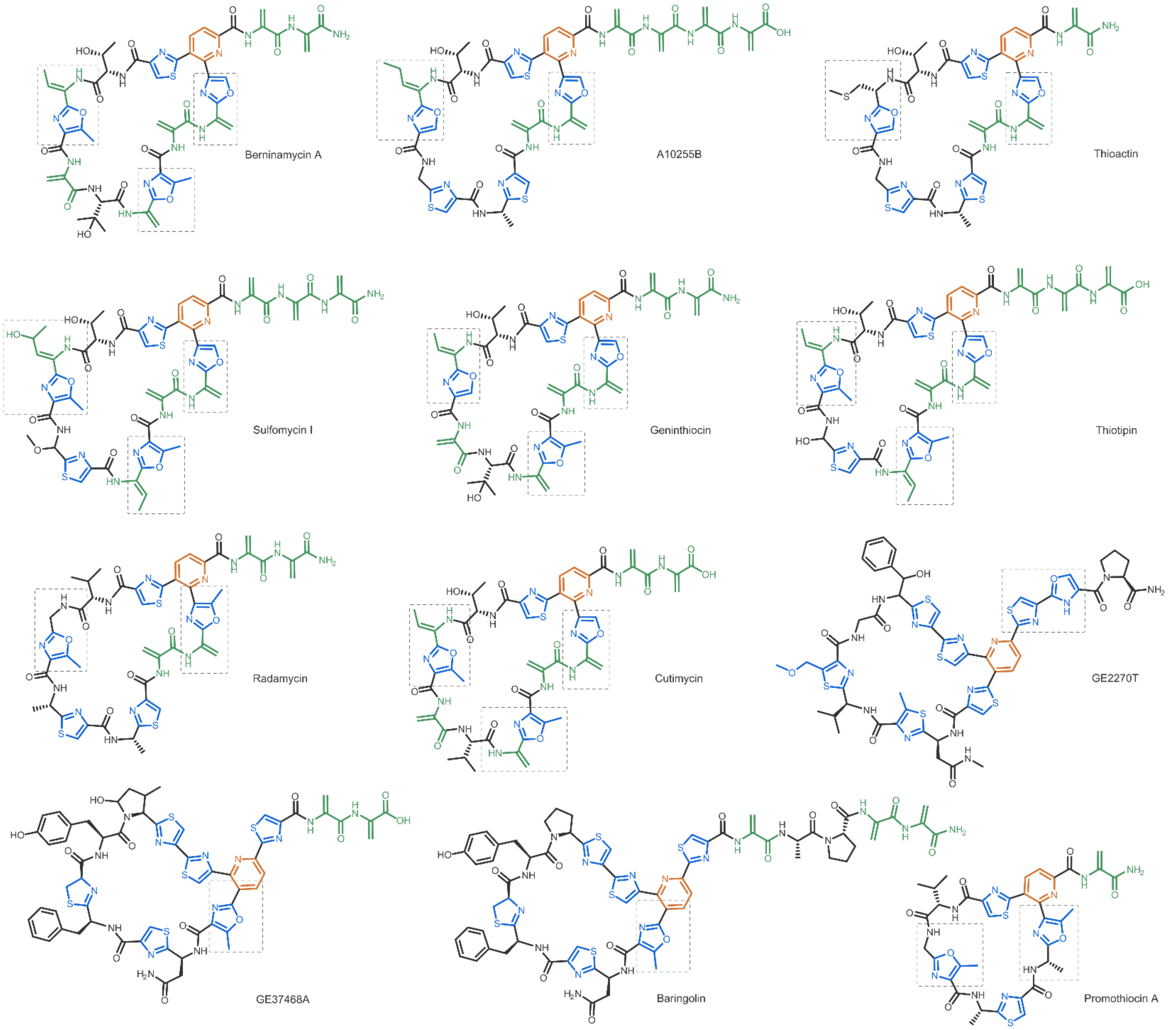

**Figure S46**. Oxz residues in thiopeptides are often flanked by a Dha in position –1: a survey of structurally characterized thiopeptides containing an Oxz residue. A number of redundant structures (e.g. berninamycin B and C, etc.) are omitted for clarity. Berninamycin-type thiopeptides often have 2 or 3 Oxz, usually flanked by a Dha. Promothiocin A notably stands out as the only thiopeptide containing 2 Oxz neither of which is conjugated to a Dha. Structures are taken from ref. [12–18]

#### 3.7. Optimized molecular geometries for Oxz/Thz tripeptides

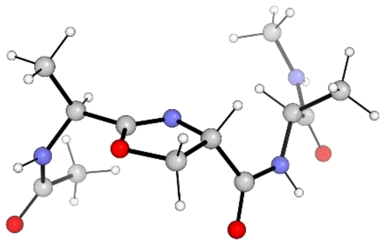

**Figure S47**. Optimized molecular structure of Ac-Ala-Oxn-Ala-NHMe. Calculated as described in section 2.6 (DFT at the B3LYP level of theory using 6-311+G(d) basis set). Molecular coordinates are below.

| | | | | | | | |
|---|---|---|---|---|---|---|---|
| HETATM | 1 | C | 0 | 1.636 | 0.588 | 0.772 | C |
| HETATM | 2 | O | 0 | 2.081 | 1.855 | 0.602 | O |
| HETATM | 3 | N | 0 | 0.384 | 0.355 | 0.653 | N |
| HETATM | 4 | C | 0 | 2.689 | -0.447 | 1.111 | C |
| HETATM | 5 | H | 0 | 2.178 | -1.409 | 1.145 | H |
| HETATM | 6 | C | 0 | -0.256 | 1.651 | 0.324 | C |
| HETATM | 7 | H | 0 | -0.974 | 1.902 | 1.110 | H |
| HETATM | 8 | C | 0 | 0.909 | 2.665 | 0.285 | C |
| HETATM | 9 | H | 0 | 0.830 | 3.445 | 1.045 | H |
| HETATM | 10 | H | 0 | 1.055 | 3.101 | -0.703 | H |
| HETATM | 11 | C | 0 | 3.329 | -0.169 | 2.479 | C |
| HETATM | 12 | H | 0 | 2.558 | -0.116 | 3.255 | H |
| HETATM | 13 | H | 0 | 4.023 | -0.976 | 2.729 | H |
| HETATM | 14 | H | 0 | 3.877 | 0.779 | 2.474 | H |
| HETATM | 15 | C | 0 | 3.769 | -1.308 | -1.024 | C |
| HETATM | 16 | O | 0 | 4.689 | -1.188 | -1.846 | O |
| HETATM | 17 | C | 0 | -0.999 | 1.524 | -1.022 | C |
| HETATM | 18 | O | 0 | -0.524 | 1.993 | -2.065 | O |
| HETATM | 19 | H | 0 | 4.453 | 0.200 | 0.117 | H |
| HETATM | 20 | N | 0 | 3.722 | -0.504 | 0.076 | N |
| HETATM | 21 | H | 0 | -2.631 | 0.746 | -1.919 | H |
| HETATM | 22 | C | 0 | -2.869 | 0.230 | 0.105 | C |
| HETATM | 23 | H | 0 | -2.129 | -0.071 | 0.847 | H |
| HETATM | 24 | N | 0 | -2.177 | 0.861 | -1.016 | N |
| HETATM | 25 | C | 0 | -3.911 | 1.174 | 0.742 | C |
| HETATM | 26 | H | 0 | -3.419 | 2.073 | 1.127 | H |
| HETATM | 27 | H | 0 | -4.652 | 1.478 | -0.006 | H |
| HETATM | 28 | H | 0 | -4.433 | 0.687 | 1.571 | H |
| HETATM | 29 | C | 0 | -3.582 | -1.015 | -0.452 | C |
| HETATM | 30 | O | 0 | -3.993 | -1.024 | -1.623 | O |
| HETATM | 31 | C | 0 | 2.685 | -2.356 | -1.187 | C |
| HETATM | 32 | H | 0 | 2.739 | -3.101 | -0.384 | H |
| HETATM | 33 | H | 0 | 1.685 | -1.912 | -1.166 | H |
| HETATM | 34 | H | 0 | 2.834 | -2.860 | -2.143 | H |
| HETATM | 35 | C | 0 | -3.278 | -2.212 | 1.754 | C |
| HETATM | 36 | H | 0 | -2.186 | -2.282 | 1.775 | H |
| HETATM | 37 | H | 0 | -3.600 | -1.392 | 2.402 | H |
| HETATM | 38 | H | 0 | -3.692 | -3.140 | 2.151 | H |
| HETATM | 39 | N | 0 | -3.783 | -2.050 | 0.390 | N |
| HETATM | 40 | H | 0 | -4.290 | -2.829 | -0.016 | H |

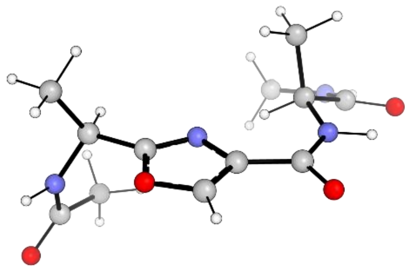

**Figure S48**. Optimized molecular structure of Ac-Ala-Oxz-Ala-NHMe. Calculated as described in section 2.6 (DFT at the B3LYP level of theory using 6-311+G(d) basis set). Molecular coordinates are below.

| | | | | | | | |
|---|---|---|---|---|---|---|---|
| HETATM | 1 | C | 0 | -1.656 | -0.787 | 0.435 | C |
| HETATM | 2 | O | 0 | -2.048 | -2.003 | -0.047 | O |
| HETATM | 3 | N | 0 | -0.382 | -0.574 | 0.313 | N |
| HETATM | 4 | C | 0 | -2.702 | 0.118 | 1.032 | C |
| HETATM | 5 | H | 0 | -2.181 | 1.037 | 1.305 | H |
| HETATM | 6 | C | 0 | 0.124 | -1.726 | -0.295 | C |
| HETATM | 7 | C | 0 | -3.330 | -0.491 | 2.296 | C |
| HETATM | 8 | H | 0 | -2.552 | -0.725 | 3.029 | H |
| HETATM | 9 | H | 0 | -4.028 | 0.225 | 2.740 | H |
| HETATM | 10 | H | 0 | -3.871 | -1.414 | 2.063 | H |
| HETATM | 11 | C | 0 | -3.764 | 1.450 | -0.857 | C |
| HETATM | 12 | O | 0 | -4.711 | 1.562 | -1.647 | O |
| HETATM | 13 | C | 0 | 1.533 | -2.008 | -0.674 | C |
| HETATM | 14 | O | 0 | 1.786 | -3.078 | -1.261 | O |
| HETATM | 15 | H | 0 | -4.543 | -0.192 | 0.009 | H |
| HETATM | 16 | N | 0 | -3.754 | 0.445 | 0.066 | N |
| HETATM | 17 | H | 0 | 3.445 | -1.423 | -0.701 | H |
| HETATM | 18 | C | 0 | 2.519 | 0.141 | 0.335 | C |
| HETATM | 19 | H | 0 | 1.590 | 0.669 | 0.130 | H |
| HETATM | 20 | N | 0 | 2.521 | -1.124 | -0.399 | N |
| HETATM | 21 | C | 0 | 2.658 | -0.080 | 1.856 | C |
| HETATM | 22 | H | 0 | 1.802 | -0.654 | 2.223 | H |
| HETATM | 23 | H | 0 | 3.577 | -0.634 | 2.078 | H |
| HETATM | 24 | H | 0 | 2.688 | 0.873 | 2.394 | H |
| HETATM | 25 | C | 0 | 3.723 | 0.945 | -0.182 | C |
| HETATM | 26 | O | 0 | 4.735 | 0.359 | -0.602 | O |
| HETATM | 27 | C | 0 | -2.600 | 2.421 | -0.859 | C |
| HETATM | 28 | H | 0 | -2.572 | 2.997 | 0.073 | H |
| HETATM | 29 | H | 0 | -1.641 | 1.904 | -0.961 | H |
| HETATM | 30 | H | 0 | -2.727 | 3.111 | -1.695 | H |
| HETATM | 31 | C | 0 | 2.498 | 3.111 | 0.260 | C |
| HETATM | 32 | H | 0 | 1.671 | 2.991 | -0.447 | H |
| HETATM | 33 | H | 0 | 2.146 | 2.878 | 1.269 | H |
| HETATM | 34 | H | 0 | 2.816 | 4.154 | 0.245 | H |
| HETATM | 35 | N | 0 | 3.651 | 2.290 | -0.112 | N |
| HETATM | 36 | H | 0 | 4.484 | 2.772 | -0.431 | H |
| HETATM | 37 | C | 0 | -0.910 | -2.589 | -0.511 | C |
| HETATM | 38 | H | 0 | -1.000 | -3.570 | -0.947 | H |

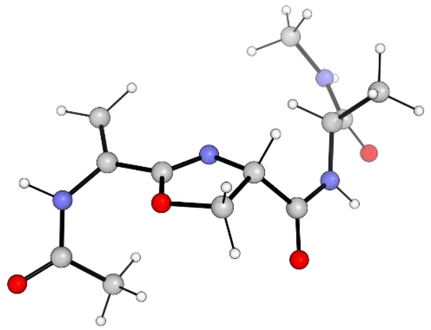

**Figure S49**. Optimized molecular structure of Ac-Dha-Oxn-Ala-NHMe. Calculated as described in section 2.6 (DFT at the B3LYP level of theory using 6-311+G(d) basis set). Molecular coordinates are below.

| | | | | | | | |
|---|---|---|---|---|---|---|---|
| HETATM | 1 | C | 0 | 1.588 | 0.222 | -0.721 | C |
| HETATM | 2 | O | 0 | 2.190 | -0.901 | -1.188 | O |
| HETATM | 3 | N | 0 | 0.315 | 0.218 | -0.571 | N |
| HETATM | 4 | C | 0 | -0.156 | -1.116 | -1.000 | C |
| HETATM | 5 | H | 0 | -0.829 | -0.998 | -1.855 | H |
| HETATM | 6 | C | 0 | 1.128 | -1.878 | -1.400 | C |
| HETATM | 7 | H | 0 | 1.150 | -2.166 | -2.451 | H |
| HETATM | 8 | H | 0 | 1.324 | -2.738 | -0.759 | H |
| HETATM | 9 | C | 0 | 4.402 | 0.313 | 0.776 | C |
| HETATM | 10 | O | 0 | 5.631 | 0.214 | 0.858 | O |
| HETATM | 11 | C | 0 | -0.927 | -1.787 | 0.158 | C |
| HETATM | 12 | O | 0 | -0.423 | -2.704 | 0.818 | O |
| HETATM | 13 | H | 0 | 4.529 | 1.652 | -0.721 | H |
| HETATM | 14 | N | 0 | 3.850 | 1.111 | -0.196 | N |
| HETATM | 15 | H | 0 | -2.639 | -1.741 | 1.223 | H |
| HETATM | 16 | C | 0 | -2.907 | -0.243 | -0.228 | C |
| HETATM | 17 | H | 0 | -2.192 | 0.452 | -0.670 | H |
| HETATM | 18 | N | 0 | -2.166 | -1.319 | 0.428 | N |
| HETATM | 19 | C | 0 | -3.866 | -0.783 | -1.309 | C |
| HETATM | 20 | H | 0 | -3.301 | -1.310 | -2.084 | H |
| HETATM | 21 | H | 0 | -4.581 | -1.483 | -0.864 | H |
| HETATM | 22 | H | 0 | -4.424 | 0.030 | -1.784 | H |
| HETATM | 23 | C | 0 | -3.719 | 0.473 | 0.866 | C |
| HETATM | 24 | O | 0 | -4.139 | -0.165 | 1.845 | O |
| HETATM | 25 | C | 0 | 3.474 | -0.415 | 1.723 | C |
| HETATM | 26 | H | 0 | 2.569 | 0.154 | 1.950 | H |
| HETATM | 27 | H | 0 | 3.175 | -1.373 | 1.282 | H |
| HETATM | 28 | H | 0 | 4.017 | -0.619 | 2.648 | H |
| HETATM | 29 | C | 0 | -3.499 | 2.670 | -0.364 | C |
| HETATM | 30 | H | 0 | -2.413 | 2.801 | -0.304 | H |
| HETATM | 31 | H | 0 | -3.764 | 2.303 | -1.360 | H |
| HETATM | 32 | H | 0 | -3.972 | 3.643 | -0.225 | H |
| HETATM | 33 | N | 0 | -3.995 | 1.781 | 0.688 | N |
| HETATM | 34 | H | 0 | -4.567 | 2.193 | 1.417 | H |
| HETATM | 35 | C | 0 | 2.480 | 1.378 | -0.457 | C |
| HETATM | 36 | C | 0 | 2.026 | 2.634 | -0.586 | C |
| HETATM | 37 | H | 0 | 2.673 | 3.486 | -0.405 | H |
| HETATM | 38 | H | 0 | 1.006 | 2.821 | -0.901 | H |

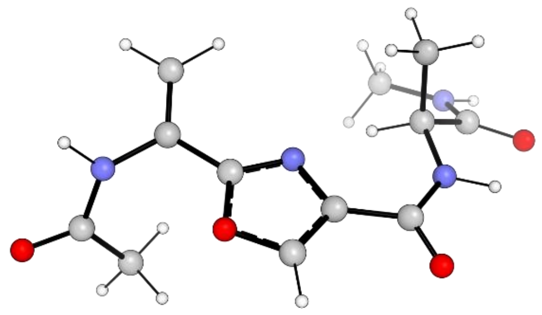

**Figure S50**. Optimized molecular structure of Ac-Dha-Oxz-Ala-NHMe. Calculated as described in section 2.6 (DFT at the B3LYP level of theory using 6-311+G(d) basis set). Molecular coordinates are below.

| | | | | | | | |
|---|---|---|---|---|---|---|---|
| HETATM | 1 | C | 0 | 1.619 | 0.238 | 0.552 | C |
| HETATM | 2 | O | 0 | 2.167 | 1.459 | 0.267 | O |
| HETATM | 3 | N | 0 | 0.320 | 0.230 | 0.472 | N |
| HETATM | 4 | C | 0 | -0.031 | 1.530 | 0.124 | C |
| HETATM | 5 | C | 0 | 4.471 | -0.587 | -0.641 | C |
| HETATM | 6 | O | 0 | 5.699 | -0.501 | -0.740 | O |
| HETATM | 7 | C | 0 | -1.398 | 2.073 | -0.095 | C |
| HETATM | 8 | O | 0 | -1.517 | 3.285 | -0.356 | O |
| HETATM | 9 | H | 0 | 4.562 | -0.829 | 1.359 | H |
| HETATM | 10 | N | 0 | 3.896 | -0.714 | 0.601 | N |
| HETATM | 11 | H | 0 | -3.369 | 1.751 | -0.190 | H |
| HETATM | 12 | C | 0 | -2.632 | -0.153 | 0.273 | C |
| HETATM | 13 | H | 0 | -1.759 | -0.682 | -0.107 | H |
| HETATM | 14 | N | 0 | -2.487 | 1.273 | -0.020 | N |
| HETATM | 15 | C | 0 | -2.773 | -0.414 | 1.786 | C |
| HETATM | 16 | H | 0 | -1.866 | -0.083 | 2.300 | H |
| HETATM | 17 | H | 0 | -3.630 | 0.136 | 2.189 | H |
| HETATM | 18 | H | 0 | -2.916 | -1.480 | 1.993 | H |
| HETATM | 19 | C | 0 | -3.904 | -0.620 | -0.453 | C |
| HETATM | 20 | O | 0 | -4.845 | 0.170 | -0.638 | O |
| HETATM | 21 | C | 0 | 3.564 | -0.561 | -1.852 | C |
| HETATM | 22 | H | 0 | 2.678 | -1.190 | -1.731 | H |
| HETATM | 23 | H | 0 | 3.233 | 0.466 | -2.044 | H |
| HETATM | 24 | H | 0 | 4.137 | -0.900 | -2.717 | H |
| HETATM | 25 | C | 0 | -2.911 | -2.922 | -0.765 | C |
| HETATM | 26 | H | 0 | -2.071 | -2.665 | -1.419 | H |
| HETATM | 27 | H | 0 | -2.544 | -3.059 | 0.256 | H |
| HETATM | 28 | H | 0 | -3.334 | -3.869 | -1.103 | H |
| HETATM | 29 | N | 0 | -3.971 | -1.914 | -0.826 | N |
| HETATM | 30 | H | 0 | -4.844 | -2.181 | -1.268 | H |
| HETATM | 31 | C | 0 | 2.520 | -0.858 | 0.929 | C |
| HETATM | 32 | C | 0 | 2.076 | -1.906 | 1.642 | C |
| HETATM | 33 | H | 0 | 2.742 | -2.721 | 1.906 | H |
| HETATM | 34 | H | 0 | 1.049 | -1.948 | 1.987 | H |
| HETATM | 35 | C | 0 | 1.110 | 2.271 | -0.001 | C |
| HETATM | 36 | H | 0 | 1.325 | 3.295 | -0.259 | H |

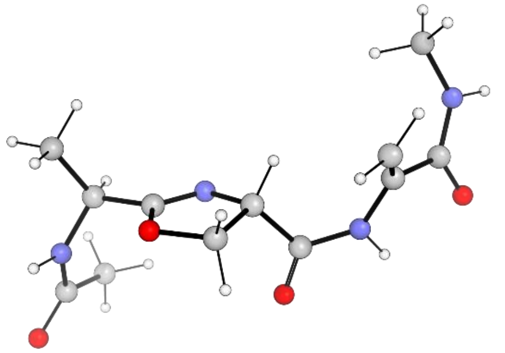

**Figure S51**. Optimized molecular structure of Ac-Ala-Oxn-Dha-NHMe. Calculated as described in section 2.6 (DFT at the B3LYP level of theory using 6-311+G(d) basis set). Molecular coordinates are below.

| | | | | | | | |
|---|---|---|---|---|---|---|---|
| HETATM | 1 | C | 0 | -1.788 | 0.829 | 0.056 | C |
| HETATM | 2 | O | 0 | -1.821 | 1.132 | 1.379 | O |
| HETATM | 3 | N | 0 | -0.690 | 0.414 | -0.449 | N |
| HETATM | 4 | C | 0 | -3.077 | 1.046 | -0.707 | C |
| HETATM | 5 | H | 0 | -2.897 | 0.694 | -1.724 | H |
| HETATM | 6 | C | 0 | 0.308 | 0.419 | 0.643 | C |
| HETATM | 7 | H | 0 | 1.061 | 1.173 | 0.410 | H |
| HETATM | 8 | C | 0 | -0.510 | 0.792 | 1.911 | C |
| HETATM | 9 | H | 0 | -0.125 | 1.664 | 2.440 | H |
| HETATM | 10 | H | 0 | -0.638 | -0.046 | 2.600 | H |
| HETATM | 11 | C | 0 | -3.457 | 2.534 | -0.752 | C |
| HETATM | 12 | H | 0 | -2.639 | 3.121 | -1.179 | H |
| HETATM | 13 | H | 0 | -4.346 | 2.665 | -1.376 | H |
| HETATM | 14 | H | 0 | -3.670 | 2.918 | 0.251 | H |
| HETATM | 15 | C | 0 | -4.563 | -1.000 | -0.454 | C |
| HETATM | 16 | O | 0 | -5.477 | -1.552 | 0.177 | O |
| HETATM | 17 | C | 0 | 0.942 | -0.973 | 0.745 | C |
| HETATM | 18 | O | 0 | 0.229 | -1.967 | 0.901 | O |
| HETATM | 19 | H | 0 | -4.647 | 0.661 | 0.675 | H |
| HETATM | 20 | N | 0 | -4.172 | 0.264 | -0.130 | N |
| HETATM | 21 | H | 0 | 2.606 | -2.077 | 0.496 | H |
| HETATM | 22 | N | 0 | 2.295 | -1.113 | 0.607 | N |
| HETATM | 23 | C | 0 | 4.429 | -0.662 | -0.418 | C |
| HETATM | 24 | O | 0 | 4.672 | -1.882 | -0.420 | O |
| HETATM | 25 | C | 0 | -3.869 | -1.685 | -1.615 | C |
| HETATM | 26 | H | 0 | -4.070 | -1.159 | -2.556 | H |
| HETATM | 27 | H | 0 | -2.784 | -1.718 | -1.475 | H |
| HETATM | 28 | H | 0 | -4.252 | -2.704 | -1.694 | H |
| HETATM | 29 | C | 0 | 4.865 | 1.614 | -1.479 | C |
| HETATM | 30 | H | 0 | 3.810 | 1.849 | -1.334 | H |
| HETATM | 31 | H | 0 | 5.472 | 2.274 | -0.849 | H |
| HETATM | 32 | H | 0 | 5.122 | 1.797 | -2.525 | H |
| HETATM | 33 | N | 0 | 5.120 | 0.204 | -1.186 | N |
| HETATM | 34 | H | 0 | 5.874 | -0.237 | -1.705 | H |
| HETATM | 35 | C | 0 | 3.339 | -0.164 | 0.503 | C |
| HETATM | 36 | C | 0 | 3.449 | 0.937 | 1.261 | C |
| HETATM | 37 | H | 0 | 2.737 | 1.169 | 2.046 | H |
| HETATM | 38 | H | 0 | 4.275 | 1.625 | 1.132 | H |

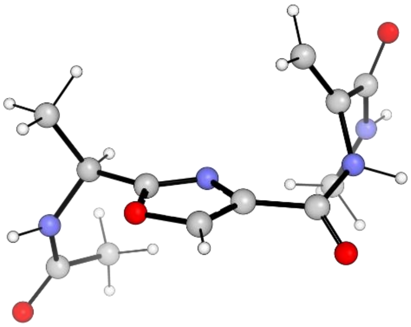

**Figure S52**. Optimized molecular structure of Ac-Ala-Oxz-Dha-NHMe. Calculated as described in section 2.6 (DFT at the B3LYP level of theory using 6-311+G(d) basis set). Molecular coordinates are below.

| | | | | | | | |
|---|---|---|---|---|---|---|---|
| HETATM | 1 | C | 0 | -1.413 | -0.687 | 0.452 | C |
| HETATM | 2 | O | 0 | -1.724 | -1.976 | 0.116 | O |
| HETATM | 3 | N | 0 | -0.212 | -0.341 | 0.107 | N |
| HETATM | 4 | C | 0 | -2.458 | 0.138 | 1.159 | C |
| HETATM | 5 | H | 0 | -2.023 | 1.132 | 1.278 | H |
| HETATM | 6 | C | 0 | 0.324 | -1.472 | -0.510 | C |
| HETATM | 7 | C | 0 | -2.786 | -0.429 | 2.549 | C |
| HETATM | 8 | H | 0 | -1.875 | -0.505 | 3.151 | H |
| HETATM | 9 | H | 0 | -3.491 | 0.235 | 3.058 | H |
| HETATM | 10 | H | 0 | -3.232 | -1.427 | 2.474 | H |
| HETATM | 11 | C | 0 | -3.980 | 1.175 | -0.594 | C |
| HETATM | 12 | O | 0 | -5.047 | 1.104 | -1.219 | O |
| HETATM | 13 | C | 0 | 1.658 | -1.554 | -1.154 | C |
| HETATM | 14 | O | 0 | 1.833 | -2.300 | -2.125 | O |
| HETATM | 15 | H | 0 | -4.378 | -0.480 | 0.480 | H |
| HETATM | 16 | N | 0 | -3.685 | 0.254 | 0.368 | N |
| HETATM | 17 | H | 0 | 3.481 | -0.769 | -1.328 | H |
| HETATM | 18 | N | 0 | 2.675 | -0.746 | -0.708 | N |
| HETATM | 19 | C | 0 | 3.645 | 1.140 | 0.595 | C |
| HETATM | 20 | O | 0 | 4.516 | 1.339 | 1.458 | O |
| HETATM | 21 | C | 0 | -2.979 | 2.285 | -0.843 | C |
| HETATM | 22 | H | 0 | -2.890 | 2.933 | 0.037 | H |
| HETATM | 23 | H | 0 | -1.983 | 1.891 | -1.070 | H |
| HETATM | 24 | H | 0 | -3.329 | 2.884 | -1.685 | H |
| HETATM | 25 | C | 0 | 2.217 | 2.146 | -1.260 | C |
| HETATM | 26 | H | 0 | 2.444 | 1.718 | -2.242 | H |
| HETATM | 27 | H | 0 | 1.354 | 1.634 | -0.834 | H |
| HETATM | 28 | H | 0 | 1.965 | 3.202 | -1.388 | H |
| HETATM | 29 | N | 0 | 3.366 | 2.065 | -0.355 | N |
| HETATM | 30 | H | 0 | 3.957 | 2.887 | -0.295 | H |
| HETATM | 31 | C | 0 | 2.894 | -0.173 | 0.571 | C |
| HETATM | 32 | C | 0 | 2.536 | -0.753 | 1.724 | C |
| HETATM | 33 | H | 0 | 2.045 | -1.719 | 1.758 | H |
| HETATM | 34 | H | 0 | 2.713 | -0.239 | 2.662 | H |
| HETATM | 35 | C | 0 | -0.614 | -2.460 | -0.508 | C |
| HETATM | 36 | H | 0 | -0.650 | -3.475 | -0.868 | H |

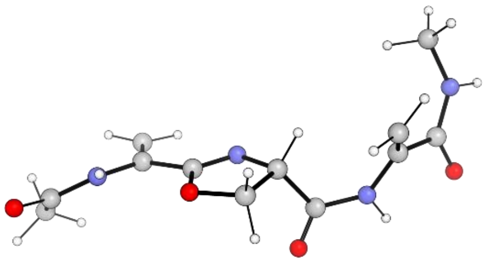

**Figure S53**. Optimized molecular structure of Ac-Dha-Oxn-Dha-NHMe. Calculated as described in section 2.6 (DFT at the B3LYP level of theory using 6-311+G(d) basis set). Molecular coordinates are below.

| | | | | | | | |
|---|---|---|---|---|---|---|---|
| HETATM | 1 | C | 0 | 1.685 | 0.118 | -0.267 | C |
| HETATM | 2 | O | 0 | 1.815 | -1.074 | -0.909 | O |
| HETATM | 3 | N | 0 | 0.510 | 0.521 | 0.036 | N |
| HETATM | 4 | C | 0 | -0.432 | -0.517 | -0.422 | C |
| HETATM | 5 | H | 0 | -1.108 | -0.070 | -1.152 | H |
| HETATM | 6 | C | 0 | 0.471 | -1.608 | -1.064 | C |
| HETATM | 7 | H | 0 | 0.286 | -1.749 | -2.130 | H |
| HETATM | 8 | H | 0 | 0.435 | -2.567 | -0.543 | H |
| HETATM | 9 | C | 0 | 5.395 | 0.148 | -0.110 | C |
| HETATM | 10 | O | 0 | 6.224 | -0.458 | -0.798 | O |
| HETATM | 11 | C | 0 | -1.208 | -1.043 | 0.794 | C |
| HETATM | 12 | O | 0 | -0.598 | -1.523 | 1.753 | O |
| HETATM | 13 | H | 0 | 3.930 | -0.358 | -1.376 | H |
| HETATM | 14 | N | 0 | 4.094 | 0.220 | -0.557 | N |
| HETATM | 15 | H | 0 | -2.978 | -1.113 | 1.747 | H |
| HETATM | 16 | N | 0 | -2.569 | -0.924 | 0.834 | N |
| HETATM | 17 | C | 0 | -4.646 | 0.257 | 0.522 | C |
| HETATM | 18 | O | 0 | -5.009 | -0.112 | 1.654 | O |
| HETATM | 19 | C | 0 | 5.774 | 0.782 | 1.207 | C |
| HETATM | 20 | H | 0 | 6.040 | 1.835 | 1.058 | H |
| HETATM | 21 | H | 0 | 4.970 | 0.735 | 1.946 | H |
| HETATM | 22 | H | 0 | 6.657 | 0.262 | 1.585 | H |
| HETATM | 23 | C | 0 | -4.858 | 1.984 | -1.341 | C |
| HETATM | 24 | H | 0 | -3.785 | 1.885 | -1.512 | H |
| HETATM | 25 | H | 0 | -5.399 | 1.610 | -2.217 | H |
| HETATM | 26 | H | 0 | -5.093 | 3.043 | -1.212 | H |
| HETATM | 27 | N | 0 | -5.249 | 1.276 | -0.122 | N |
| HETATM | 28 | H | 0 | -6.043 | 1.654 | 0.385 | H |
| HETATM | 29 | C | 0 | -3.512 | -0.498 | -0.132 | C |
| HETATM | 30 | C | 0 | -3.512 | -0.875 | -1.419 | C |
| HETATM | 31 | H | 0 | -2.781 | -1.572 | -1.816 | H |
| HETATM | 32 | H | 0 | -4.265 | -0.514 | -2.107 | H |
| HETATM | 33 | C | 0 | 2.954 | 0.848 | -0.011 | C |
| HETATM | 34 | C | 0 | 2.944 | 2.008 | 0.664 | C |
| HETATM | 35 | H | 0 | 3.835 | 2.603 | 0.815 | H |
| HETATM | 36 | H | 0 | 2.012 | 2.374 | 1.079 | H |

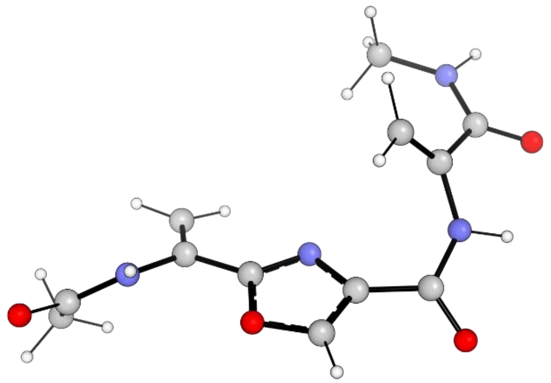

**Figure S54**. Optimized molecular structure of Ac-Dha-Oxz-Dha-NHMe. Calculated as described in section 2.6 (DFT at the B3LYP level of theory using 6-311+G(d) basis set). Molecular coordinates are below.

| | | | | | | | |
|---|---|---|---|---|---|---|---|
| HETATM | 1 | C | 0 | 1.383 | 0.399 | -0.023 | C |
| HETATM | 2 | O | 0 | 1.768 | 1.636 | 0.425 | O |
| HETATM | 3 | N | 0 | 0.117 | 0.331 | -0.317 | N |
| HETATM | 4 | C | 0 | -0.375 | 1.606 | -0.072 | C |
| HETATM | 5 | C | 0 | 4.924 | -0.525 | 0.135 | C |
| HETATM | 6 | O | 0 | 5.841 | -0.215 | 0.904 | O |
| HETATM | 7 | C | 0 | -1.762 | 2.058 | -0.351 | C |
| HETATM | 8 | O | 0 | -1.963 | 3.221 | -0.725 | O |
| HETATM | 9 | H | 0 | 3.554 | 0.137 | 1.440 | H |
| HETATM | 10 | N | 0 | 3.623 | -0.342 | 0.546 | N |
| HETATM | 11 | H | 0 | -3.672 | 1.511 | -0.662 | H |
| HETATM | 12 | N | 0 | -2.800 | 1.172 | -0.258 | N |
| HETATM | 13 | C | 0 | -4.094 | -0.842 | -0.098 | C |
| HETATM | 14 | O | 0 | -5.067 | -0.227 | -0.571 | O |
| HETATM | 15 | C | 0 | 5.202 | -1.075 | -1.244 | C |
| HETATM | 16 | H | 0 | 5.282 | -2.168 | -1.202 | H |
| HETATM | 17 | H | 0 | 4.427 | -0.819 | -1.971 | H |
| HETATM | 18 | H | 0 | 6.166 | -0.681 | -1.572 | H |
| HETATM | 19 | C | 0 | -2.954 | -3.099 | 0.239 | C |
| HETATM | 20 | H | 0 | -1.998 | -2.601 | 0.079 | H |
| HETATM | 21 | H | 0 | -3.001 | -3.466 | 1.271 | H |
| HETATM | 22 | H | 0 | -3.021 | -3.954 | -0.438 | H |
| HETATM | 23 | N | 0 | -4.060 | -2.190 | -0.062 | N |
| HETATM | 24 | H | 0 | -4.916 | -2.618 | -0.400 | H |
| HETATM | 25 | C | 0 | -2.951 | -0.033 | 0.466 | C |
| HETATM | 26 | C | 0 | -2.311 | -0.335 | 1.604 | C |
| HETATM | 27 | H | 0 | -1.600 | 0.343 | 2.062 | H |
| HETATM | 28 | H | 0 | -2.491 | -1.274 | 2.113 | H |
| HETATM | 29 | C | 0 | 2.408 | -0.650 | -0.108 | C |
| HETATM | 30 | C | 0 | 2.163 | -1.801 | -0.757 | C |
| HETATM | 31 | H | 0 | 2.881 | -2.612 | -0.772 | H |
| HETATM | 32 | H | 0 | 1.230 | -1.935 | -1.292 | H |
| HETATM | 33 | C | 0 | 0.641 | 2.397 | 0.378 | C |
| HETATM | 34 | H | 0 | 0.724 | 3.425 | 0.693 | H |

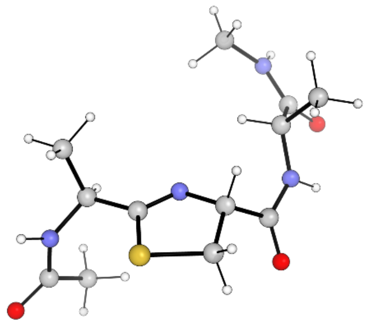

**Figure S55**. Optimized molecular structure of Ac-Ala-Thn-Ala-NHMe. Calculated as described in section 2.6 (DFT at the B3LYP level of theory using 6-311+G(d) basis set). Molecular coordinates are below.

| | | | | | | | |
|---|---|---|---|---|---|---|---|
| HETATM | 1 | C | 0 | 1.491 | -0.033 | -0.581 | C |
| HETATM | 2 | N | 0 | 0.256 | -0.087 | -0.268 | N |
| HETATM | 3 | C | 0 | 2.287 | 1.265 | -0.478 | C |
| HETATM | 4 | H | 0 | 1.825 | 1.844 | 0.327 | H |
| HETATM | 5 | C | 0 | -0.357 | -1.389 | -0.575 | C |
| HETATM | 6 | H | 0 | -0.965 | -1.251 | -1.477 | H |
| HETATM | 7 | C | 0 | 0.732 | -2.455 | -0.846 | C |
| HETATM | 8 | H | 0 | 0.476 | -3.094 | -1.693 | H |
| HETATM | 9 | H | 0 | 0.906 | -3.071 | 0.037 | H |
| HETATM | 10 | C | 0 | 2.173 | 2.071 | -1.782 | C |
| HETATM | 11 | H | 0 | 1.122 | 2.260 | -2.014 | H |
| HETATM | 12 | H | 0 | 2.687 | 3.031 | -1.668 | H |
| HETATM | 13 | H | 0 | 2.620 | 1.530 | -2.624 | H |
| HETATM | 14 | C | 0 | 4.249 | 0.804 | 1.061 | C |
| HETATM | 15 | O | 0 | 5.470 | 0.619 | 1.171 | O |
| HETATM | 16 | C | 0 | -1.284 | -1.794 | 0.594 | C |
| HETATM | 17 | O | 0 | -0.936 | -2.645 | 1.423 | O |
| HETATM | 18 | H | 0 | 4.356 | 1.053 | -0.932 | H |
| HETATM | 19 | N | 0 | 3.696 | 1.049 | -0.161 | N |
| HETATM | 20 | H | 0 | -3.048 | -1.409 | 1.491 | H |
| HETATM | 21 | C | 0 | -3.044 | -0.130 | -0.175 | C |
| HETATM | 22 | H | 0 | -2.229 | 0.433 | -0.631 | H |
| HETATM | 23 | N | 0 | -2.476 | -1.164 | 0.686 | N |
| HETATM | 24 | C | 0 | -3.957 | -0.728 | -1.266 | C |
| HETATM | 25 | H | 0 | -3.383 | -1.403 | -1.909 | H |
| HETATM | 26 | H | 0 | -4.773 | -1.296 | -0.806 | H |
| HETATM | 27 | H | 0 | -4.391 | 0.057 | -1.894 | H |
| HETATM | 28 | C | 0 | -3.872 | 0.801 | 0.730 | C |
| HETATM | 29 | O | 0 | -4.425 | 0.350 | 1.745 | O |
| HETATM | 30 | C | 0 | 3.325 | 0.787 | 2.261 | C |
| HETATM | 31 | H | 0 | 2.893 | 1.780 | 2.433 | H |
| HETATM | 32 | H | 0 | 2.496 | 0.084 | 2.126 | H |
| HETATM | 33 | H | 0 | 3.901 | 0.497 | 3.141 | H |
| HETATM | 34 | C | 0 | -3.356 | 2.767 | -0.773 | C |
| HETATM | 35 | H | 0 | -2.268 | 2.791 | -0.651 | H |
| HETATM | 36 | H | 0 | -3.602 | 2.299 | -1.730 | H |
| HETATM | 37 | H | 0 | -3.721 | 3.795 | -0.794 | H |
| HETATM | 38 | N | 0 | -4.009 | 2.087 | 0.346 | N |
| HETATM | 39 | H | 0 | -4.600 | 2.645 | 0.952 | H |
| HETATM | 40 | S | 0 | 2.265 | -1.509 | -1.227 | S |

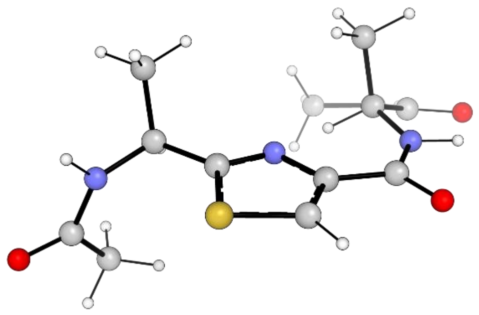

**Figure S56**. Optimized molecular structure of Ac-Ala-Thz-Ala-NHMe. Calculated as described in section 2.6 (DFT at the B3LYP level of theory using 6-311+G(d) basis set). Molecular coordinates are below.

| | | | | | | | |
|---|---|---|---|---|---|---|---|
| HETATM | 1 | C | 0 | 1.456 | 0.408 | 0.374 | C |
| HETATM | 2 | N | 0 | 0.161 | 0.443 | 0.255 | N |
| HETATM | 3 | C | 0 | 2.227 | -0.822 | 0.826 | C |
| HETATM | 4 | H | 0 | 1.787 | -1.679 | 0.310 | H |
| HETATM | 5 | C | 0 | -0.284 | 1.697 | -0.122 | C |
| HETATM | 6 | C | 0 | 2.077 | -1.043 | 2.341 | C |
| HETATM | 7 | H | 0 | 1.018 | -1.108 | 2.605 | H |
| HETATM | 8 | H | 0 | 2.572 | -1.976 | 2.630 | H |
| HETATM | 9 | H | 0 | 2.522 | -0.215 | 2.904 | H |
| HETATM | 10 | C | 0 | 4.254 | -1.153 | -0.669 | C |
| HETATM | 11 | O | 0 | 5.482 | -1.054 | -0.796 | O |
| HETATM | 12 | C | 0 | -1.718 | 2.048 | -0.354 | C |
| HETATM | 13 | O | 0 | -2.001 | 3.208 | -0.710 | O |
| HETATM | 14 | H | 0 | 4.279 | -0.397 | 1.196 | H |
| HETATM | 15 | N | 0 | 3.644 | -0.741 | 0.481 | N |
| HETATM | 16 | H | 0 | -3.636 | 1.497 | -0.385 | H |
| HETATM | 17 | C | 0 | -2.705 | -0.259 | 0.258 | C |
| HETATM | 18 | H | 0 | -1.803 | -0.750 | -0.102 | H |
| HETATM | 19 | N | 0 | -2.705 | 1.131 | -0.198 | N |
| HETATM | 20 | C | 0 | -2.763 | -0.357 | 1.797 | C |
| HETATM | 21 | H | 0 | -1.870 | 0.107 | 2.225 | H |
| HETATM | 22 | H | 0 | -3.650 | 0.160 | 2.178 | H |
| HETATM | 23 | H | 0 | -2.801 | -1.401 | 2.127 | H |
| HETATM | 24 | C | 0 | -3.958 | -0.914 | -0.347 | C |
| HETATM | 25 | O | 0 | -4.973 | -0.236 | -0.576 | O |
| HETATM | 26 | C | 0 | 3.380 | -1.737 | -1.761 | C |
| HETATM | 27 | H | 0 | 2.977 | -2.709 | -1.452 | H |
| HETATM | 28 | H | 0 | 2.534 | -1.086 | -2.003 | H |
| HETATM | 29 | H | 0 | 3.990 | -1.880 | -2.654 | H |
| HETATM | 30 | C | 0 | -2.775 | -3.145 | -0.444 | C |
| HETATM | 31 | H | 0 | -1.997 | -2.909 | -1.178 | H |
| HETATM | 32 | H | 0 | -2.343 | -3.117 | 0.560 | H |
| HETATM | 33 | H | 0 | -3.130 | -4.160 | -0.631 | H |
| HETATM | 34 | N | 0 | -3.922 | -2.245 | -0.563 | N |
| HETATM | 35 | H | 0 | -4.784 | -2.633 | -0.930 | H |
| HETATM | 36 | S | 0 | 2.251 | 1.937 | 0.024 | S |
| HETATM | 37 | C | 0 | 0.709 | 2.633 | -0.290 | C |
| HETATM | 38 | H | 0 | 0.599 | 3.665 | -0.589 | H |

**Table S4.** Calculated total Gibbs free energies after thermal correction for the tripeptides used in the reduction potential calculations; a. u. stands for atomic units (Hartrees). From these data, $E^{0'}$ values can be calculated as described in section 2.6.

| compound | phase | $\Delta G^0_{tot}$, a.u. |
|---|---|---|
| Ac-Ala-Oxn-Ala-NHMe | water | -989.047017 |
| Ac-Dha-Oxn-Ala-NHMe | water | -987.836932 |
| Ac-Ala-Oxn-Dha-NHMe | water | -987.831092 |
| Ac-Ala-Thn-Ala-NHMe | water | -1312.019420 |
| Ac-Dha-Oxn-Dha-NHMe | water | -986.624703 |
| Ac-Ala-Oxz-Ala-NHMe | water | -987.862694 |
| Ac-Dha-Oxz-Ala-NHMe | water | -986.655107 |
| Ac-Ala-Oxz-Dha-NHMe | water | -986.646305 |
| Ac-Ala-Thz-Ala-NHMe | water | -1310.836559 |
| Ac-Dha-Oxz-Dha-NHMe | water | -985.441029 |

#### 3.8. Electronic absorption spectra of LazF/FMN and phenosafranine

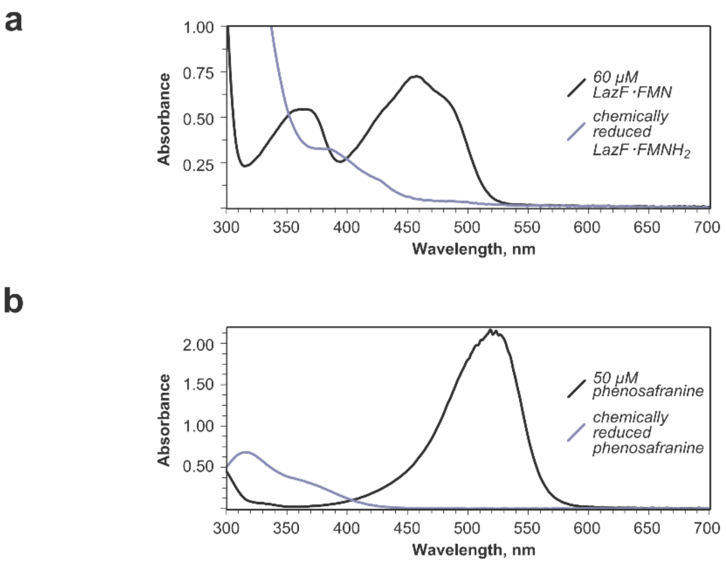

**Figure S57**. Electronic absorption spectra of LazF/FMN (panel a) and phenosafranine (panel b). Upon reduction, both compounds lose their absorption in the visible range, which enables quantification of reduced and oxidized forms for each component. At 540 nm, where oxidized phenosafranine absorbs, spectral interference of either oxidized or reduced LazF is minimal; at 457 nm, the absorption maximum of LazF/FMN ($\varepsilon^{457}$ = 11900 $M^{-1}cm^{-1}$), phenosafranine has $\varepsilon^{457}$ = 11300 $M^{-1}cm^{-1}$, and thus, when calculating the fraction of reduced FMN, the spectrum had to be corrected for phenosafranine absorption.
